# Analysis of gene expression changes after Latrunculin B-induced actin depolymerization

**DOI:** 10.64898/2026.10.14.759175

**Authors:** Giorgos Kargas, Michael J. Layden, Dimitrios Vavylonis

## Abstract

Links between the cytoskeleton and gene expression are increasingly appreciated. The cytoskeleton dynamically regulates its own expression and organization through transcriptional feedback loops that sense cellular concentrations and mechanical forces. Disruption of cytoskeletal dynamics can also influence cell fate. However, the breadth, temporal dynamics, and mechanistic basis of the cytoskeleton-transcription coupling have yet to be fully resolved. To characterize this response more broadly, we re-analyzed time-resolved bulk RNA-seq data that were previously used to demonstrate that treatment with actin depolymerization drug Latrunculin B (LatB) drives differentiation of human pancreatic progenitor cells in culture. Beyond the previously identified induction of cell differentiation, we find that LatB produced extensive transcriptional changes within the actin cytoskeleton, including downregulation of actin, myosin II heavy chains, myosin I, tropomyosins, α-actinins, filamins, talin, vinculin and plastin. The changes were not restricted to the actin system, but extended to microtubule and intermediate filament systems, with noticeable changes in the expression of KIF19, MAP1A, MAP1B, Type I and II keratins and Type IV neurofilaments. More broadly, the response extended to genes involved in transcription, vesicle trafficking, metabolism, translation, ion transport, protein homeostasis, and cell signaling. The breadth of this response suggested engagement of multiple transcriptional regulatory programs. Transcription factor activity inference using decoupleR identified MRTF/SRF as the most inactivated TF network, consistent with its known role in actin homeostasis, in addition to Hippo/YAP-TEAD, MYC, stress-response and other pathways. These results provide the broader landscape of cellular response to actin depolymerization, serving as a foundation for future studies to distinguish general mechanisms of cytoskeletal homeostasis from differentiation-associated responses, and to narrow the underlying regulatory mechanisms linking cytoskeletal state to gene expression.

## Introduction

Cells sense and respond to the physical properties of their environment through the cytoskeleton, which connects extracellular and subcellular mechanical cues to changes in gene expression (Iskratsch, Wolfenson, & Sheetz, 2014; Uhler & Shivashankar, 2017). This mechanical sensing allows cells to interpret signals like matrix stiffness and cell shape, and translate them into fate decisions. Two key transcriptional regulators in this process are YAP and TAZ, which respond to F-actin organization and cytoskeletal tension through both Hippo-dependent (LATS1/2-mediated phosphorylation) and Hippo-independent mechanisms (Dupont et al., 2011; Panciera, Azzolin, Cordenonsi, & Piccolo, 2017). When cells spread on stiff substrates and build robust actin networks, YAP/TAZ enter the nucleus and drive proliferation (Aragona et al., 2013). When tension drops, they stay in the cytoplasm, allowing differentiation to proceed (Panciera et al., 2017; Vogel & Sheetz, 2006).

Maintaining cytoskeletal organization presents an additional regulatory challenge because cytoskeletal components span a wide range of cellular concentrations, from nM levels up to ∼100 μM for highly abundant structural proteins such as actin. Proteins acting at filament ends typically function at relatively low concentrations, whereas proteins that bind monomers, along filament sides, crosslink filaments, or constitute the polymers themselves may be required at much higher concentrations. Despite these differences in abundance and function, their expression and activity must remain coordinated as cytoskeletal assembly and mechanical state change. This raises the question of whether the relative abundance of cytoskeletal components is also coordinated at the level of gene expression. In the typical steady state condition, a large fraction of actin exists in filamentous form (F-actin), while soluble actin is distributed among complexes with profilin, cofilin, thymosin-β4, and a smaller fraction remains as free monomeric actin (G-actin) (Skruber, Read, & Vitriol, 2018). A shift in the G-actin/F-actin balance therefore changes the availability of binding sites for these proteins, raising the question of whether their encoding genes are coordinately regulated at the gene expression level (Golla et al., 1997). Homeostatic regulation is also observed between cytoplasmic actin isoforms, with perturbations of ACTB accompanied by compensatory changes in the expression of other actin isoforms, including ACTG1 (Vedula et al., 2017). The actin-MRTF-SRF pathway provides a link between the actin cytoskeleton and transcription (Olson & Nordheim, 2010). G-actin binds and traps MRTF cofactors in the cytoplasm, limiting their nuclear accumulation (Miralles, Posern, Zaromytidou, & Treisman, 2003). When actin polymerizes into filaments and reduces the available G-actin pool, MRTF enters the nucleus and cooperates with SRF to activate genes encoding cytoskeletal proteins (Olson & Nordheim, 2010). This feedback provides a direct means by which changes in actin polymerization state can be translated into transcriptional responses in order to regulate cytoskeletal composition.

The coupling between cytoskeletal state, mechanotransduction, and cell fate is particularly evident in pancreatic development. Bipotent pancreatic progenitors choose between ductal and endocrine fates based partly on mechanical signals (Mamidi et al., 2018; Rosado-Olivieri, Anderson, Kenty, & Melton, 2019). When progenitors (Tian, Rump, Ebeid, Mamidi, & Semb, 2025) spread on fibronectin-rich matrix, integrin engagement triggers F-actin bundling and YAP activation, pushing cells toward a ductal fate, on laminin/collagen substrates, reduced spreading keeps YAP inactive and permits endocrine differentiation (Mamidi et al., 2018). Accordingly, YAP inhibition enhances the generation of insulin-producing β cells from stem cell-derived progenitors (Rosado-Olivieri et al., 2019). This dependence of pancreatic progenitor fate on F-actin organization has been exploited using latrunculins to directly perturb the actin cytoskeleton.

Latrunculin B (LatB), like the related Latrunculin A (LatA), binds G-actin monomers and prevents their polymerization, causing rapid F-actin disassembly (Morton, Ayscough, & Mclaughlin, 2000; Spector, Shochet, Kashman, & Groweiss, 1983). It is thus a useful tool for studying cellular responses to F-actin disruption (Morton et al., 2000). Treatment of fibroblasts with LatA resulted in accelerated actin mRNA degradation together with reduced vinculin RNA abundance (Bershadsky et al., 1995; Lyubimova, Bershadsky, & Ben-Ze’ev, 1997).

Recent studies have shown that latrunculin treatment induces differentiation of endocrine progenitor cells through expression of NEUROG3 (Hogrebe, Augsornworawat, Maxwell, Velazco-Cruz, & Millman, 2020; Hogrebe et al., 2026; Mamidi et al., 2018; Tian et al., 2025). LatB increases the expression of salt-inducible kinases (SIKs), which suppress YAP1 nuclear activity and promote the specification of NEUROG3⁺ endocrine progenitor cells. While these studies established an important connection between actin organization and endocrine differentiation, the broader cellular response to acute actin depolymerization remains less well characterized. In particular, it remains unclear whether disruption of actin assembly produces wide transcriptional changes among actin and its interacting proteins, and whether the physical and functional coupling among actin, microtubule, and intermediate filament networks is reflected in cross-cytoskeletal changes at the level of gene expression. Here we revisit the experiments by Tian et al. (Tian et al., 2025) who performed bulk RNA-seq over successive time points within a 24 hours window following LatB treatment. We re-analyzed these time-resolved data to characterize the broader transcriptomic response to actin depolymerization and the candidate regulatory programs associated with it. We perform a broader analysis by looking at the whole cell response within the context of what these measurements may reveal not only for differentiation, but also for cytoskeletal homeostasis, mechanotransduction, stress responses, and broader cellular pathway perturbations.

In the following sections we start by looking at the effects of LatB on the main genes of the actin system, monitoring both their time-averaged (“pooled”) as well as time-resolved response. This analysis is then extended to the microtubule and intermediate filament systems, and more broadly to genes involved in various cellular functions. Finally, we perform transcription factor activity analysis with decoupleR and ChEA3 to look into the candidate transcriptional regulators associated with these changes.

## Materials and Methods

### Dataset

Bulk RNA-seq data were obtained from the Gene Expression Omnibus (GEO) repository (Edgar, Domrachev, & Lash, 2002) under accession number GSE275775 (Tian et al., 2025). The dataset comprises 16 samples from bipotent pancreatic progenitors derived from human pluripotent stem cells, collected across four LatB treatment timepoints (3, 6, 12, and 24 hr) and two control timepoints (0 and 24 hr)(Tian et al., 2025). Raw sequencing data are available through the Sequence Read Archive (SRA)(Leinonen, Sugawara, & Shumway, 2011) with run accessions SRR30416715-SRR30416730 as follows (Tian et al., 2025): Control 0hr (SRR30416728, SRR30416729, SRR30416730), representing baseline untreated bipotent pancreatic progenitors, LatB 3hr (SRR30416723, SRR30416724, SRR30416725), LatB 6hr (SRR30416720, SRR30416721, SRR30416722), LatB 12hr (SRR30416718, SRR30416719), LatB 24hr (SRR30416715, SRR30416716, SRR30416717), and Control 24hr (SRR30416726, SRR30416727), serving as control to distinguish treatment effects from temporal changes in culture.

#### Read Alignment and Quantification

Raw sequencing reads were aligned to the human reference genome GRCh38,(Schneider et al., 2017; Yates et al., 2026) (GCA_000001405.15) using STAR v2.7.11b (Dobin et al., 2013). Gene-level counts were produced using --quantMode GeneCounts, all other parameters used STAR defaults. Gene annotation was provided by Ensembl release 110 (Homo_sapiens.GRCh38.110.gtf). The STAR genome index was generated with a splice junction overhang of 100 bp.

In the study of Tian et al.(Tian et al., 2025), a total of 516,723,347 single-end reads (75 bp) were sequenced across the 16 samples, with individual libraries ranging from 26.2 to 37.7 million reads (mean: 32.3 million reads per sample). We found alignment quality was consistently high across all samples, with uniquely mapped reads ranging from 92.00% to 92.48% (mean: 92.24%). Multi-mapped reads accounted for 6.28% to 6.72% of reads (mean: 6.43%), while unmapped reads due to insufficient length comprised only 1.06% to 1.35% (mean: 1.21%). The overall mismatch rate per base was low, ranging from 0.35% to 0.41% (mean: 0.36%), indicating high-quality sequencing data.

Gene-level raw read counts were obtained directly from STAR’s built-in quantification (Dobin et al., 2013) (version STAR v2.7.11b) using the ReadsPerGene.out.tab output files. The unstranded count column (column 2) was used, consistent with unstranded library preparation. Individual sample count files were compiled into a unified gene count matrix containing 62,754 genes across all 16 samples (Supplementary Data File 1). Genes with fewer than 10 total reads summed across all samples were removed prior to analysis, retaining 28,058 genes. Raw read counts were verified to be integers, as required for DESeq2 analysis (Love, Huber, & Anders, 2014). Gene identifiers were retained in Ensembl format (ENSG IDs) for differential expression analysis and subsequently converted to gene symbols using the org.Hs.eg.db annotation package (Gentleman et al., 2004) for functional annotation and pathway analyses. The 28,058 analyzed Ensembl identifiers corresponded to 22,685 unique gene symbols, genome-wide differential-expression counts reported in the Results are tallied per unique gene symbol, so this set of 22,685 expressed genes constitutes the denominator for those counts.

### Differential Expression Analysis

Differential expression analysis was performed using DESeq2 v1.44.0 (Love et al., 2014). The count data were modeled using a generalized linear model with negative binomial distribution. Each experimental condition and timepoint (Control 0hr, Control 24hr, LatB 3hr, LatB 6hr, LatB 12hr, LatB 24hr) was treated as an independent group. This design enables direct pairwise comparisons between any two such groups (Supplementary Data File 2). DESeq2 normalization was performed using the median-of-ratios method (size factor normalization)(Anders & Huber, 2010), which accounts for differences in library size and RNA composition between samples. Size factors ranged from 0.828 to 1.166 across the 16 samples (coefficient of variation = 10.7%), indicating comparable sequencing depth across conditions. Dispersion estimates were obtained using the DESeq2 default shrinkage approach (Love et al., 2014), and statistical significance was assessed using the Wald test. Log₂ fold change values are unshrunken maximum likelihood estimates (MLE) as returned by DESeq2’s results() function, LFC shrinkage was not applied.

For pairwise comparisons, multiple testing correction was applied using the Benjamini-Hochberg procedure (Benjamini & Hochberg, 1995) to control the false discovery rate (FDR). MA plots were used to compare the combined (pooled) LatB data over all time points to the combined control data at 0 and 24 hrs (Figures 1, 4, and 5). In the MA plots, the x-axis represents the mean of DESeq2 size factor-normalized counts averaged across all 16 samples (baseMean), extracted directly from DESeq2 results output and log₂-transformed for display. The y-axis represents the log₂ fold change from the pooled DESeq2 analysis comparing all LatB-treated samples against all control samples. We used two criteria to evaluate significance in pooled and pairwise comparisons with respect to controls. Genes with adjusted p-value (padj) less than 0.05 in at least one timepoint (or the given set of timepoints for pairwise comparisons) were classified as P-significant. P-significant genes that also satisfy |log₂ fold change| > 0.5 (corresponding to ≥1.41-fold change) at the same timepoint were classified as PFC-significant.

**Figure 1.**
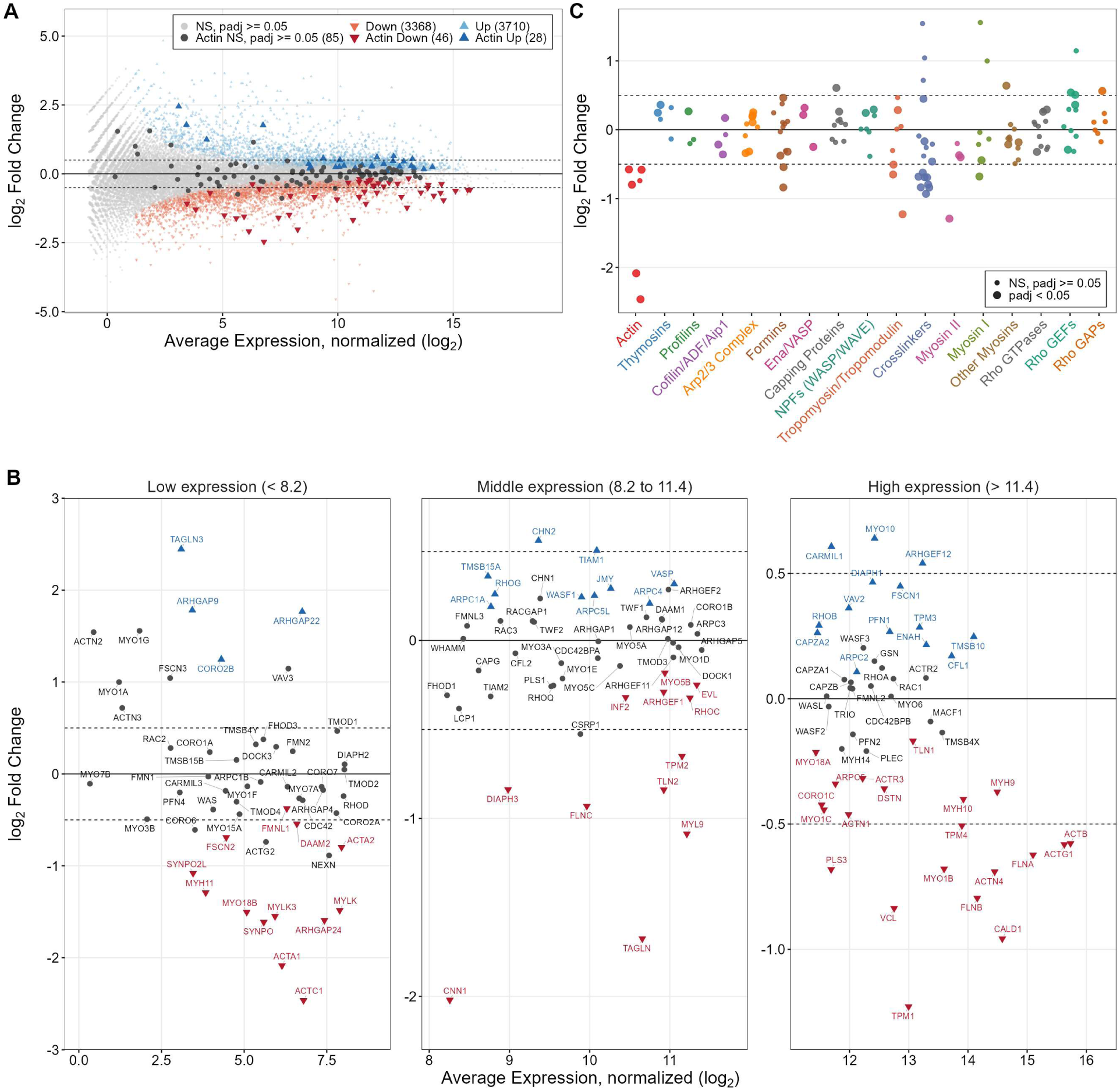
Genome-wide and actin cytoskeleton differential expression following Latrunculin B treatment in bipotent pancreatic progenitors. **(A)** MA plot showing genome-wide differential expression between LatB-treated (pooled across 3, 6, 12, and 24 hr timepoints) and control samples (pooled 0 and 24 hr timepoints). Colors indicate P-significance with respect to the padj value. Gray points indicate non-P-significant genes (padj ≥ 0.05), blue and orange points denote P-significantly downregulated (n = 3,414) and P-significantly upregulated (n = 3,738) genes, respectively. Actin cytoskeleton genes (n = 159) are highlighted in gray (non-P-significant) and color (P-significantly differentially expressed). Dashed lines mark log₂FC thresholds of ±0.5. (B) Detailed views of panel A with labels for individual genes. (C) Category-resolved expression changes of 138 actin cytoskeleton genes organized into 17 functional categories, ordered from core actin regulatory machinery (left) through motor proteins (center) to Rho GTPase signaling (right). Point size indicates P-significance (large: padj < 0.05)

For time course visualization of individual gene expression dynamics, DESeq2 size factor-normalized counts were plotted directly without log transformation, as comparisons were made within individual genes across conditions rather than between genes. The size factor-normalized counts were extracted using the counts() function with normalized=TRUE (Love et al., 2014).

For heatmap visualization (Figure 9), log₂ fold change values from the per-timepoint DESeq2 comparisons were arranged into a gene × timepoint matrix. Rows were hierarchically clustered within each annotated branch using Euclidean distance and complete linkage, columns were not clustered and represent the fixed time course (LatB 3hr, 6hr, 12hr, 24hr, and Control 24hr, each vs. Control 0hr). The heatmap was rendered using the ComplexHeatmap R package (Gu, Eils, & Schlesner, 2016) with a diverging color scale centered at zero (range −2 to +2 log₂FC).

For Figure 7, we classified PFC-significant differentially expressed genes into different cell functional categories using a vote-based Gene Ontology annotation approach. GO term annotations were retrieved from org.Hs.eg.db v3.19.1 via AnnotationDbi, yielding 135,695 BP annotations across 15,038 genes (Supplementary Data File 3). For each gene, GO BP term names were matched against custom category-specific keyword lists, and the gene was assigned to the category whose keywords produced the highest number of matches (minimum one match required), genes with no matches were classified as ’Other’. Structural overrides were then applied to reassign canonical gene families to their primary category regardless of vote outcome (e.g., TUBA/B/D/E/G and KIF to Microtubules, ACTA/B/C/G and MYH/MYL/MYO to Actin Cytoskeleton, RAB and SYT to Vesicle Trafficking, CACN, SCN, and KCN to Ion Transport). Finally, gene biotype annotations were used to reassign lncRNAs and pseudogenes to dedicated categories. This yielded 18 final functional categories displayed in Figure 7 (Supplementary Data File 4).

### Transcription Factor Activity Inference

To identify transcription factors (TFs) mediating the response to LatB treatment, TF activity inference (Figure 8) was performed using the decoupleR package (v2.6.0)(Badia-I-Mompel et al., 2022) with the CollecTRI network, which contains TF-target interactions for 737 human TFs compiled from 12 different resources (Müller-Dott et al., 2023). The univariate linear model (run_ulm) of decoupleR was applied to DESeq2 Wald statistics (log₂ fold change divided by standard error) as input for each timepoint comparison versus Control 0hr (Badia-I-Mompel et al., 2022). This statistic provides a measure of gene-level differential expression that accounts for both effect size and uncertainty. Positive activity scores indicate TF activation relative to control, negative scores indicate inactivation. A mean activity score was computed across the four timepoints.

The 737 TFs were classified into functional signaling pathway categories using a two-layer rule-based approach. In Layer 1, each TF gene symbol was tested against a set of manually curated regular expression patterns defining the pathway categories, applied in priority order with the first match assigned (e.g., SRF|MKL|MYOCD → MRTF/SRF, TEAD|YAP1|WWTR1 → Hippo/YAP-TEAD, SMAD →SMAD/TGF-β, NOTCH|HES|HEY|RBPJ → Notch Signaling). Category membership was defined by manual curation based on established literature knowledge of pathway components, Layer 1 captures TFs belonging to well-characterized signaling pathways whose members follow recognizable gene-name conventions, while Layer 2 extends coverage to TFs whose pathway membership is encoded in functional annotation rather than gene nomenclature. In Layer 2, TFs not assigned by Layer 1 were annotated using GO Biological Process terms retrieved from org.Hs.eg.db as the sole external annotation source, with GO term descriptions matched against category-specific keyword lists (e.g., “response to hypoxia” → HIF/Hypoxia, “unfolded protein response” → UPR/ER Stress). TFs remaining unclassified after both layers, and any category containing fewer than four members, were consolidated into “Other TFs,” yielding 24 final categories ordered by median mean activity score (Figure 8).

Additionally, TF enrichment analysis was performed using ChEA3 (ChIP-X Enrichment Analysis 3)(Keenan et al., 2019) via the rChEA3 R package. ChEA3 takes as input discrete lists of significantly differentially expressed genes and identifies TFs whose known target gene sets are statistically overrepresented. PFC-significant differentially expressed genes from the pooled LatB versus control comparison (padj < 0.05, |log₂FC| > 0.5) were queried separately for upregulated and downregulated gene sets. The gene set of the top 300 upregulated and downregulated genes, ranked by their |log₂FC| values, were given as inputs, after excluding non-protein coding genes. TF ranks were determined using the integrated mean rank across all ChEA3 libraries.

### Statistical Software

All analyses were performed in R (v4.5.2)(‘R Core Team (2024) R A Language and Environment for Statistical Computing. R Foundation for Statistical Computing. - References - Scientific Research Publishing’, n.d.). Key packages included: DESeq2(Love et al., 2014) for differential expression analysis, ggplot2, ComplexHeatmap (Gu et al., 2016) and pheatmap for visualization, dplyr for data manipulation, org.Hs.eg.db and AnnotationDbi (Gentleman et al., 2004) for gene annotation, and ggrepel for plot labeling.

### Use of AI

Claude Sonnet 5 and ChatGPT version 5.5 were used in the creation of scripts and document editing. The authors reviewed and edited the content as needed and take full responsibility for the content of the publication.

## Results

### Latrunculin B Treatment Induces Widespread Transcriptional Changes Involving Multiple Actin Cytoskeleton Associated Genes

Time-resolved bulk RNA-seq data from bipotent pancreatic progenitors derived from human pluripotent stem cells, treated with LatB, provided the basis of our analysis. They were obtained from the Gene Expression Omnibus (GEO) repository(Edgar et al., 2002) under accession number GSE275775(Tian et al., 2025). The dataset includes 16 total samples collected across four timepoints following LatB treatment (Tian et al., 2025) (3, 6, 12, 24 hrs) and control samples at 0 and 24 hrs. The cells were differentiated from SA121 and HuES4 human embryonic stem cell lines following an established protocol to generate PDX1⁺/NKX6.1⁺ progenitors by day 19 (Tian et al., 2025). At the time of LatB treatment, cells had formed a confluent monolayer with cell-cell contacts and were attached to the culture substrate through integrin-mediated adhesions to the extracellular matrix (Tian et al., 2025). This baseline cellular organization is relevant because the transcriptional response to actin disruption is likely to depend on the pre-existing cytoskeletal and mechanical state of the cells: both MRTF/SRF and YAP/TAZ are sensitive to actin organization and cytoskeletal tension generated through these cell-ECM and cell-cell adhesion complexes,(Dupont et al., 2011). We quantified the transcriptional response with differential expression analysis using DESeq2.

To provide an overall view of the transcriptional response to LatB, we first pooled all LatB treated timepoints and compared them to the pooled control samples at 0 and 24 hr using an MA plot (Figure 1A, Supplementary Data File 5). This analysis provides a comparison between the time-averaged response to LatB-treated and the time-averaged response of untreated cells. We observe a significant genome-wide LatB perturbation with the number of perturbed genes far greater than the differential expression changes between 0 and 24 hr of control samples (MA plot in Supplementary Figure 1). Across individual timepoints, 11,975 unique genes (52.8% of 22,685 expressed genes) showed P-significant differential expression (padj < 0.05) in at least one timepoint, of which 6,632 (55.4%) also exceeded the fold-change threshold (|log₂FC| > 0.5) at the same timepoint and were considered PFC-significantly differentially expressed.

To examine the direct effects on the actin cytoskeleton, we specifically analyzed a selection of 159 actin cytoskeleton-associated genes encompassing core structural components (actin monomers, crosslinkers, tropomyosins), regulatory proteins (Arp2/3 complex, formins, capping proteins), motor proteins (myosins), and upstream signaling components (Rho GTPases, GEFs, and GAPs). These genes, individually indicated in Figure 1A and in Supplementary Data File 6, were selected after first assembling a set of core actin genes querying GO, KEGG, MSigDB, BioMart, and UniProt databases followed by manual curation to verify their direct roles in actin regulation and structure. The selected actin cytoskeleton genes showed a higher rate of PFC-significant differential expression in response to LatB (24.5%, 39/159 genes, padj < 0.05 and |log₂FC| > 0.5) compared to the genome-wide average (12.2% of 28,058 genes). Of these, 30 genes were significantly downregulated compared to 9 significantly upregulated genes (Figure 1A where colors indicated P-significance with respect to padj value only).

Differential expression of the selected actin cytoskeleton genes across 17 functional categories (containing 138 of the 159 genes) revealed category-specific response patterns in this pooled analysis (Figure 1C, Supplementary Data File 7). Actin monomers exhibited the most pronounced downregulation (mean log₂FC = −1.21), with the weakly expressed cardiac and skeletal muscle actins (ACTC1, ACTA1) showing the largest decreases (log₂FC = −2.46 and −2.09, respectively). Contractile machinery components were similarly affected, with Myosin II heavy chains (mean log₂FC = −0.57) and tropomyosins/tropomodulins (mean log₂FC = −0.24) showing consistent downregulation. Formins, Arp2/3 complex subunits and nucleation-promoting factors (NPFs/WASP/WAVE) showed mixed trends. Crosslinking proteins including filamins, α-actinins, and focal adhesion components were predominantly downregulated (mean log₂FC = −0.22), with filamin C (FLNC, log₂FC = −0.93), talin 2 (TLN2, log₂FC = −0.84), vinculin (VCL, log₂FC = −0.84) and plastin 3 (PLS3, log₂FC = −0.68) among the most affected. Rho GTPase signaling components showed divergent responses: while Rho GTPases themselves remained relatively unchanged, their activators (Rho GEFs) were predominantly upregulated (mean log₂FC = +0.22), including ARHGEF12 (log₂FC = +0.54) and VAV2 (log₂FC = +0.36). This upregulation of Rho GEFs and GAPs, which activate and deactivate Rho GTPases (Rossman, Der, & Sondek, 2005), may represent a compensatory feedback mechanism whereby cells attempt to adjust cytoskeletal integrity in response to LatB-induced depolymerization,(Morton et al., 2000; Spector et al., 1983).

### Expression of Actin Genes over time in response to LatB

To examine the temporal dynamics of the actin cytoskeleton genes, we analyzed DESeq2 size-factor normalized count trajectories across the LatB treatment time course (0, 3, 6, 12, and 24 hours), for each separate experimental sample(Tian et al., 2025). The time response of each gene is listed in Supplementary Data File 8. The time response of representative core actin cytoskeleton components is shown in Figures 2 and 3. To better track the response of cytoskeletal proteins of varying cellular concentrations, we plot counts across the time course for each individual gene. Additional time course plots in Supplementary Data File 9 show the response of several actin cytoskeleton families (crosslinkers, capping proteins, etc). Notably, for nearly all genes, the control (no LatB) counts at 0h have very similar values to counts at 24h, this indicates that deviation from these values at intermediate times show the specific response to LatB.

**Figure 2.**
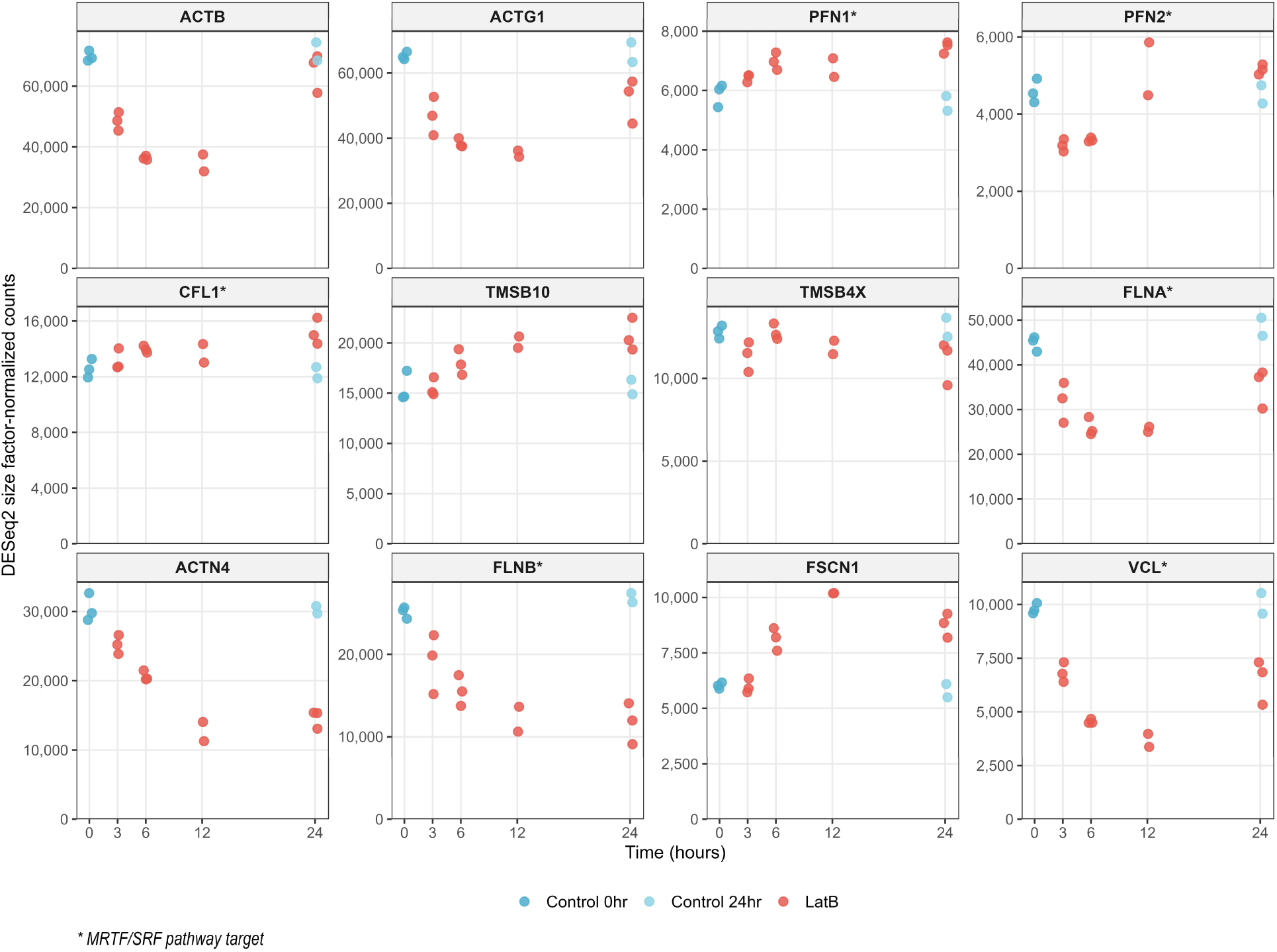
Time-resolved expression of core actin cytoskeleton genes following LatB treatment. Time course of DESeq2 size-factor-normalized counts for representative actin cytoskeleton genes. Each point corresponds to a separate sample(Tian et al., 2025). Blue points show control (dark blue: 0hr, light blue: 24hr) and red show LatB-treated samples. Actin genes (ACTB, ACTG1) show downregulation within 3 hours. Profilins (PFN1, PFN2) and cofilins (CFL1) remain relatively stable. G-actin sequestering thymosins (TMSB10, TMSB4X) show variable responses. Crosslinking proteins (FLNA, FLNB, ACTN4) and the focal adhesion component vinculin (VCL) exhibit progressive downregulation. FSCN1 shows modest upregulation. Asterisks denote MRTF/SRF pathway target genes (Supplementary Data File 10).

**Figure 3.**
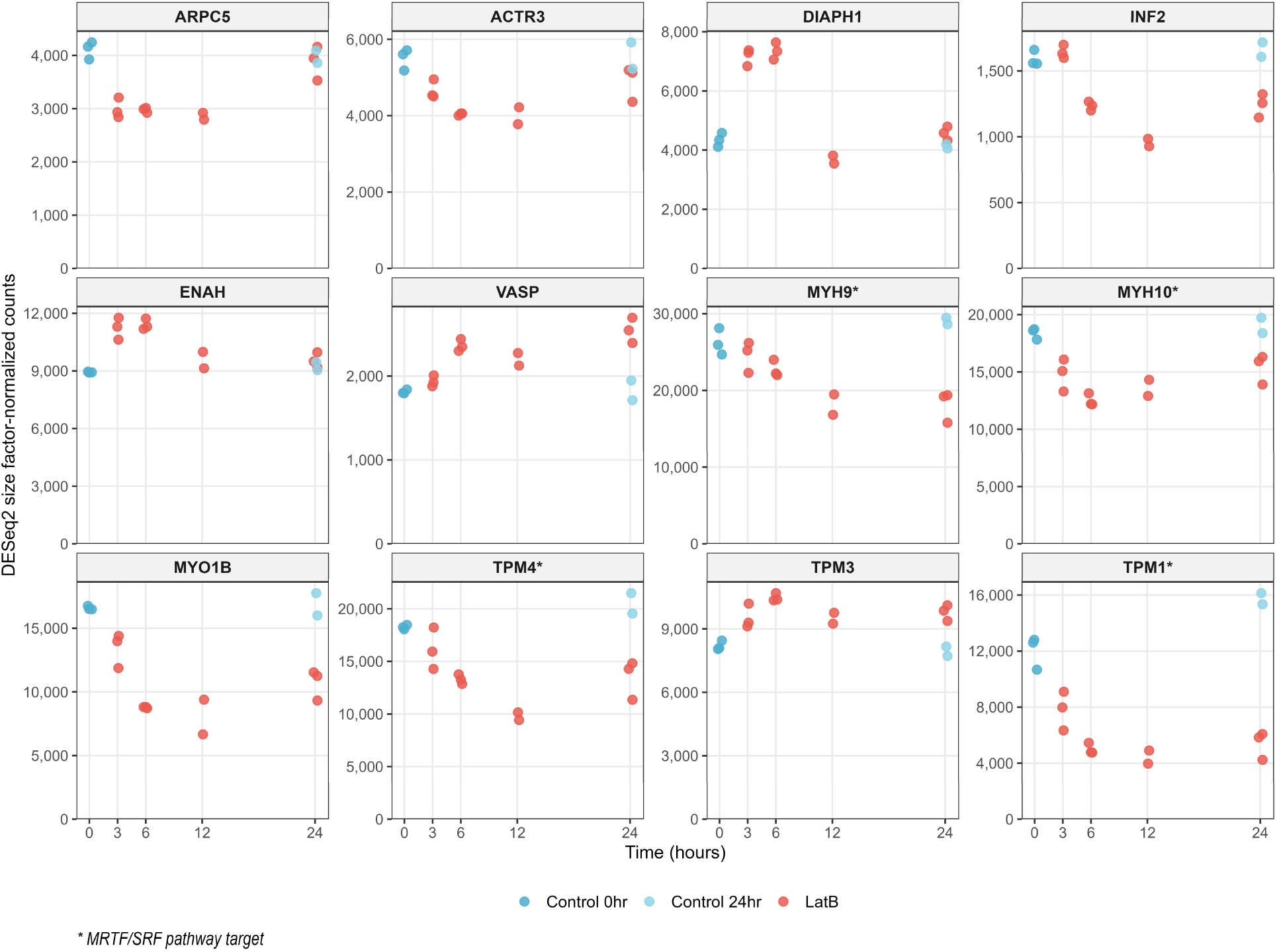
Time-resolved expression of actin nucleation machinery and motor proteins following Latrunculin B treatment. Same as Figure 2, for selected Arp2/3 complex subunits (ARPC5, ACTR3), formins (DIAPH1, INF2), Ena/VASP proteins (ENAH, VASP), non-muscle myosins (MYH9, MYH10, MYO1B), and tropomyosins (TPM1, TPM3, TPM4). Arp2/3 complex subunits show recovery at 24 hours. Formins display divergent responses. Myosins and tropomyosins show sustained downregulation throughout the time course. Asterisks denote MRTF/SRF pathway target genes.

The analysis reveals that the expression of the two most highly expressed actins in these cells, β-actin and γ-actin (ACTB, ACTG1), is noticeably reduced after 3 hours, reaching a maximum reduction by a factor of order 2 between 6-12 hrs post LatB treatment, the expression recovers back to control values after 24 hrs (Figure 2). Earlier LatA studies had shown reduction in actin mRNA through mRNA degradation(Bershadsky et al., 1995; Lyubimova et al., 1997). The reduction in actin expression could also reflect the cell’s response to the increased monomeric actin pool after LatB treatment through the MRTF/SRF pathway that senses actin abundance through measurements of actin monomer concentrations,(Salvany, Muller, Guccione, & Rørth, 2014; Sotiropoulos, Gineitis, Copeland, & Treisman, 1999).

The comparatively smaller increase of actin monomer binding protein β-thymosin, TMSB10, and profilin 1, PFN1, is consistent with increased expression of proteins that can bind to the excess of actin monomers produced by LatB. Unlike ACTB and ACTG1 however, their expression does not appear to recover back to control values. Notably, profilin 2, PFN2, follows the opposite trend which matches the time course of the two actins. Prior work has shown MRTF regulates the cellular levels of both profilin isoforms through STAT1, largely by post-transcriptional mechanisms that include altered cellular profilin retention, and that profilin levels can in turn influence MRTF/SRF signaling(Joy, Gau, Castellucci, Prywes, & Roy, 2017). PFN1 and PFN2 can also have distinct effects on actin organization, with PFN2 preferentially supporting Ena/VASP-dependent actin assembly (Mouneimne et al., 2012).

The crosslinking proteins FLNA, FLNB, ACTN4, and VCL are all downregulated, without fully recovering to control values after 24 hrs. By contrast, FSCN1 shows a continuously increasing upregulation. Cofilin CFL1 did not show any notable change. Components of the Arp2/3 branched actin nucleation complex displayed a divergent response, with the most notable change being a slight downregulation of ARPC5 and ACTR3. Actin polymerization regulators formins DIAPH1 and INF2, as well as ENAH and VASP showed a variable response. DIAPH1 and ENAH had an apparently quick increase after 3-6hr followed by a reduction to control levels after 12 hr. VASP showed a mild continuous increase with time while INF2 showed a reduction between 6-12 hrs followed by recovery to control levels at 24 hrs. The expression of myosin II heavy chains MYH9 and MYH10, the two major non-muscle myosin II isoforms (IIA and IIB), generally decreased over time, indicating a reduced expression of contractile machinery. The most highly-expressed myosin I, MYO1B, as well as tropomyosins TPM1, TPM2, TPM4, had a similar decrease over time. TPM3 by contrast had a mild increase. The most notable change of Rho family GEF and GAP regulators was ARHGEF12, which showed immediate elevation at 3 hours that persisted throughout the time course, and VAV2 which similarly exhibited sustained increases above control levels between 6-12 hr (Supplementary Data File 9).

In conclusion, LatB produces a non-uniform change of expression across the actin system. Aspects of this variability have appeared in earlier studies. Increasing thymosin-β4 expression in fibroblasts led to increases in the concentration of actin and in myosin IIA, α-actinin, tropomyosin, talin and vinculin, while profilin did not increase(Golla et al., 1997). Similarly, increasing the G-actin/F-actin ratio through INF2 depletion reduced MRTF/SRF activity together with expression of several filament-associated and adhesion genes, including FLNA, TLN1, TPM1 and VCL(Fernández-Barrera et al., 2018). Altogether, these observations suggest that changes in actin polymerization state alters the relative expression of different components rather than maintaining fixed expression ratios.

### Transcriptional Changes of Microtubule and Intermediate Filament Cytoskeleton Genes in Response to LatB

Given the extensive interconnections between cytoskeletal systems (Dogterom & Koenderink, 2019), we next examined whether F-actin disruption affects the expression of microtubule and intermediate filament network components. Previous work has shown that INF2 silencing in epithelial RPE-1 cells leads to reduction of α-tubulin acetyltransferase 1 (α-TAT1) expression through the actin-MRTF/SRF transcriptional circuit, coupling G-actin levels to microtubule acetylation(Fernández-Barrera et al., 2018). The following analysis addresses the broader question of whether LatB perturbation remains confined to actin-related genes or extends to other cytoskeletal systems through transcriptional crosstalk.

We analyzed 123 selected microtubule-associated genes and found extensive transcriptional changes. Of the 123 analyzed genes, 47 showed P-significant differential expression (padj < 0.05), as indicated by the color coding in Figure 4A. Of these, 14 met the stricter PFC-significance threshold (padj < 0.05 and |log₂FC| > 0.5), comprising 8 upregulated and 6 downregulated genes.

**Figure 4.**
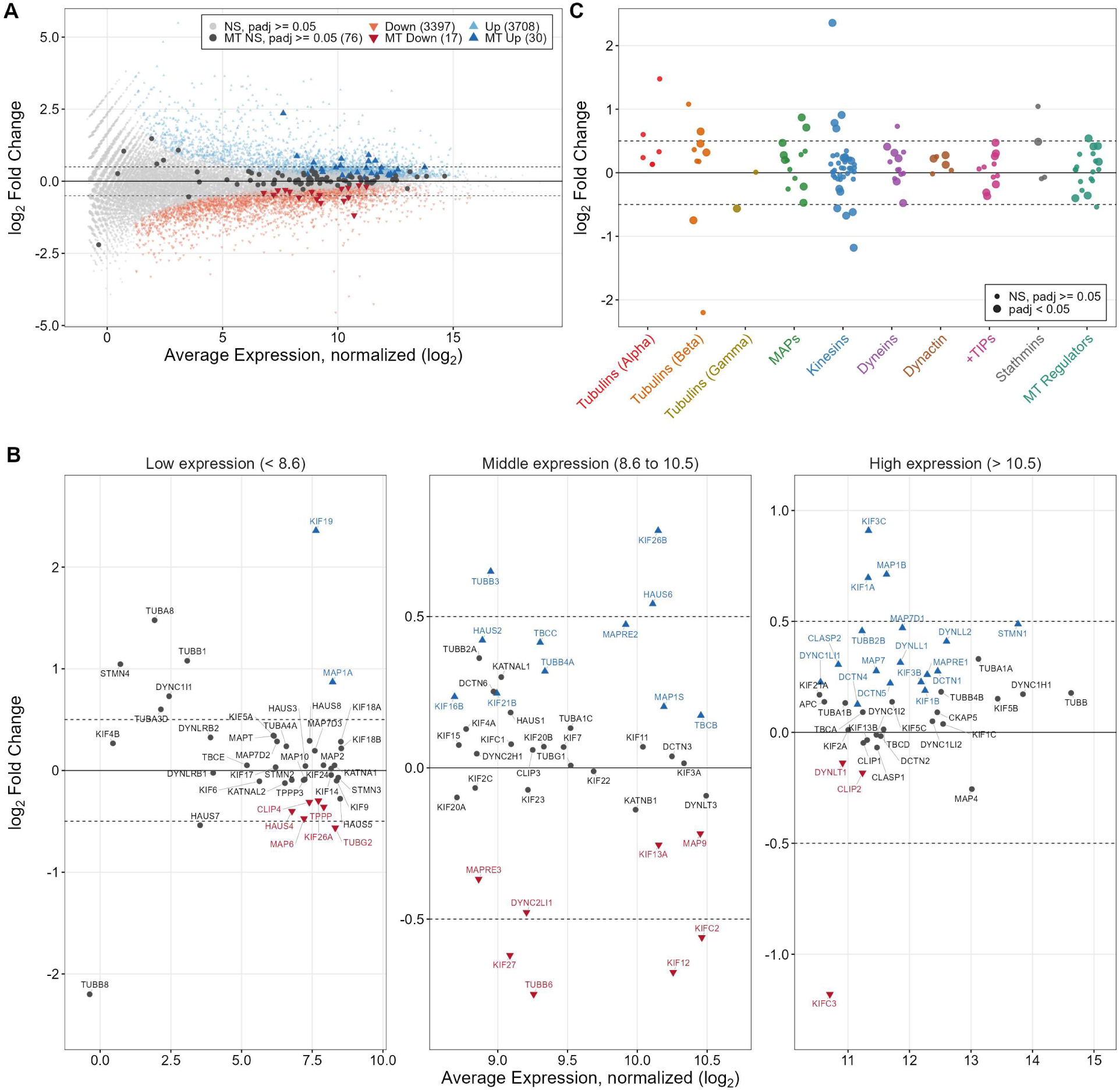
Microtubule network gene expression changes following LatB treatment. (A) MA plot displaying genome-wide differential expression comparing pooled LatB-treated versus Control samples. Gray points indicate non-P-significant genes, orange points show P-significantly downregulated genes (3,414), blue points show P-significantly upregulated genes (3,738). Microtubule-associated genes are highlighted in gray (non-P-significant) and color (P-significantly differentially expressed). Key MT genes are labeled. (B) Detailed views of panel A with labels for individual genes. (C) Log₂ fold change distribution of microtubule genes organized by functional category. Categories include α-tubulins, β-tubulins, γ-tubulins, microtubule-associated proteins (MAPs), kinesins, dyneins, dynactin complex, plus-end tracking proteins (+TIPs), stathmins, and MT regulators. Large dots indicate P-significant differential expression (padj < 0.05), small dots indicate non-P-significant changes.

Visualization of fold changes by functional category demonstrated that the response was not uniform across microtubule components (Figure 4C). Kinesins showed the most variable response, with KIF19 displaying the strongest upregulation (log₂FC = +2.36) while other kinesins were significantly downregulated. Microtubule-associated proteins (MAPs) including MAP1A and MAP1B were predominantly upregulated, suggesting altered expression of microtubule-stabilizing machinery. Among tubulin genes, β-tubulins showed mixed responses with TUBB3 upregulated and TUBB6 downregulated, α-tubulins remained largely unchanged, and among γ-tubulins TUBG2 was PFC-significantly downregulated (log₂FC = −0.56, padj < 0.001) while TUBG1 was stable. This pattern suggests that cells remodel microtubule-dependent processes, particularly motor protein-based transport and MT stability, in response to LatB disruption.

Consistent with the previously described α-TAT1 decrease, ATAT1 showed a modest P-significant decrease following LatB treatment, although it did not meet the PFC-significance threshold. Among the appreciably expressed PFC-significant genes involved in tubulin post-translational modifications, the tubulin detyrosinase VASH2 increased, whereas the polyglutamylase TTLL9, the glycylase TTLL3, and the deglutamylases AGBL2 and AGBL3 decreased (Supplementary Data File 5). Thus, LatB led to a transcriptional response involving multiple tubulin modification pathways.

We examined 36 intermediate filament genes that were present in the dataset. Of these, 12 showed P-significant differential expression (padj < 0.05), with a pattern of class-specific responses (Figure 5A-C). Type I and Type II keratins showed coordinated downregulation, with KRT4 exhibiting the strongest suppression (log₂FC = -2.22), followed by KRT18 (log₂FC = -0.97) and KRT8 (log₂FC = -0.75). In contrast, Type IV neurofilaments NEFL and NEFH were significantly upregulated (log₂FC = +1.08 and +0.58, respectively). Type V lamins (LMNA, LMNB1, LMNB2) involved in nuclear envelope organization remained stable. IF-associated proteins including plakophilins (PKP2, PKP4) were downregulated. A transcriptional connection between actin depolymerization and intermediate filaments was previously observed in mesenchymal stem/stromal cells, where LatB rapidly induced several keratin and keratin-associated genes and was followed by increased keratin filament formation (Chang et al., 2014). In contrast, the response observed here is characterized by suppression of several keratins together with induction of neurofilament genes, suggesting that the intermediate-filament response to actin disruption may strongly depend on cellular context.

**Figure 5.**
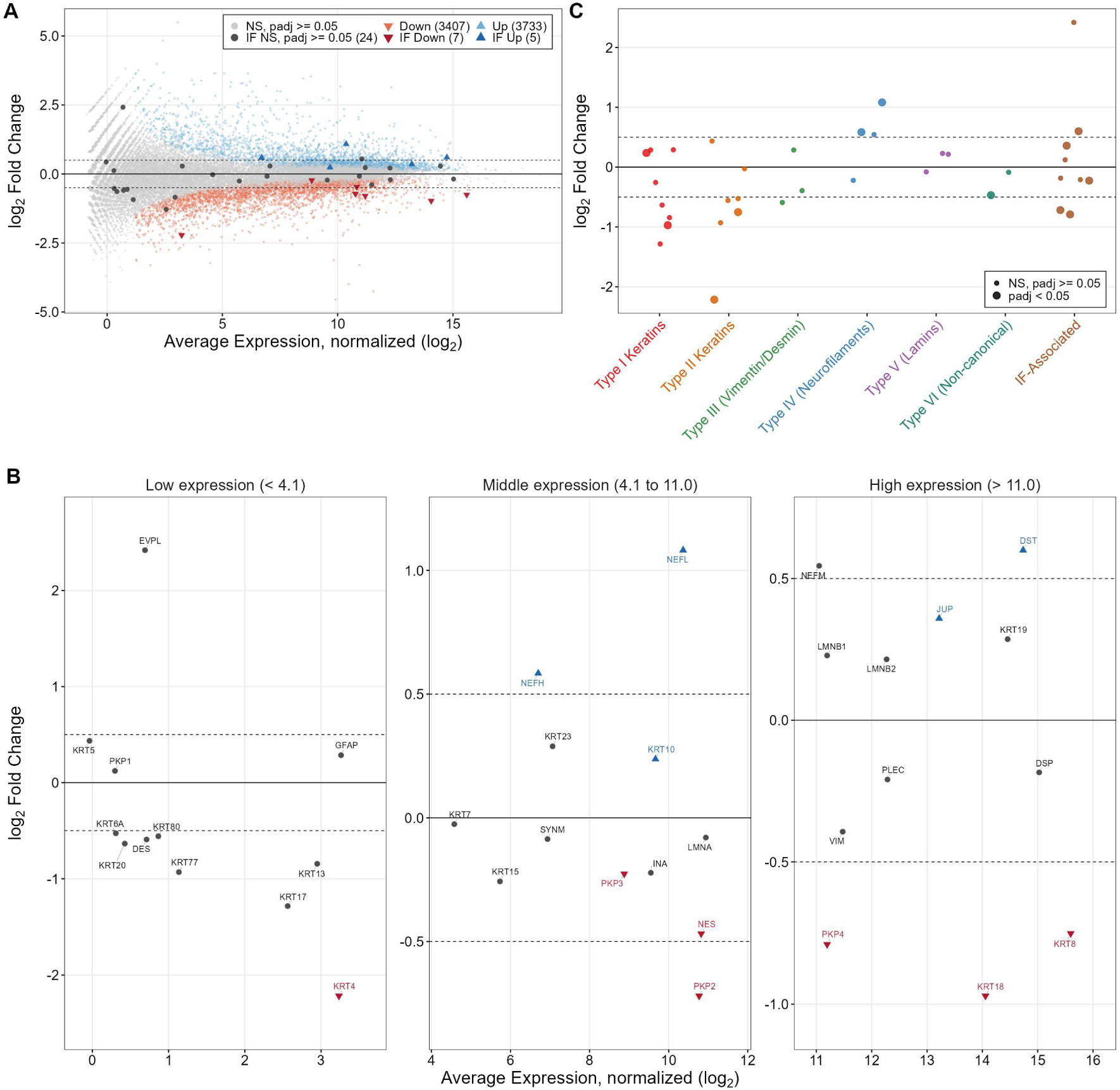
Intermediate filament network remodeling following LatB treatment. (A) MA plot displaying genome-wide differential expression comparing pooled LatB-treated versus Control samples. Gray points indicate non-P-significant genes, blue points show P-significantly downregulated genes (3,414), orange points show P-significantly upregulated genes (3,738). Intermediate filament genes are highlighted in gray (non-P-significant) and color (P-significant differentially expressed). (B) Detailed views of panel A with labels for individual genes including keratins (KRT4, KRT8, KRT18) and neurofilaments (NEFL, NEFH). (C) Log₂ fold change distribution of intermediate filament genes organized by IF type. Categories include Type I keratins, Type II keratins, Type III (vimentin/desmin), Type IV (neurofilaments), Type V (lamins), Type VI (non-canonical), and IF-associated proteins. Large dots indicate P-significant differential expression (padj < 0.05), small dots indicate non-P-significant changes.

To determine the overall relative timing of actin, microtubule and intermediate filament changes, we analyzed the temporal dynamics of differential expression (Figure 6). All three cytoskeletal systems showed P-significant transcriptional changes at the earliest timepoint examined (3hr) that persisted through 24 hr, with a combination of upregulated and downregulated genes of comparable magnitudes. We observe persistent downregulation of ACTA1, ACTC1, overexpression of TUBB1, and downregulation of KRT4, KRT13, KRT17 that intensified over later times, though we note that the expression levels of these genes are moderate to low (Figures 1, 4, 5).

**Figure 6.**
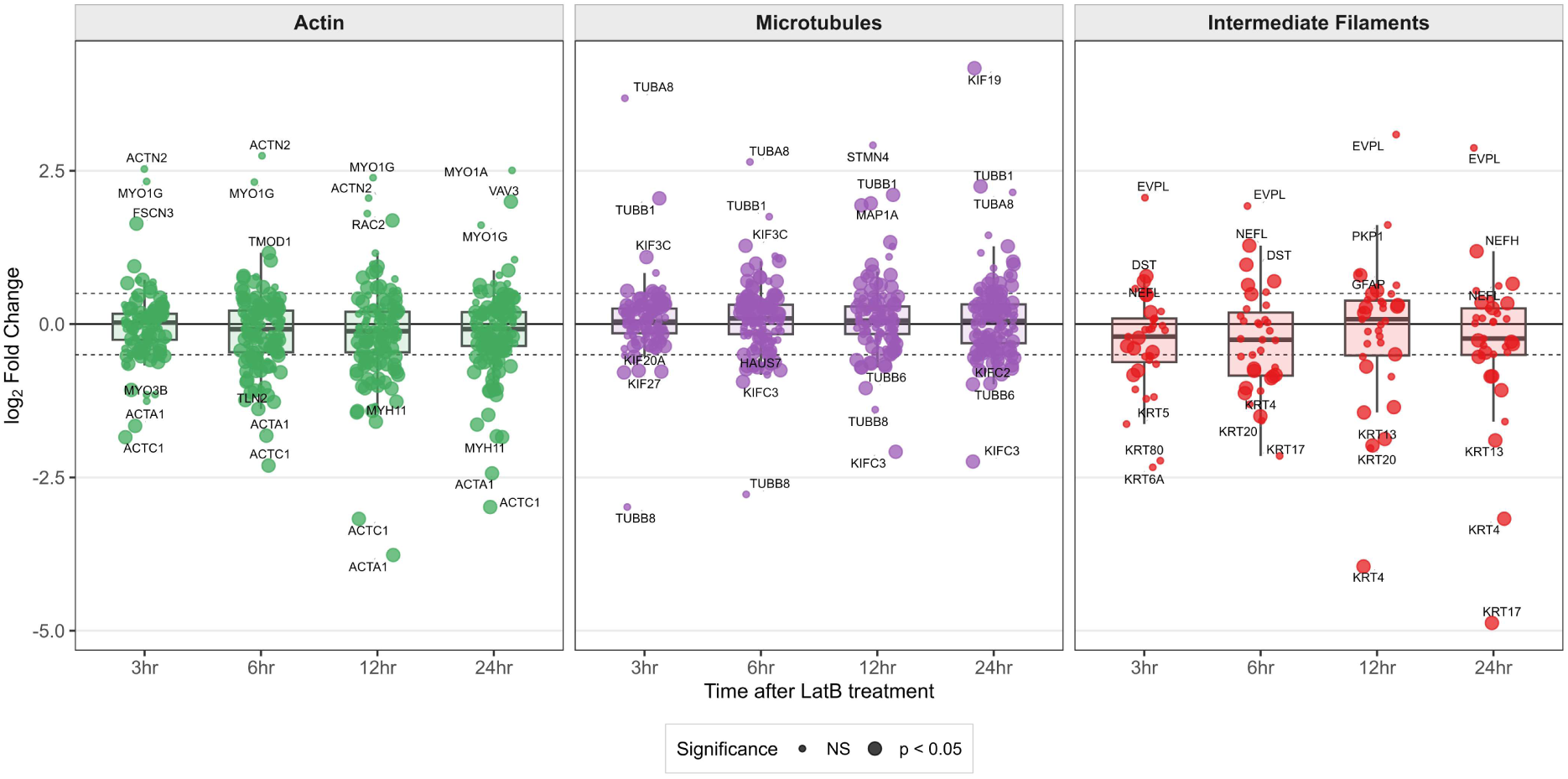
Temporal log₂ fold change distribution of cytoskeletal genes following Latrunculin B treatment. Boxplots (Box: IQR (25th–75th percentile), center line: median, whiskers: 1.5×IQR) show the distribution of log₂ fold change values for all genes in each cytoskeletal system at each LatB treatment timepoint (3, 6, 12, and 24 hr), relative to the 0 hr control. Large dots indicate P-significant differential expression (padj < 0.05) and small dots indicate non-P-significant changes. Dashed lines mark log₂FC thresholds of ±0.5.

Together, this analysis shows that transcriptional consequences of F-actin disruption extend well beyond the actin system itself, highlighting the interconnected nature of cytoskeletal networks even at the level of gene expression.

### Genome-wide Transcriptional Changes in Response to LatB

Previous RNA-seq analysis of LatB-treated primary endothelial cells revealed a broad transcriptional response within 4 hr, indicating that effects of actin depolymerization propagate beyond the cytoskeleton(Gegenfurtner et al., 2018). In order to further assess the full scale of the cellular response to LatB disruption beyond the actin, microtubule, and intermediate filament systems, we created expanded categories that correspond to additional cellular processes or functions, and plotted the corresponding numbers of differentially expressed genes across the time course (Figure 7, pairwise comparisons to 0hr Control). The categories of Figure 7 include expanded actin cytoskeleton, microtubule, and intermediate filaments, and were created by matching customized keyword terms relevant to each category to GO annotations in Biological Process ontology (see Methods).

**Figure 7.**
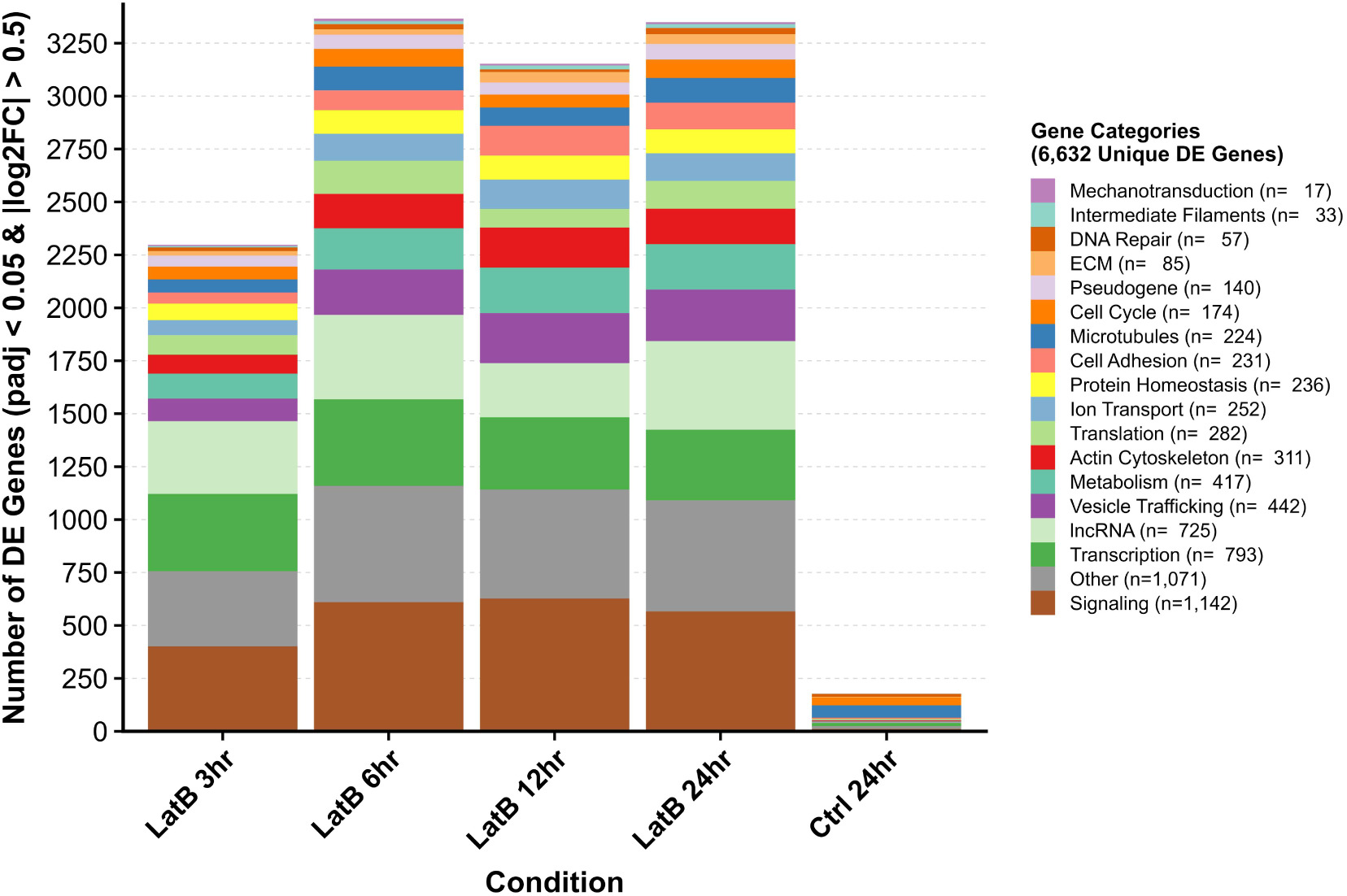
Temporal distribution of differentially expressed genes across functional categories following Latrunculin B treatment. Stacked bar plot showing PFC-significantly differentially expressed genes (padj < 0.05 and |log₂FC| > 0.5, pairwise comparison for each timepoint to 0hr Control) across 18 functional categories at each timepoint. The transcriptional response peaks at 6 hours and remains elevated through 24 hours. Legend indicates unique genes per category across all timepoints (n = 6,632 total unique PFC-significant DE genes).

The transcriptional response was extensive, with 2,298 PFC-significantly differentially expressed genes (padj < 0.05, |log₂FC| > 0.5) at 3 hours, peaking at 6 hours, and remaining for 24 hours. These numbers are much larger than the number of differentially expressed genes of untreated cells after 24 hours. The response extended broadly to genes involved in transcription, vesicle trafficking, metabolism, translation, ion transport, protein homeostasis.

The analysis confirms the overexpression of NEUROG3 ^,,^(Hogrebe et al., 2020; Mamidi et al., 2018; Tian et al., 2025) (Supplementary Figure 2) and SIK1, SIK2, SIK3 overexpression (Supplementary Figure 3) reported by Tian et al.(Tian et al., 2025) who demonstrated that SIKs transduce F-actin-triggered changes in YAP1 activity to promote endocrine progenitor differentiation. Our analysis reproduces the endocrine gene time response (Figure 1B in Tian et al., Supplementary Data File 8) and differentially expressed kinases (Figure 1C in Tian et al., Supplementary Data File 2). For example, NKX2-2 increased from log₂FC = +2.05 at 3 hours to +5.91 at 24 hours, while PAX4 and NEUROD1 showed similar progressive upregulation and RFX6 showed sustained upregulation from 3hr onward (Supplementary Data File 2), consistent with the endocrine differentiation program (Tian et al., 2025).

### Transcription Factors Involved in Response to LatB Treatment

We next sought to infer the transcription factors (TFs) associated with the differential expression in the preceding figures. The effects on transcription must be channeled through intermediary signaling molecules and transcription factor networks. We anticipate the involvement of the actin cytoskeleton, YAP and TAZ, and MRTF-SRF pathways as described in the Introduction, in addition to the multiple other pathways as evident in the extensive perturbation of the signaling genes (Figure 7) and the triggering of the endocrine progenitor differentiation program.

To infer a TF activity score, we used decoupleR to calculate a TF activity score by comparing the 0hr Control to each time point of the LatB treatment, using the CollecTRI gene regulatory network of 737 human TFs and their target genes. We calculated the average TF activity scores over all time points, where positive values correspond to activation with respect to 0hr and negative inactivation. Figure 8 and Supplementary Data File 11 show the results of this analysis with TFs classified into functional categories. TFs were classified into functional categories by gene-symbol pattern matching for pathway membership, with GO Biological Process annotation applied to remaining TFs. Categories with fewer than four members were merged into “Other TFs” yielding 24 categories ordered by median activity score. The top 10 activated and top 10 inactivated TFs are shown in Supplementary Figure 4.

**Figure 8:**
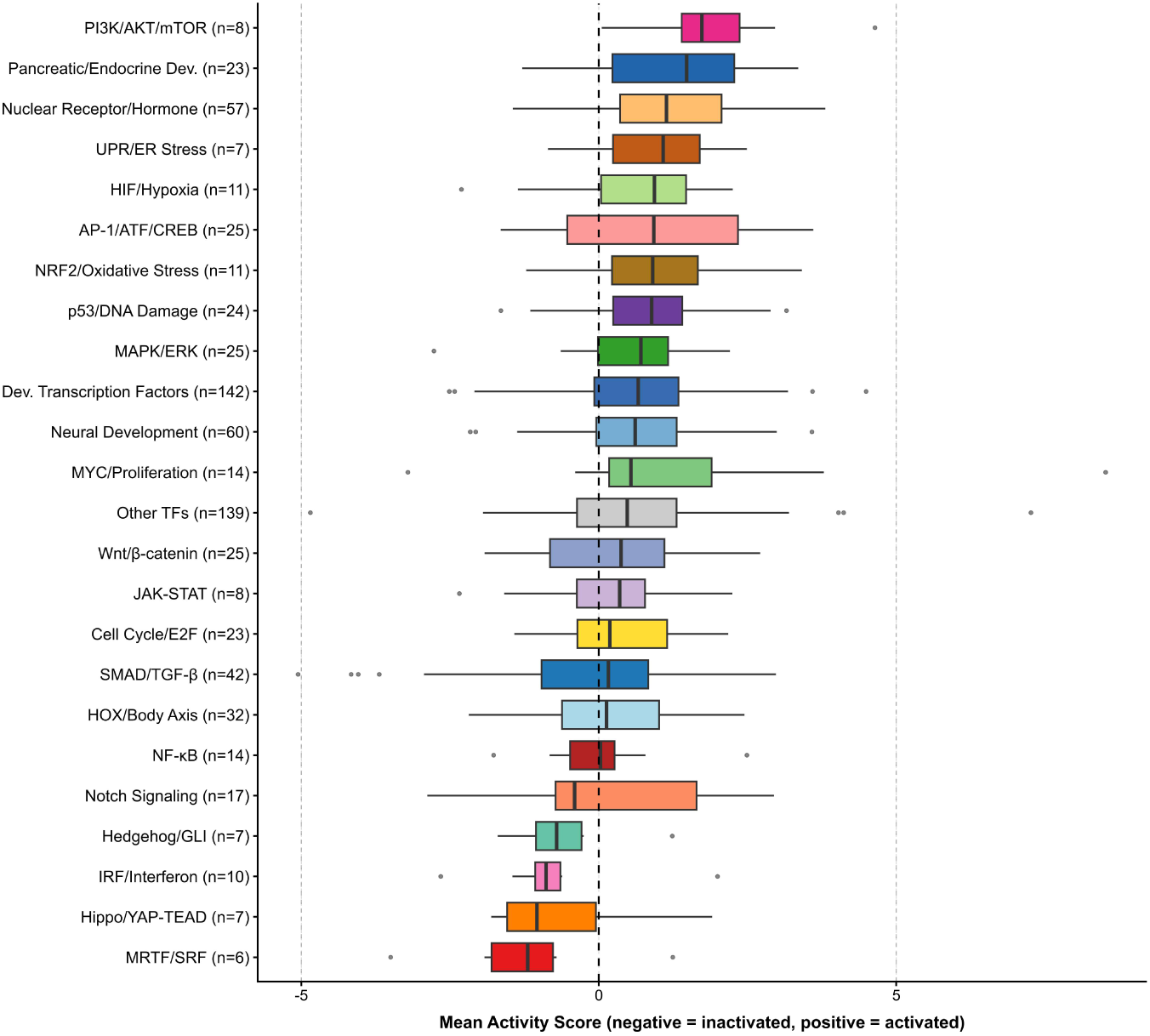
Transcription factor activity organized by signaling pathway and developmental program following Latrunculin B treatment. TF activity scores (mean across 4 timepoints) were inferred using decoupleR (ULM method) with the CollecTRI network. TFs were classified into 24 functional categories using a two-layer approach: (1) gene-name pattern matching for canonical pathway members, followed by (2) GO Biological Process annotation via org.Hs.eg.db for remaining TFs, categories with fewer than 4 members were merged into “Other TFs.” Categories ordered by median activity score, boxplots show median, interquartile range, and 1.5× IQR whiskers. Dashed lines indicate activity score of 0 (no change) and ±5 (strong activation/inactivation). n is number of TFs.

This decoupleR analysis identified MRTF/SRF as the TF network with the most negative median activity score, consistent with G-actin forming a complex with MRTF’s RPEL domain, blocking access to its nuclear localization signal and preventing importin-mediated nuclear entry,(Mouilleron, Langer, Guettler, McDonald, & Treisman, 2011; Pawłowski, Rajakylä, Vartiainen, & Treisman, 2010). Although MRTF cofactors (MKL1/MKL2) are not represented in the TF-target database, the negative SRF activity score serves as a readout of pathway disruption. Of the 107 SRF target genes in the CollecTRI network, 29 showed downregulation, including CCN2, ACTC1, ACTA1, CNN1, and TAGLN. The Hippo/YAP-TEAD pathway had the second most negative median activity score, with TEAD1 showing a mean activity score of −1.38, consistent with YAP/TAZ cytoplasmic retention upon reduced cytoskeletal tension. Canonical YAP/TAZ targets ANKRD1 (log₂FC = −4.26) and CCN2/CTGF (log₂FC = −3.99) were among the most strongly downregulated genes, with CCN2 also a known MRTF-SRF target, consistent with dual regulation by both MRTF-SRF and Hippo/YAP-TEAD pathways(Foster, Gualdrini, & Treisman, 2017).

Several TFs critical for pancreatic endocrine differentiation showed increased activity scores, including NEUROG3 (+3.35), HNF4A (+3.21), and SOX4 (+4.49), as expected from the activation of the endocrine differentiation program(Tian et al., 2025). The analysis also showed increased activity scores of stress response TFs including ATF5 (+3.42), NFE2L2/Nrf2 (+3.42), and CREB1 (+3.61), consistent with engagement of stress and antioxidant response pathways.

The decoupleR analysis also showed very negative activity scores for SMAD/TGF-β signaling, with SMAD9 (−5.05), SMAD5 (−4.16), SMAD4 (−4.04), and SMAD1 (−3.69) representing four of the top six most negatively scored TFs. MYC/Proliferation category showed the widest spread of activity scores across its members with MYC exhibiting the highest mean activation score (+8.52), alongside stress-responsive TFs including NR4A1 and BHLHE40. The chromatin corepressor TRIM28/KAP1 (in Other) ranked second among the most negatively scored TFs (mean score −4.85).

We also performed ChEA3(Keenan et al., 2019) analysis, a separate analysis complementary to decoupleR, that provides a ranked list of TFs associated with the top upregulated and downregulated genes (Supplementary Data File 12). We used the top 300 upregulated and downregulated genes in response to LatB as input to ChEA3. The resulting top four TFs for upregulated genes (FEV, EGR4, NEUROG3, INSM1) are also present in the CollecTRI network with positive decoupleR activity scores, spanning the Neural Development (FEV, INSM1), Pancreatic/Endocrine Development (NEUROG3), and MAPK/ERK (EGR4) categories (Figure 8). Among the top four TFs for downregulated genes (TWIST2, ATOH8, MKX, FOXS1), TWIST2 and MKX had weak negative scores in the decoupleR analysis. TWIST2 regulates smooth muscle, connective tissue, and dermal progenitor gene programs(Li, Cserjesi, & Olson, 1995) while MKX (Mohawk) is critical for tendon differentiation, regulating type I collagen and extracellular matrix gene expression(Ito et al., 2010). ATOH8 and FOXS1 were not part of the CollecTRI network. ATOH8 is a direct target of NEUROG3 expressed in both endocrine and exocrine pancreatic precursor cells(Lynn, Sanchez, Gomis, German, & Gasa, 2008). FOXS1 is an early marker of the sensory nervous system expressed in neural crest-derived sensory neurons(Montelius et al., 2007). The ChEA3 analysis(Keenan et al., 2019) provided additional support for Hippo/YAP-TEAD pathway involvement by TEAD3 at rank 11 of downregulated genes (absent from CollecTRI network). SRF itself ranked 265th in the ChEA3 downregulated analysis despite having a strong negative decoupleR score.

### Gene expression Changes in MAPK Pathways

LatA treatment in fission yeast activates Sty1(Mutavchiev, Leda, & Sawin, 2016), the p38 MAPK homolog, and LatB treatment of HMC-1 cells activates p38 and ERK1/2 (Meyer et al., 2007). We thus examined expression changes of core MAPK pathway components over time. Figure 9 shows the per-timepoint expression of core MAPK pathway components (Supplementary Data File 13), with branch annotations assigned based on established MAPK cascade biochemistry.

**Figure 9.**
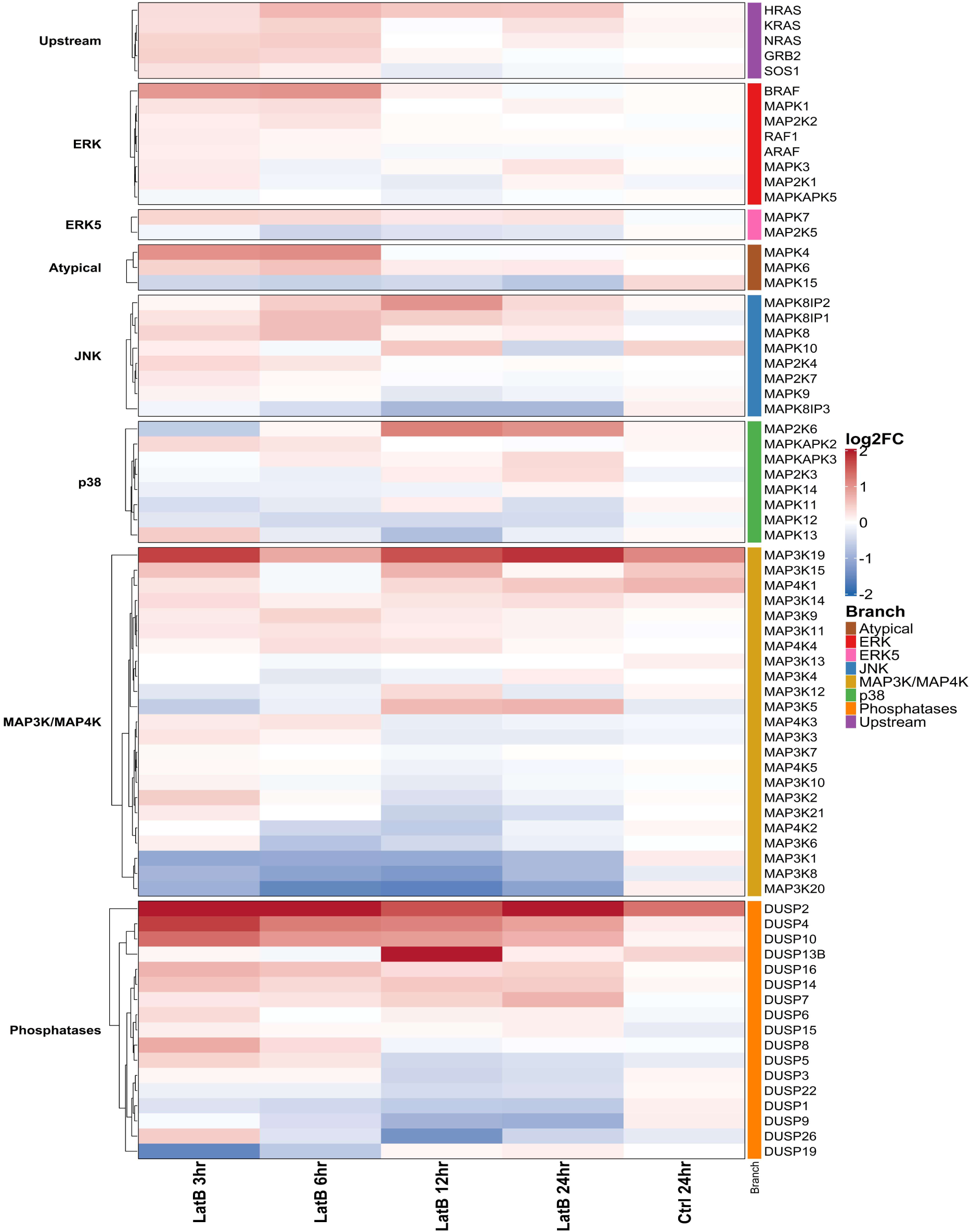
MAPK pathway kinase and phosphatase expression dynamics. Log₂ fold changes for core MAPK components retrieved from MSigDB (KEGG MAPK pathways, GO: BP MAPK-related terms). Branch annotations (Upstream, ERK, ERK5, Atypical, JNK, p38, MAP3K/MAP4K, Phosphatases) assigned based on gene nomenclature and established MAPK cascade biochemistry. Rows clustered hierarchically within each branch (Euclidean distance, complete linkage). Ctrl 24hr represents untreated cells at 24 hours versus Control 0hr baseline.

The mammalian p38 MAPK genes (MAPK14, MAPK11, MAPK12, MAPK13) showed no substantial transcriptional changes. Sty1 activation occurs via phosphorylation rather than transcriptional upregulation(Mutavchiev et al., 2016), a process RNA-seq cannot detect. Whether similar phosphorylation of p38 MAPKs occurs in human pancreatic progenitors following actin disruption remains to be determined experimentally.

Our analysis did, however, reveal some notable gene expression changes. The dual-specificity phosphatases DUSP4 and DUSP10 showed strong sustained upregulation across all timepoints relative to control, indicating negative transcriptional feedback within MAPK signaling. DUSP4 acts preferentially on ERK, whereas DUSP10 targets the p38 and JNK branches(Patterson, Brummer, O’Brien, & Daly, 2009). Conversely, several MAP3K family members (MAP3K1, MAP3K8, MAP3K20) were progressively downregulated compared to control, suggesting reduced expression of specific MAPK signaling components.

## Discussion

Prior work used Latrunculin treatments to link actin state to pancreatic endocrine fate. Hogrebe et al. showed by bulk and single-cell RNA sequencing that actin depolymerization by LatA promotes NEUROG3-dependent endocrine specification in human pluripotent stem cell (hPSC)-derived pancreatic progenitor cells(Hogrebe et al., 2020). Tian et al. identified salt-inducible kinases as the F-actin-to-YAP1 transducers driving pancreatic progenitors toward the NEUROG3⁺ endocrine lineage after LatB treatment (Tian et al., 2025). More recent work used bulk and single cell RNA-seq to monitor the exit from pluripotency response of hPSCs treated by LatA during the first 24 h of definitive endoderm formation (Hogrebe et al., 2026). Consistent with these results, our reanalysis confirms progressive activation of the endocrine differentiation program across the 24-hour time course (Supplementary Figures 2, 3, 4, Supplementary Data File 8).

In addition to the earlier findings on endocrine differentiation, our analysis shows that LatB treatment induced a multi-layered response encompassing cytoskeletal transcriptional feedback, with strong negative inferred activity for MRTF/SRF and YAP/TAZ-TEAD pathways, alongside increased activity scores of stress-responsive TFs. Although LatB directly targets mainly actin, transcriptional changes extended across all three cytoskeletal systems. We note that this analysis does not resolve how these cytoskeletal expression changes relate specifically to the endocrine differentiation program versus general mechanisms of cytoskeletal homeostasis; for example, the coordinated suppression of keratins alongside induction of neurofilament genes (notably NEFL) might reflect intermediate filament identity as part of the differentiation response.

LatB treatment, which increases the monomeric actin pool, was accompanied by downregulation of both ACTB and ACTG1 and a strong negative SRF activity score (Figures 2 and 8). In Actb-knockout fibroblasts, several SRF target genes, including γ-actin and α-smooth-muscle actin were upregulated(Bunnell, Burbach, Shimizu, & Ervasti, 2011). Genetic perturbation of the Actb locus can also be accompanied by upregulation of other actin isoforms, including Actg1 (Vedula et al., 2017). Together, these observations support an association between monomeric actin availability and SRF transcriptional output, consistent with the G-actin–MRTF/SRF feedback(Sotiropoulos et al., 1999).

Because many mammalian protein half-lives are typically on the order of days, including actin, profilin, and cofilin (Schwanhäusser et al., 2011), the transcriptional changes observed here within hours (Figures 2, 3) may alter protein levels only beyond the 24-h window analyzed. Prior work in *Chlamydomonas reinhardtii* has shown the existence of a specific Latrunculin B response program where the Latrunculin-sensitive actin IDA5 is replaced by the Latrunculin-resistant actin NAP1 within ∼ 2 hrs,(Onishi, Pecani, Jones, Pringle, & Cross, 2018; Onishi, Pringle, & Cross, 2016). Interestingly, the increase in cofilin and profilin reported in *Chlamydomonas* echoes the mild increase in PFN1 (Figure 2). The exchange of actin in this organism is enabled by the upregulation of genes encoding both NAP1 and proteins of the ubiquitin-proteasome system that degrade IDA5,(Onishi et al., 2018, 2016). While this NAP1-based compensation is specific to *Chlamydomonas*, it raises the question of whether the ubiquitin-proteasome system might be engaged in degradation of excess G-actin and other cytoskeletal proteins in human cells, as a mechanism of quick adaptation of cytoskeletal protein concentrations. We do see changes in multiple genes related to protein homeostasis (Figure 7). Several proteasome subunit genes are modestly upregulated (for example PSMB2, PSMC1, and PSMD3), as is the polyubiquitin gene UBB. However, these changes are small in magnitude and some ubiquitin ligases known to target cytoskeletal substrates show mixed regulation: TRIM3 which targets γ-actin (Cho & Kim, 2026), ASB2 which targets filamins A and B (Heuzé et al., 2008) and TRIM63/MuRF1, all decrease in expression, whereas the atrophy-associated ligase FBXO32/atrogin-1 is modestly upregulated (Bodine et al., 2001). Whether the ubiquitin-proteasome system is actively engaged in adapting cytoskeletal protein concentrations therefore remains an open question for future work.

More broadly, transcriptomic profiling of cells following cytoskeletal-drug treatment can uncover feedback between the cytoskeleton and gene expression. It is interesting to compare the transcriptional response of the microtubule system to indirect perturbation by LatB (Figure 4) with its response to direct microtubule targeting drugs,(Gasic, Boswell, & Mitchison, 2019; Horin, Sonnett, Li, & Mitchison, 2025). The LatB effects on key microtubule genes were concentrated on motors and MAPs: the kinesin KIF19 was the most strongly upregulated microtubule gene (log₂FC = +2.36), the microtubule-associated proteins MAP1A and MAP1B were induced (log₂FC +0.87 and +0.71), and several other kinesins were downregulated (for example log₂FC for KIFC3 and KIF12 were −1.18 and −0.68) (Supplementary Data File 5). This pattern differs from the response to direct microtubule drugs in cultured cells studied by Gasic et al.(Gasic et al., 2019) who showed that microtubule-targeting agents, including the stabilizer paclitaxel and the destabilizer combretastatin A-4, drive coordinated upregulation and downregulation of α-, β-, and γ-tubulin isoforms through regulated tubulin mRNA stability. Notably, this response was specific to tubulins and did not extend to microtubule motors, MAPs, or plus-tip proteins (Gasic et al., 2019). In the response to LatB analyzed here, the abundant α- and β-tubulin isotypes were largely unchanged and the larger tubulin fold changes visible in Figure 4 (TUBB8, TUBA8, TUBB1) correspond to minimally expressed isotypes. Thus, noting the difference between the two experimental setups, the transcriptional response of the microtubule system to LatB appears to be distinct from that of direct microtubule drugs.

The broader strategy of coupling cytoskeletal perturbation with transcriptomic profiling may prove valuable for dissecting developmental morphogenesis, where coordinated cytoskeletal remodeling generates the differential cell properties that drive tissue folding and shape change in tight coupling with underlying gene-expression programs. Tractable cnidarian models such as the sea anemone *Nematostella vectensis*, in which embryonic patterning and morphogenesis are readily accessible to manipulation (Layden, Rentzsch, & Röttinger, 2016), offer a promising context for this approach. Pharmacological perturbation has already been used productively in *Nematostella* to dissect the signaling basis of early development (Layden, Johnston, et al., 2016), raising the possibility that latrunculin B could be applied the same way in developing tissues to disentangle how mechanical force and gene expression jointly shape morphogenesis.

## Supporting information

Scripts

## Acknowledgments

This work was supported by the National Institutes of Health (NIH) grant R35 GM136372 to D.V. and the Lehigh College of Arts and Sciences Dean’s Opportunity Grant (to M.J.L. and D.V.).

## Code availability

The analysis scripts used to generate the figures in this study are provided as supplementary code.

## Author contributions

G.K. performed the analysis and prepared the figures. D.V. supervised the study. G.K., M.J.L. and D.V. interpreted the results and wrote the manuscript.

