## Supplementary Material for "Analysis of gene expression changes after Latrunculin B-induced actin depolymerization"

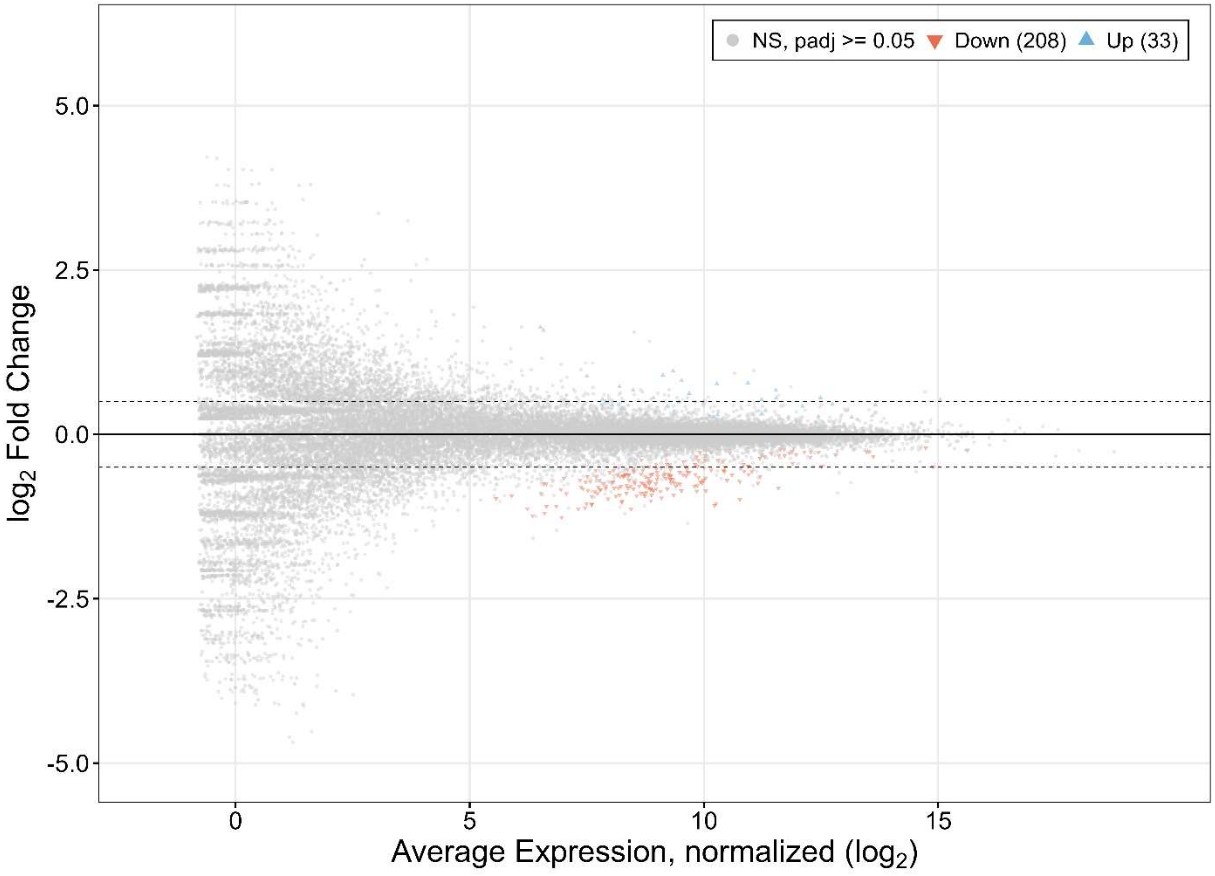


**Supplementary Figure 1: MA plot comparing the change of the Control sample at 24 hr with respect to Control 0hr.** Each point represents a gene plotted by average expression, normalized (x-axis) versus $log_2$ fold change (y-axis). Salmon: P-significantly downregulated (n = 208), blue: P-significantly upregulated (n = 33), grey: not P-significant.

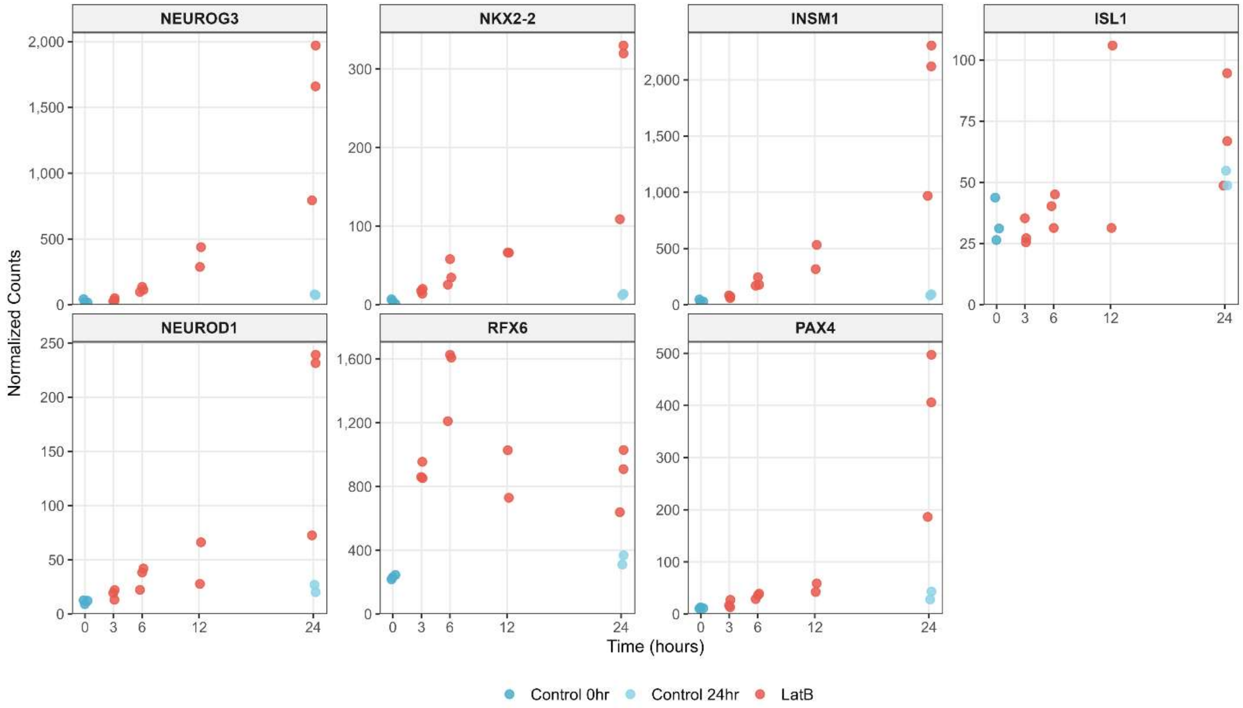


**Supplementary Figure 2: Expression of key pancreatic endocrine differentiation markers following Latrunculin B treatment.** DESeq2 size factor-normalized counts for NEUROG3, NKX2-2, INSM1, ISL1, NEUROD1, RFX6, and PAX4 across the LatB treatment time course. All seven genes show upregulation consistent with activation of the pancreatic endocrine differentiation program described by Tian et al. *Stem Cell Reports* **20**, 102444 (2025).

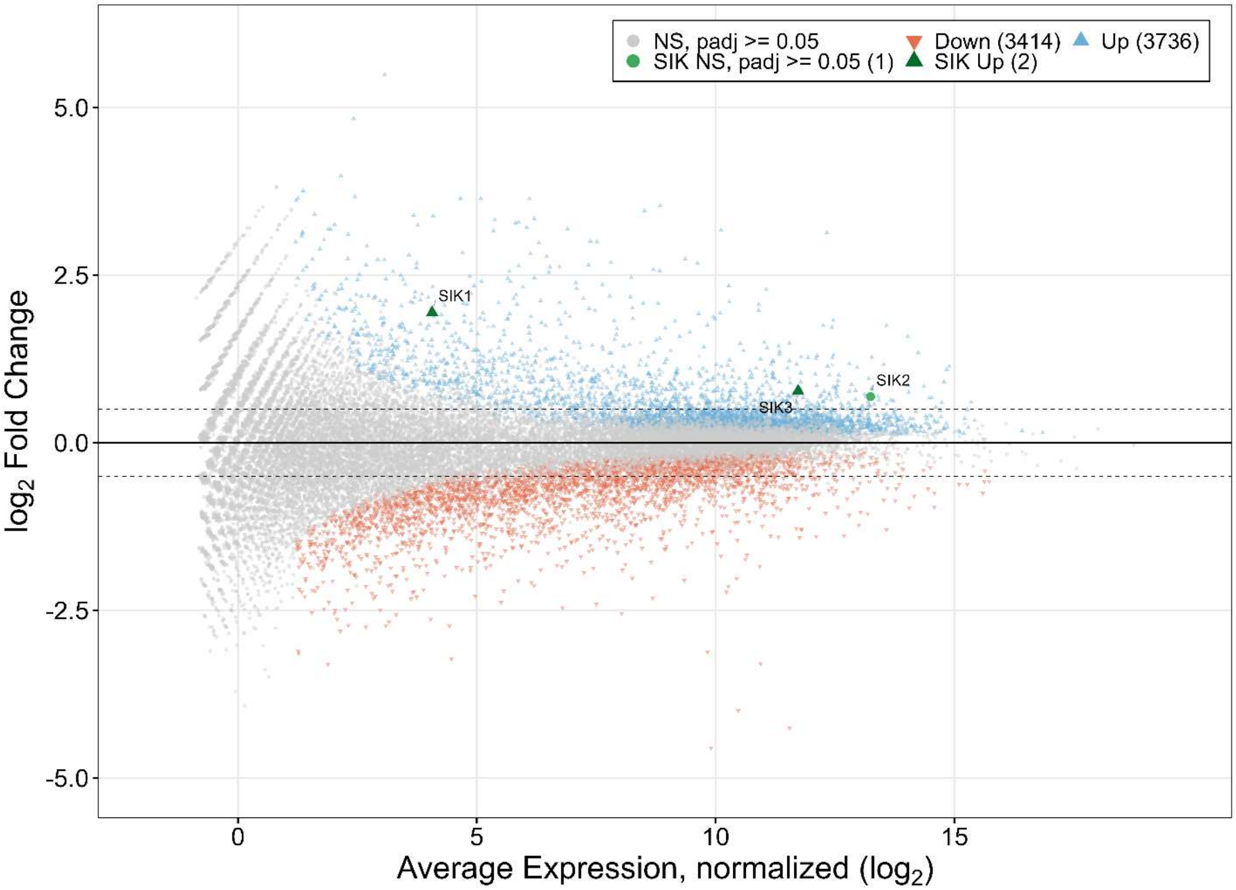


**Supplementary Figure 3: MA-plot as in Figure 1, identifying salt-inducible kinase (SIK) family members following Latrunculin B treatment.** Each point represents a gene plotted by average expression, normalized (x-axis) versus $\log_2$ fold change (y-axis), with dashed lines at $\log_2$ fold change of ±0.5. Grey: not significant, blue: P-significantly downregulated, salmon: P-significantly upregulated (padj < 0.05).

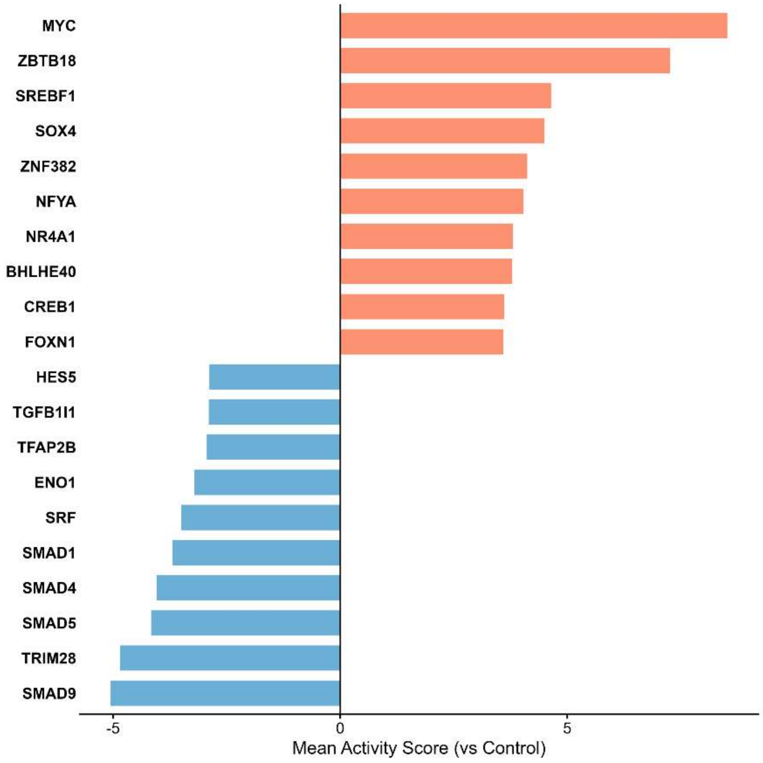


**Supplementary Figure 4: Transcription factor activity changes following Latrunculin B treatment.** Mean activity scores for the top 10 positive activity scores (salmon) and top 10 negative inactivity scores (blue) transcription factors, computed using decoupleR with the CollecTRI network. SMAD family members (SMAD9, SMAD5, SMAD4, SMAD1), the chromatin corepressor TRIM28, and SRF show strong negative scores. MYC exhibits the highest activity score, alongside metabolic regulators (SREBF1), stress-responsive factors (NR4A1, BHLHE40), and developmental TFs (SOX4, ZBTB18

**Supplementary Data Files.**

**Supplementary Data File 1.** Genome-wide raw count matrix with per-timepoint statistics. Integer raw (unnormalized) read counts for all 28,058 genes across the 16 samples. Includes gene annotation, the 16 raw count columns labeled by time point (log$_2$FC, padj, baseMean) for the six contrasts. See correspondence to SRA run below.

**Supplementary Data File 2.** Per-timepoint DESeq2 comparison tables. A folder of six CSV files with the full DESeq2 results for the individual pairwise contrasts resolving the LatB response over time. Each reports all 28,058 genes (Ensembl gene ID plus baseMean, log2FoldChange, lfcSE, stat, pvalue, padj). The six contrasts are Control 24 hr vs Control 0 hr, LatB 3/6/12/24 hr each vs Control 0 hr, and LatB 24 hr vs Control 24 hr. This file contains overlapping information with data tables of Supplementary Data File 8.

**Supplementary Data File 3.** Gene-level Gene Ontology annotations. The gene-to-GO-term mapping used for functional categorization, in long format. Columns give the gene symbol, GO accession, evidence code, ontology domain (Biological Process, Cellular Component, and Molecular Function), and term name. Only the Biological Process annotations were used for Figure 7.

**Supplementary Data File 4.** Significant differentially expressed genes by functional category (Figure 7). The 6,632 genes significant in at least one LatB timepoint (padj < 0.05 and absolute log$_2$FC > 0.5), each assigned to one of 18 genome-wide functional categories used for Figure 7. Columns give gene annotation, per-timepoint log$_2$FC and padj for the four LatB-versus-Control 0 hr comparisons, per-timepoint significance flags, and the final category label.

**Supplementary Data File 5.** Genome-wide pooled differential expression. DESeq2 size factor-normalized counts and pooled differential expression statistics for all 28,058 genes. Columns include gene annotation (SYMBOL, ensembl_id, description, gene_biotype), normalized counts for the 16 samples (LatB 3/6/12/24 hr and Control 0/24 hr), and the pooled LatB-versus-control statistics (log$_2$FC, padj, baseMean, pvalue, lfcSE, and Wald stat).

**Supplementary Data File 6.** Actin cytoskeleton expression summary (pooled). Pooled LatB-versus-control differential expression for 159 actin cytoskeleton genes, supporting Figure 1. Each row gives the gene symbol, log2FC, adjusted p-value, direction, and a PFC-significance flag (padj < 0.05 and |log$_2$FC | > 0.5).

**Supplementary Data File 7.** Actin cytoskeleton genes by functional family. 138 actin cytoskeleton genes assigned to 17 functional families, with their pooled LatB-versus-control statistics (log2FC, padj, and a P-significance flag), underlying Figure 1B.

**Supplementary Data File 8.** Genome-wide normalized counts with per-timepoint statistics. This file contains overlapping information with data tables of Supplementary Data File 2. DESeq2 size factor-normalized counts for all 28,058 genes across the 16 samples, plus per-timepoint differential expression statistics (log$_2$FC, padj, baseMean) for the six contrasts. Provides the normalized expression trajectories used for the per-gene time-course analysis.

**Supplementary Data File 9.** Time-course expression plots of cytoskeletal gene families. A folder of pre-rendered plots. Contains per-gene panels showing DESeq2 size-factor normalized expression across the

LatB time course (Control 0/24 hr and LatB 3/6/12/24 hr) for actin, microtubule, intermediate-filament, and coronin gene families.

**Supplementary Data File 10.** MRTF/SRF target gene panel (pooled). 54 MRTF/SRF target and pathway genes with their pooled LatB-versus-control statistics. Columns give the gene symbol, functional category, $log_2FC$, padj, baseMean, biotype, description, and a significance flag. Single pooled statistic per gene, no per-sample counts.

**Supplementary Data File 11.** Transcription factor activity scores (decoupleR). Inferred activity for 737 transcription factors across the LatB time course, with per-timepoint scores (LatB 3/6/12/24 hr), summary statistics, and a multi-layer pathway classification (Final_Pathway plus the layer and source of each assignment). Underlies Figure 8.

**Supplementary Data File 12.** ChEA3 transcription factor enrichment. ChEA3 (Integrated meanRank) TF enrichment results for the top 300 up- and top 300 down-regulated genes from the pooled comparison, provided as a single long-format table with up- and down-regulated results combined. Lists ranked candidate transcription factors with direction, integrated rank, score, contributing source libraries, and overlapping target genes. Complements the decoupleR analysis in Supplementary Data File 11 and Figure 8.

**Supplementary Data File 13.** MAPK pathway genes with branch annotation. The 63 core MAPK pathway genes used for Figure 9, each assigned to one of seven signaling branches. Two columns: gene symbol and branch.

**Sample Mapping for Supplementary Data Files 1, 5, and 8**

SRR30416728, SRR30416729, SRR30416730: Control 0hr (replicates 1–3)

SRR30416726, SRR30416727: Control 24hr (replicates 1–2)

SRR30416723, SRR30416724, SRR30416725: LatB 3hr (replicates 1–3)

SRR30416720, SRR30416721, SRR30416722: LatB 6hr (replicates 1–3)

SRR30416718, SRR30416719: LatB 12hr (replicates 1–2)

SRR30416715, SRR30416716, SRR30416717: LatB 24hr (replicates 1–3)
