## Supplementary figures and images for "Analysis of gene expression changes after Latrunculin B-induced actin depolymerization"

### _Actin_Monomers.png

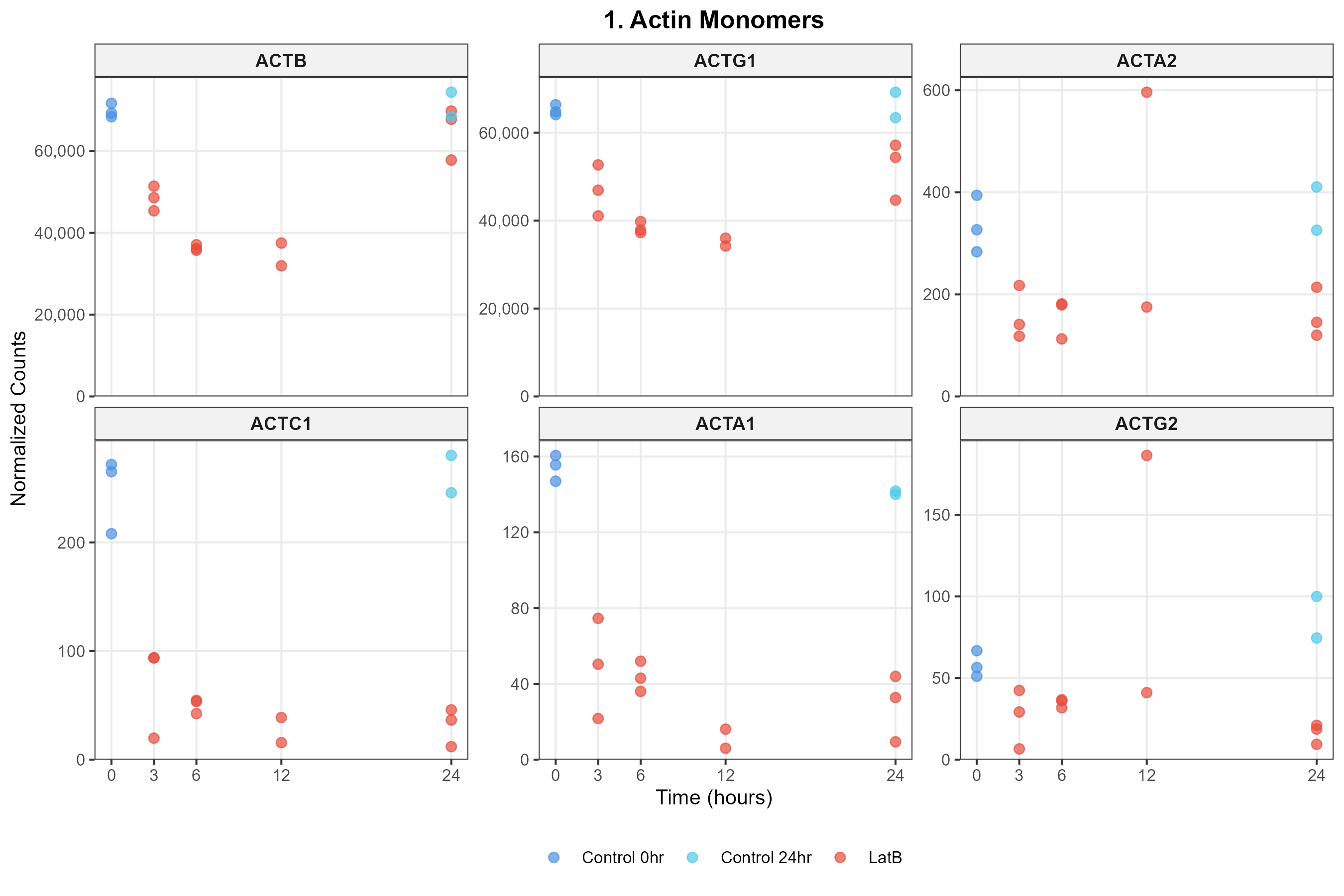

### _Arp2_3_Complex.png

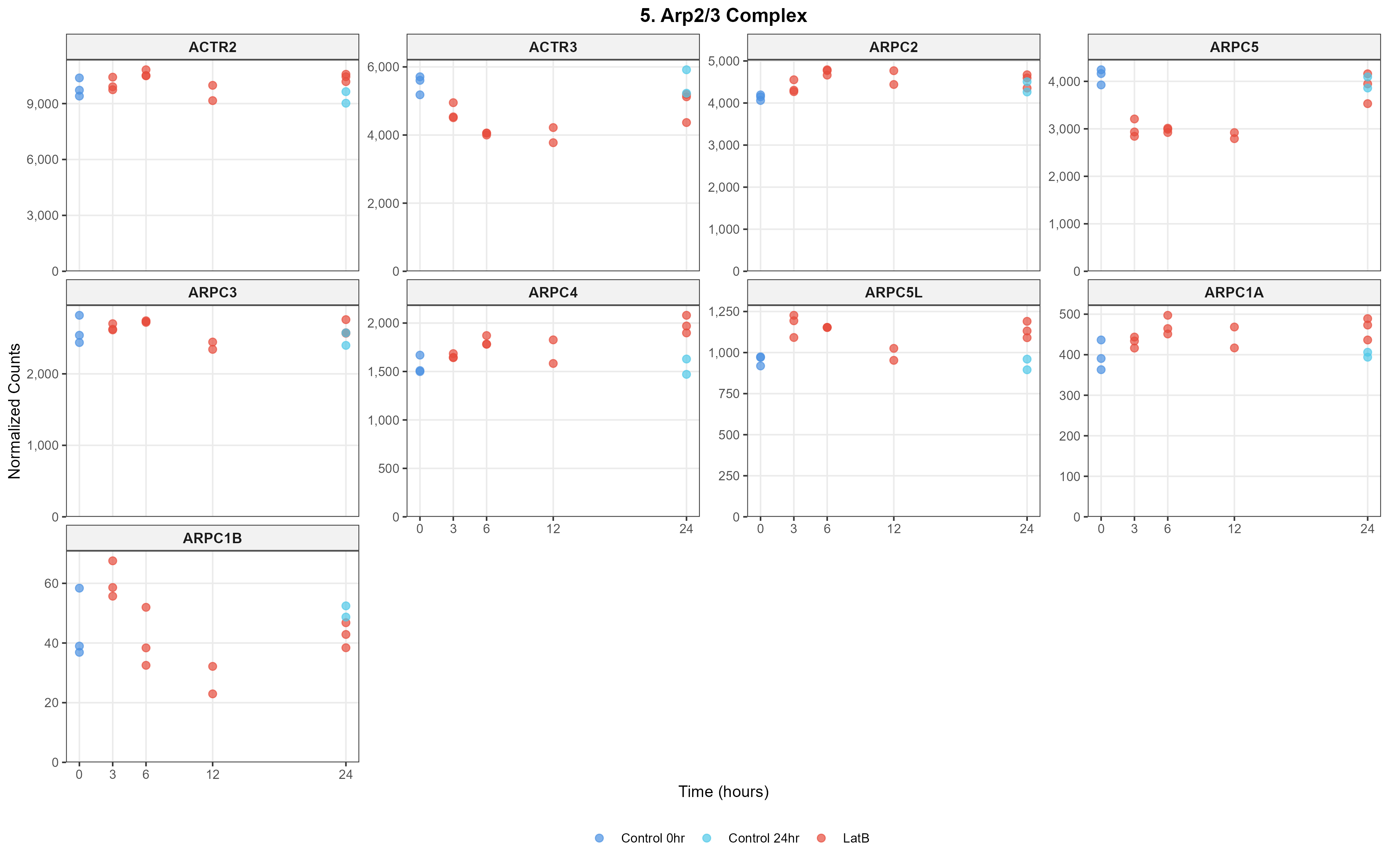

### _Capping___Related.png

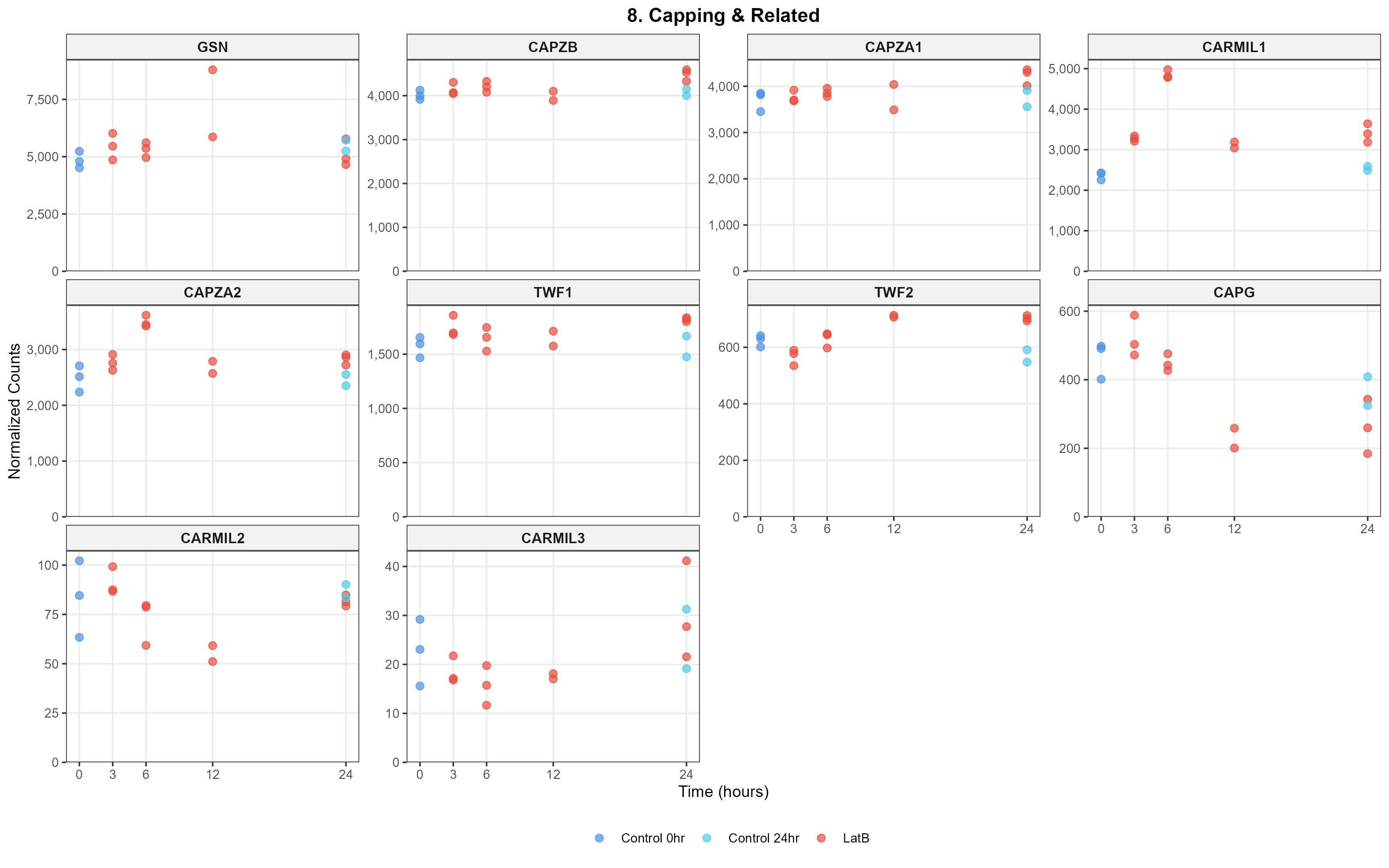

### _Cofilin_ADF.png

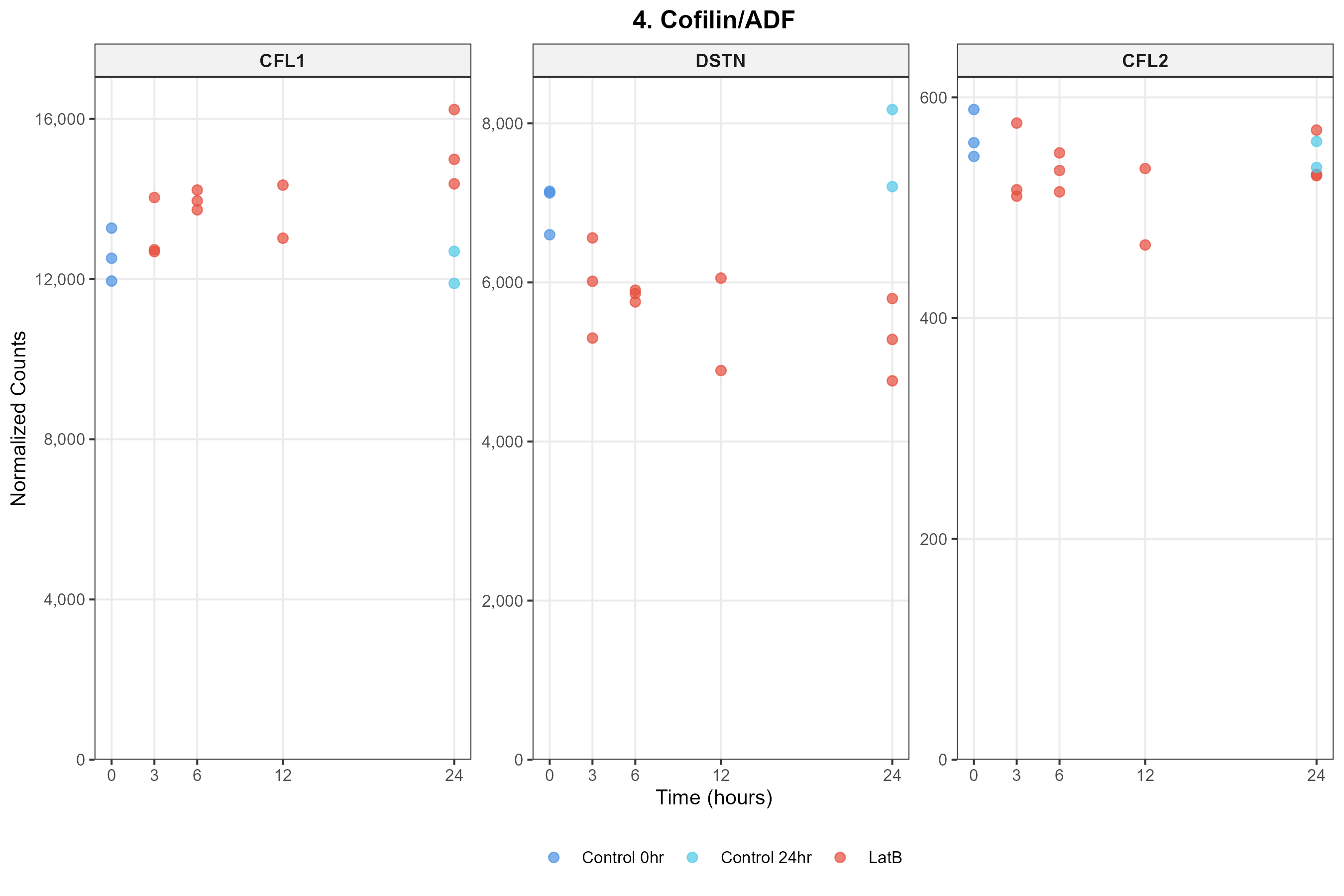

### _Coronins.png

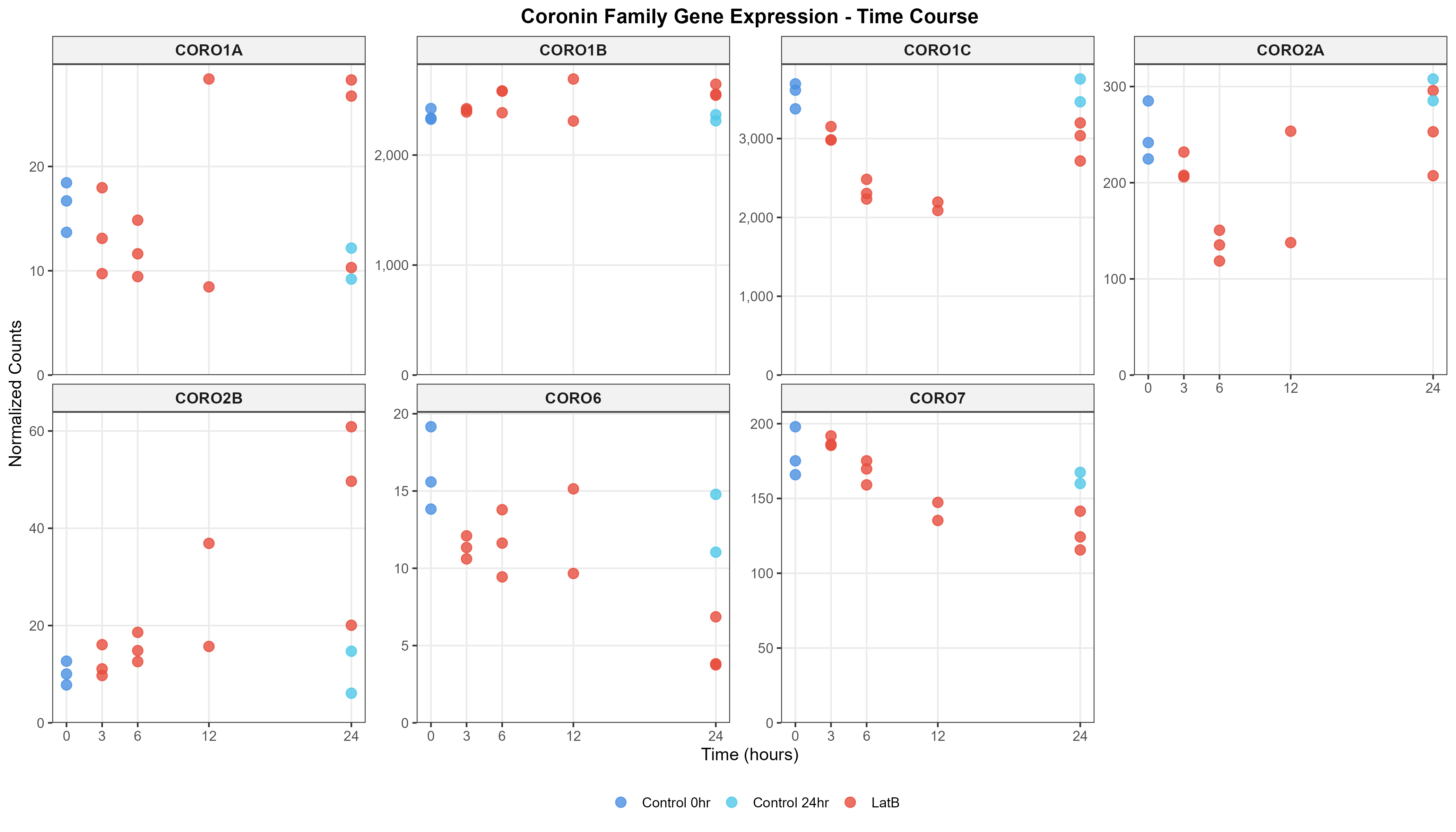

### _Crosslinkers.png

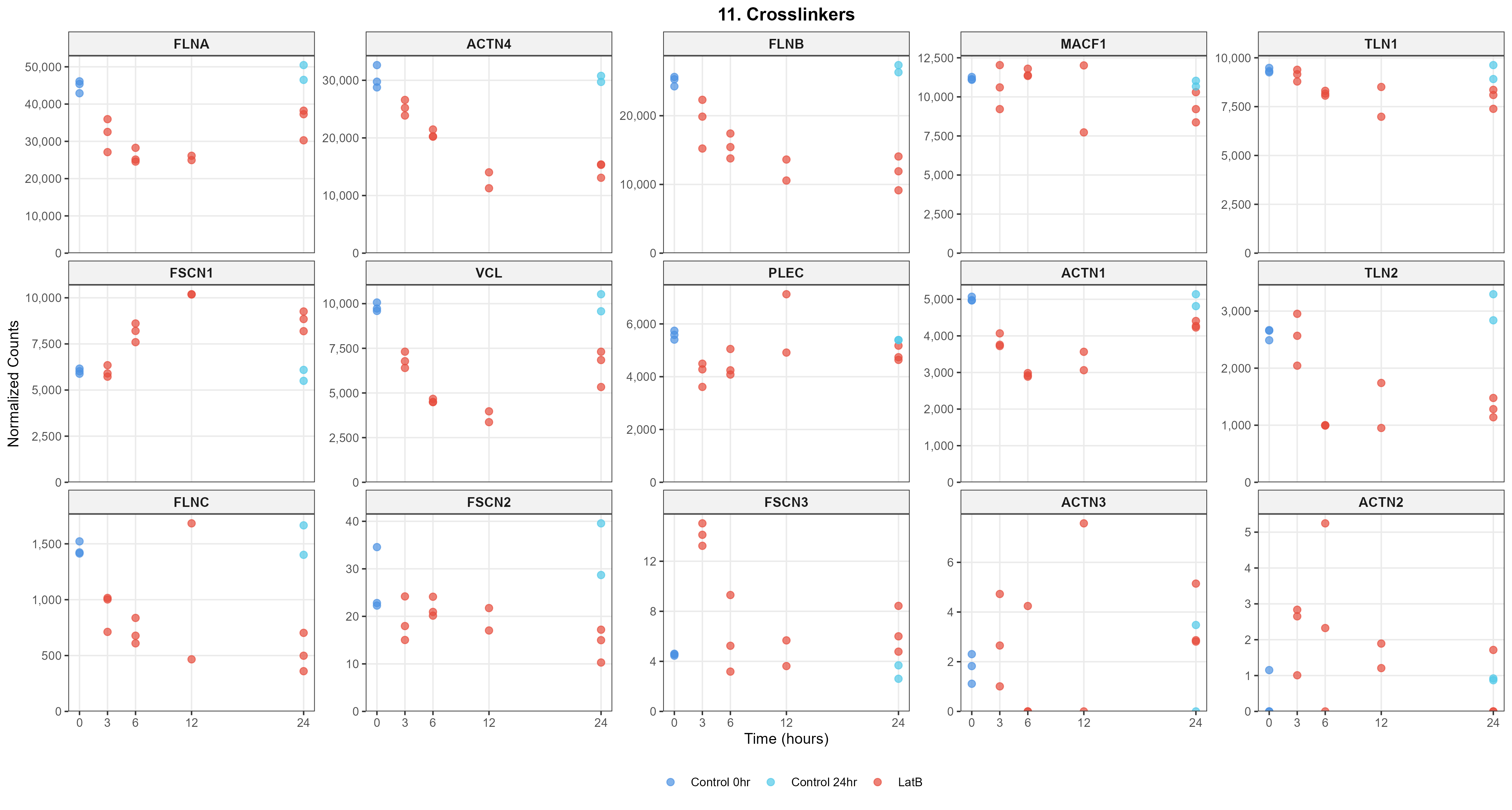

### _Ena_VASP.png

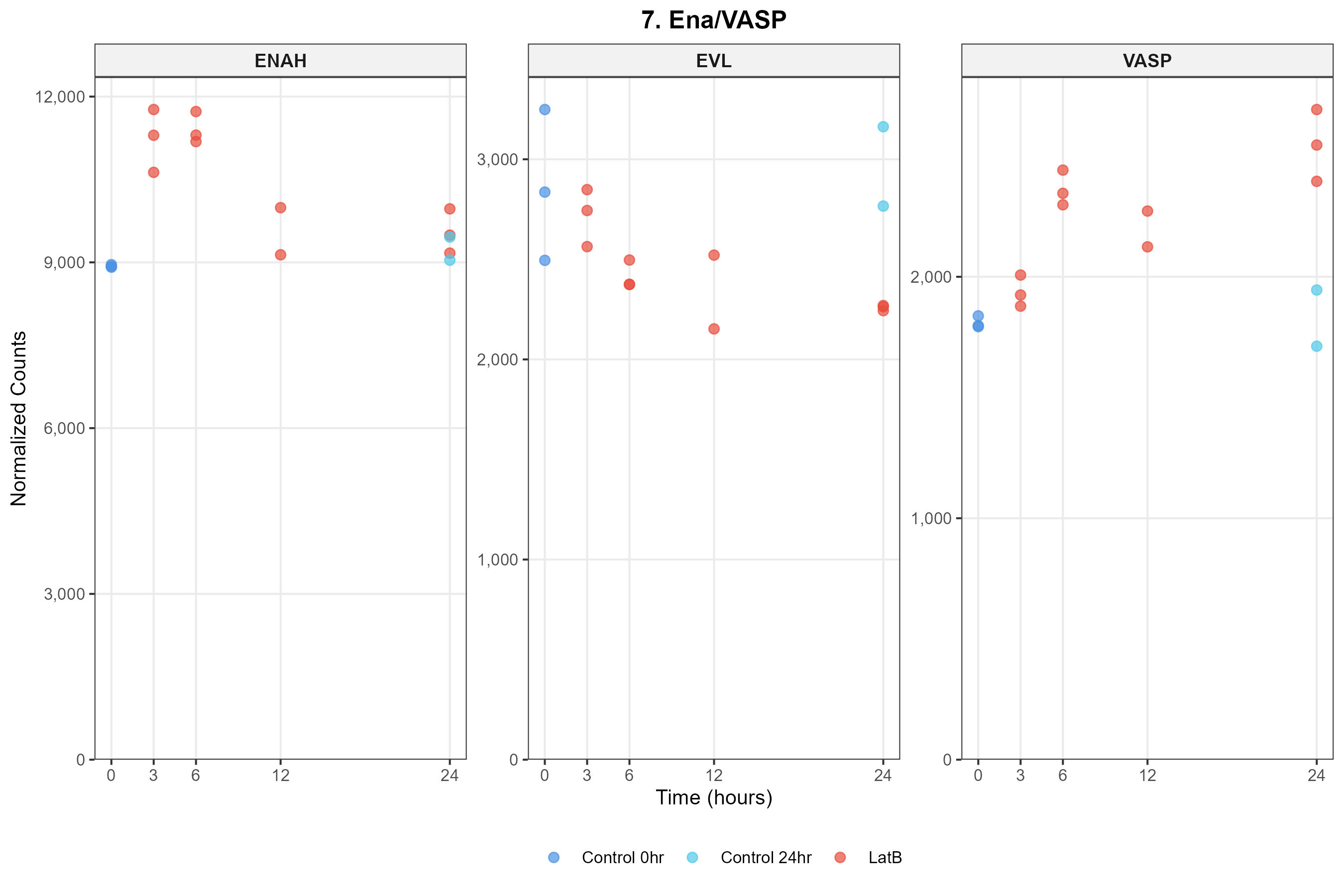

### _Formins.png

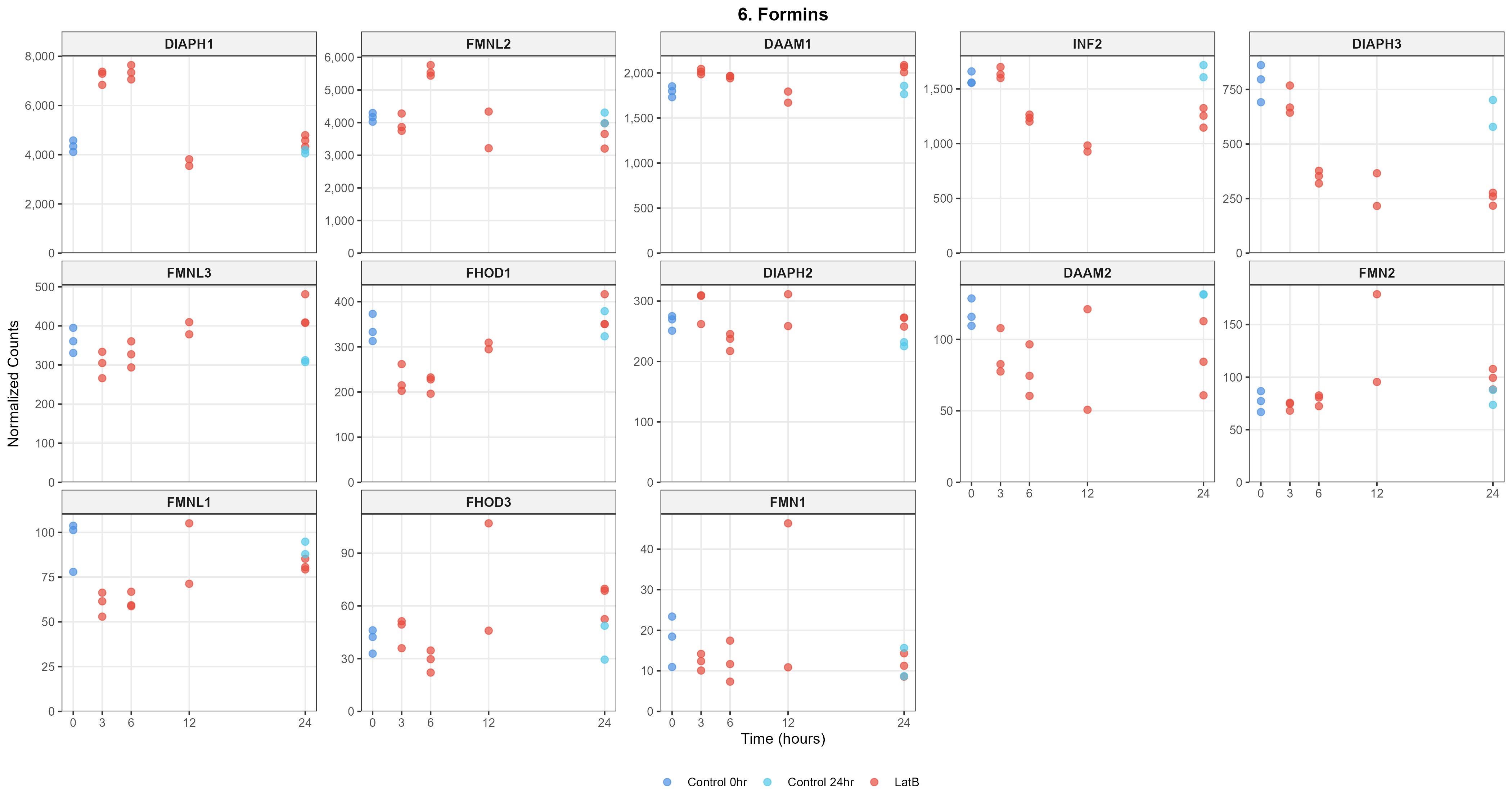

### _Myosin_I.png

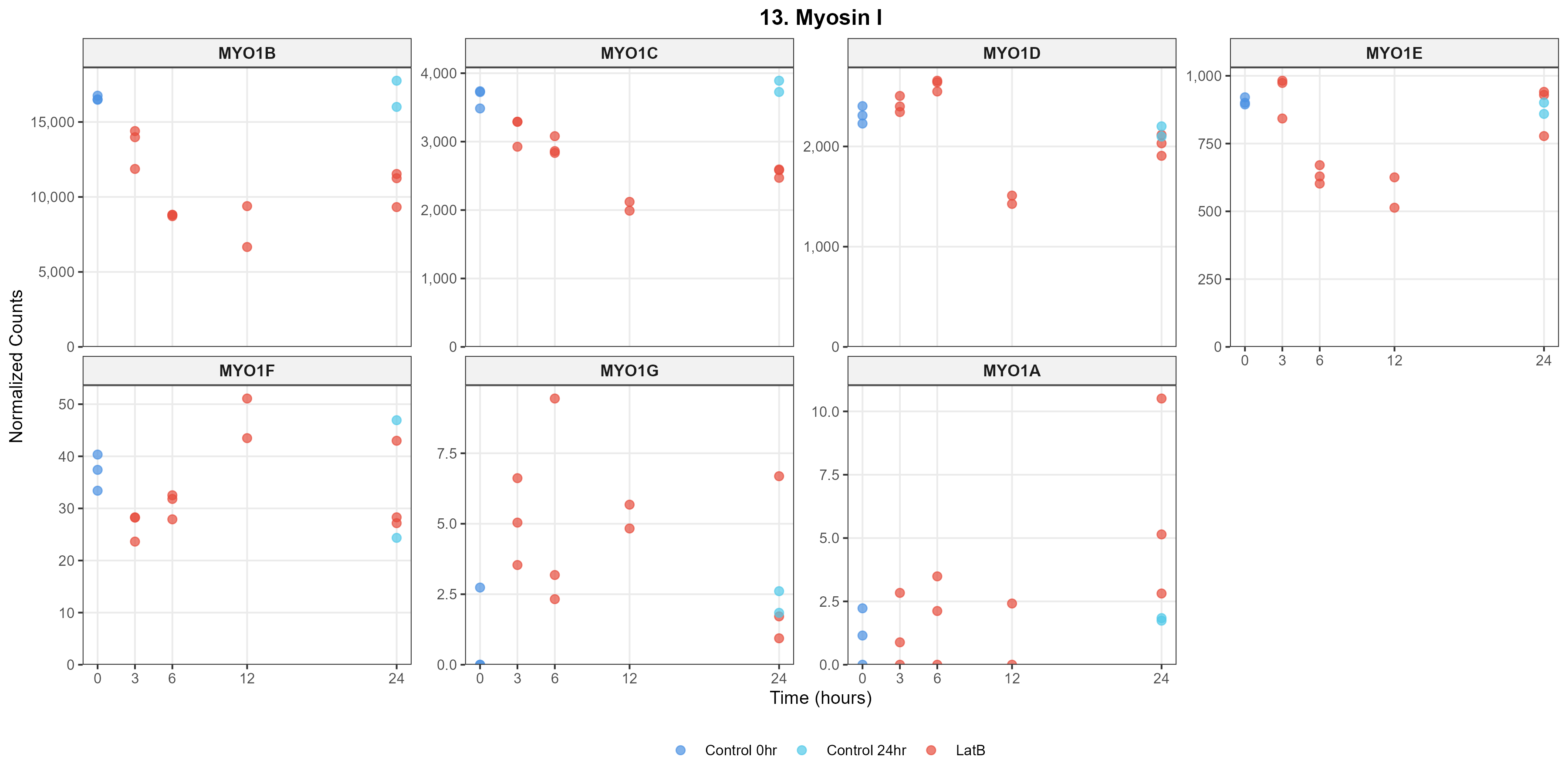

### _Myosin_II.png

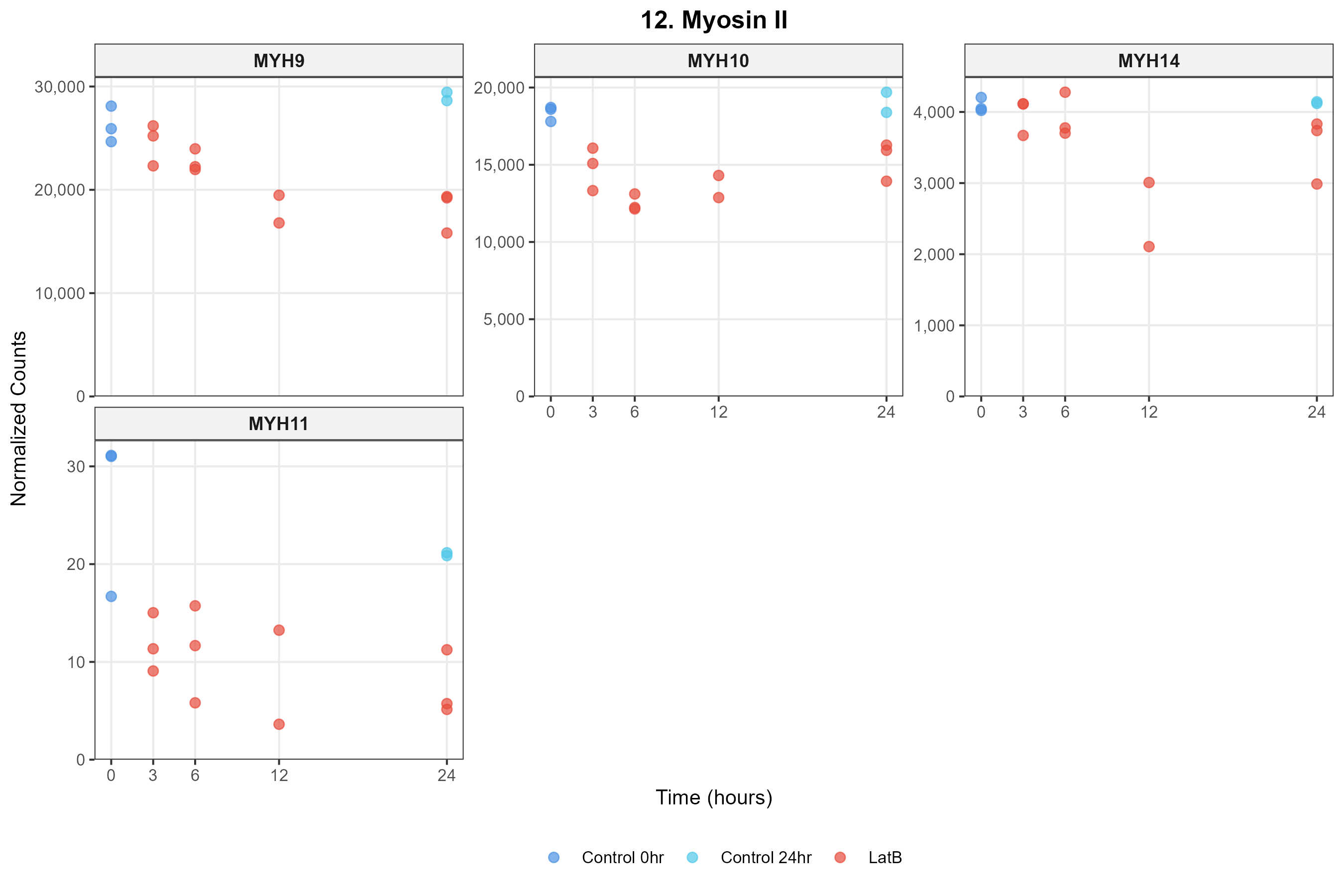

### _Myosin_V_VI_X.png

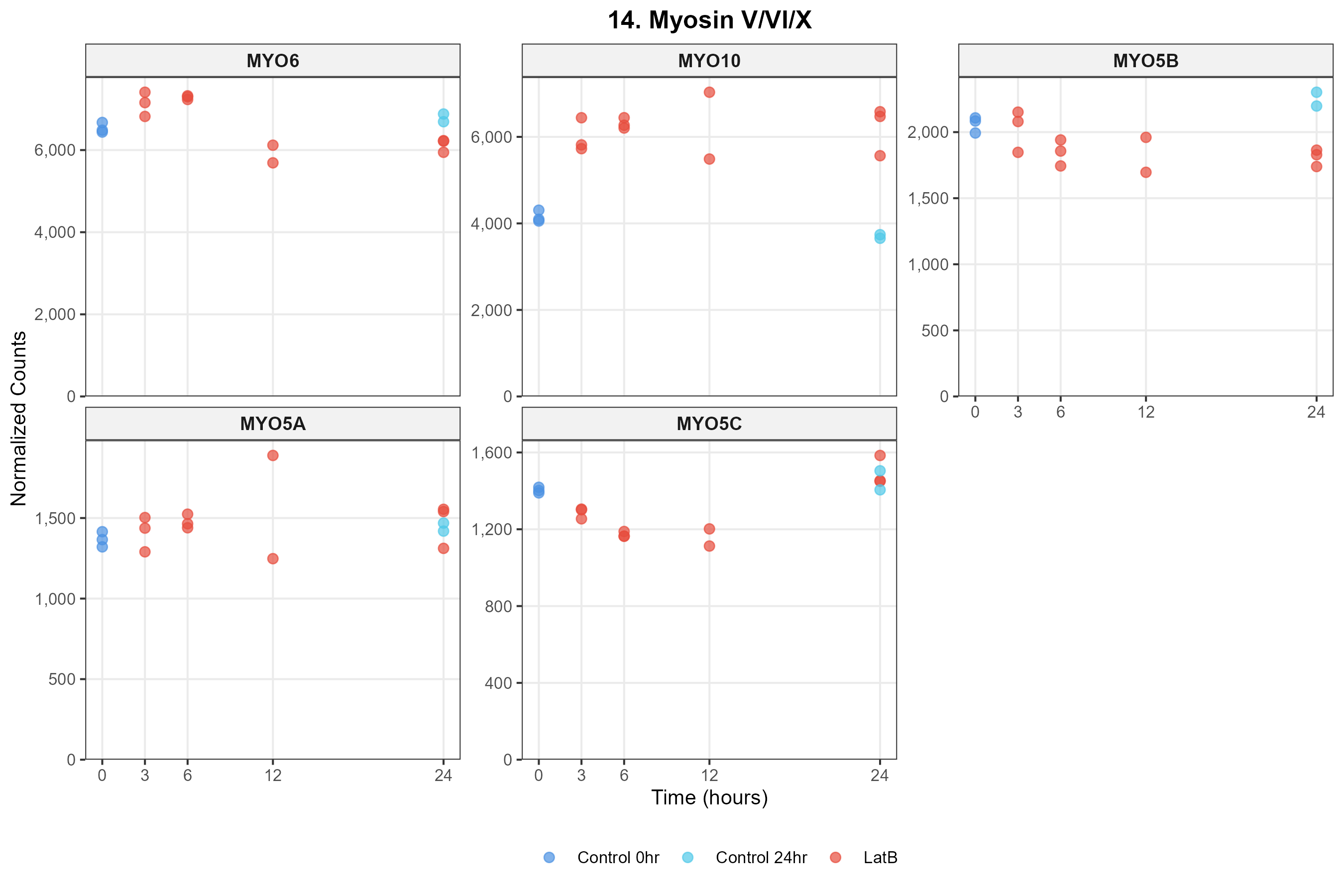

### _NPFs__WASP_WAVE_.png

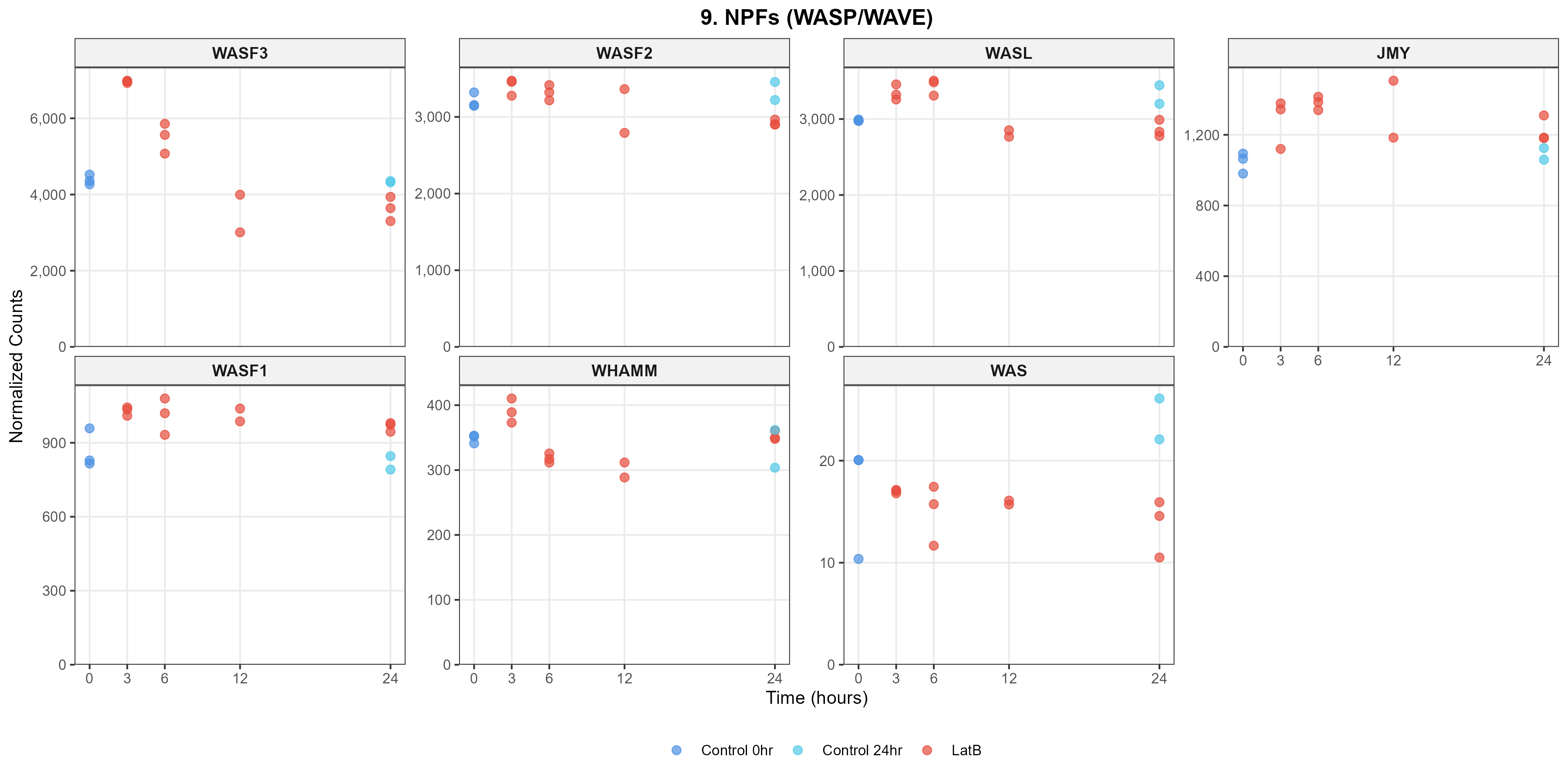

### _Other_Myosins.png

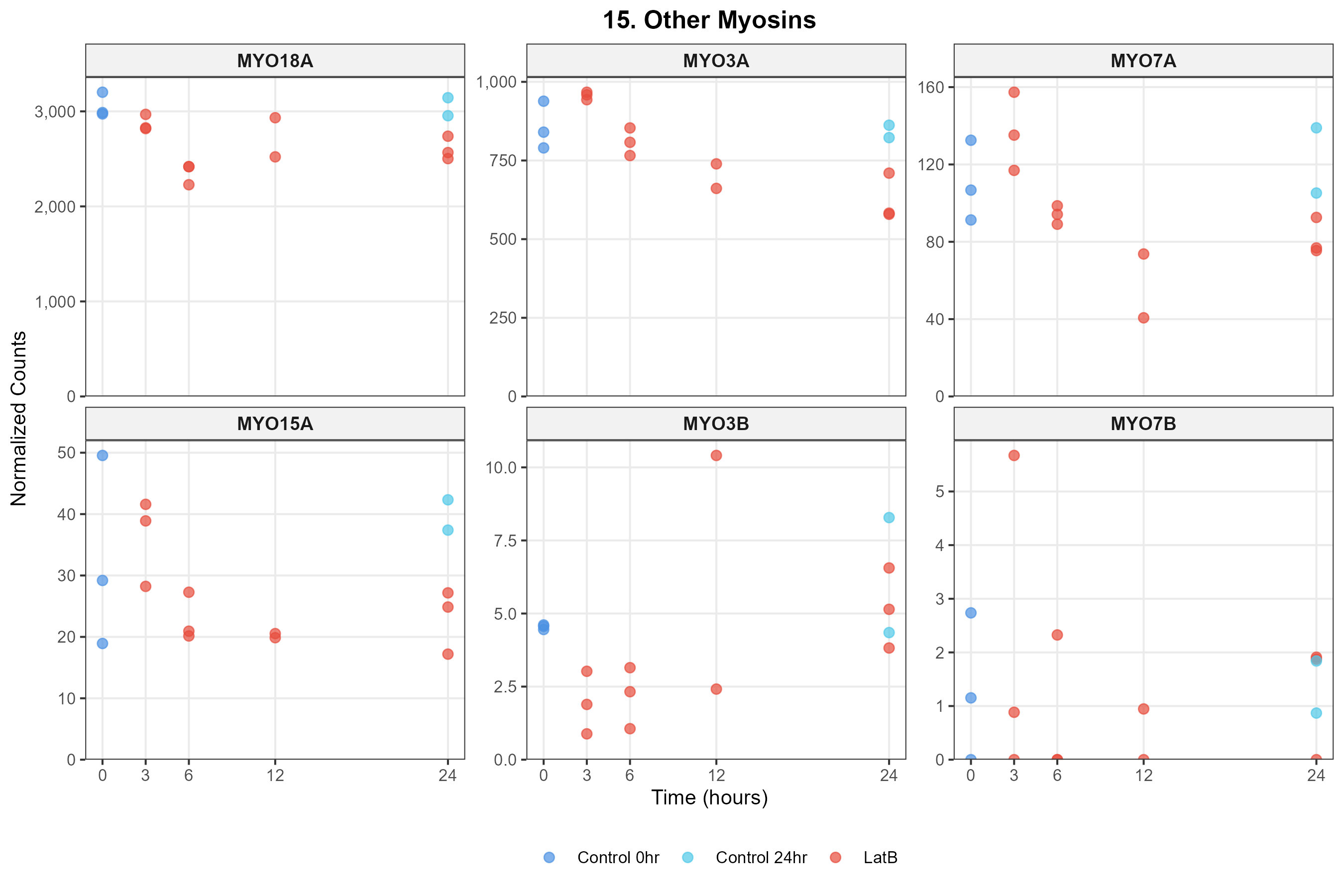

### _Profilin.png

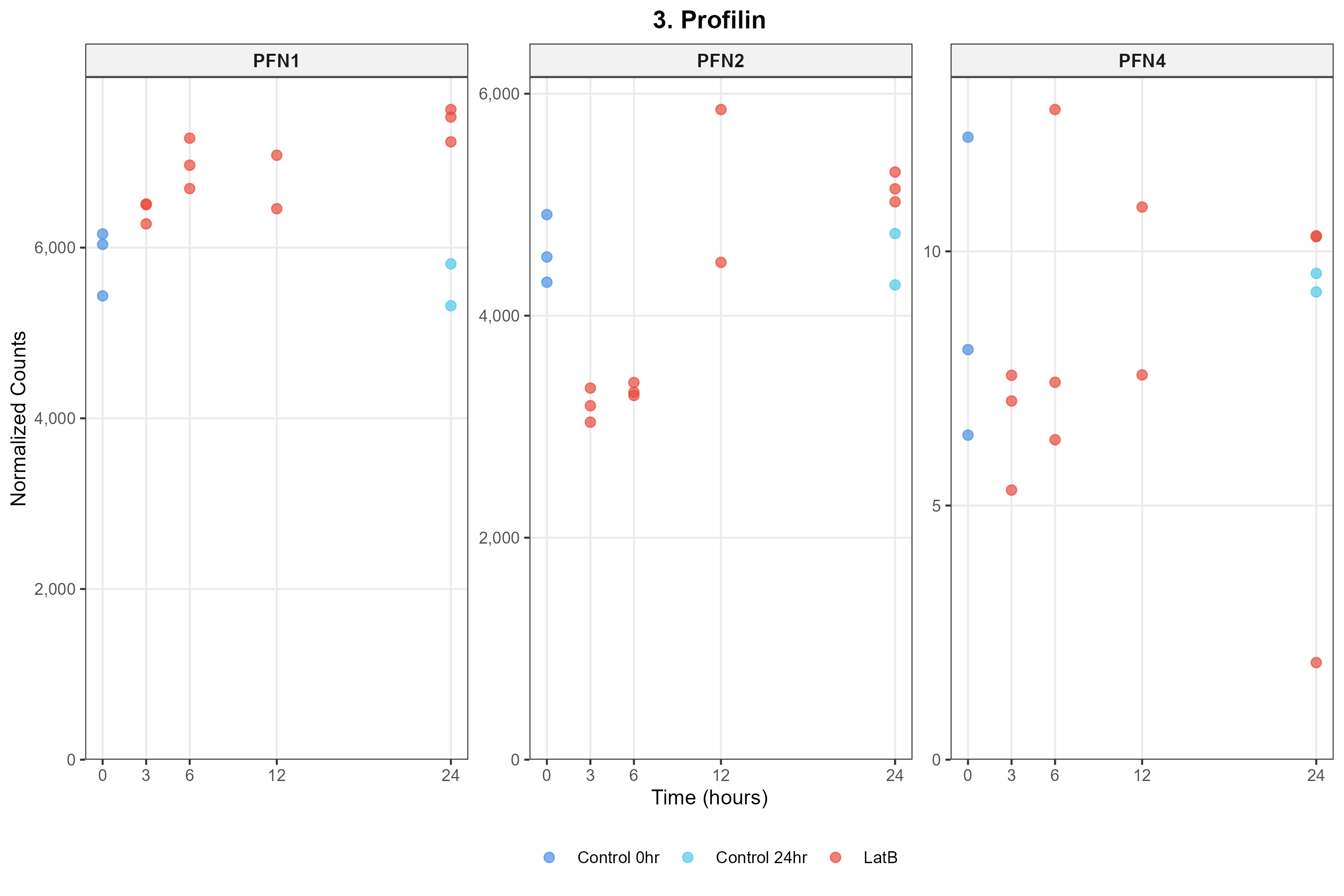

### _Rho_GAPs.png

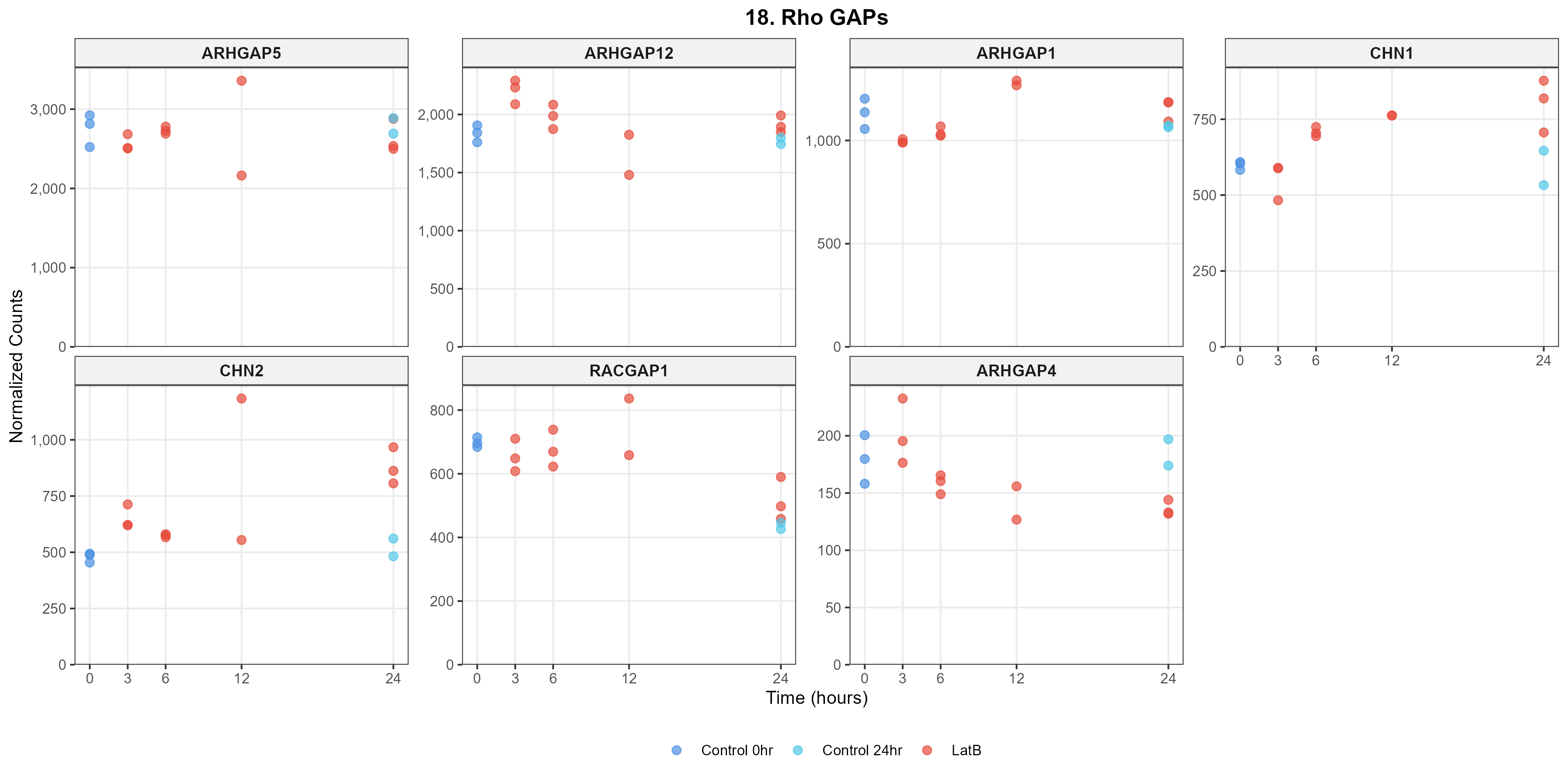

### _Rho_GEFs.png

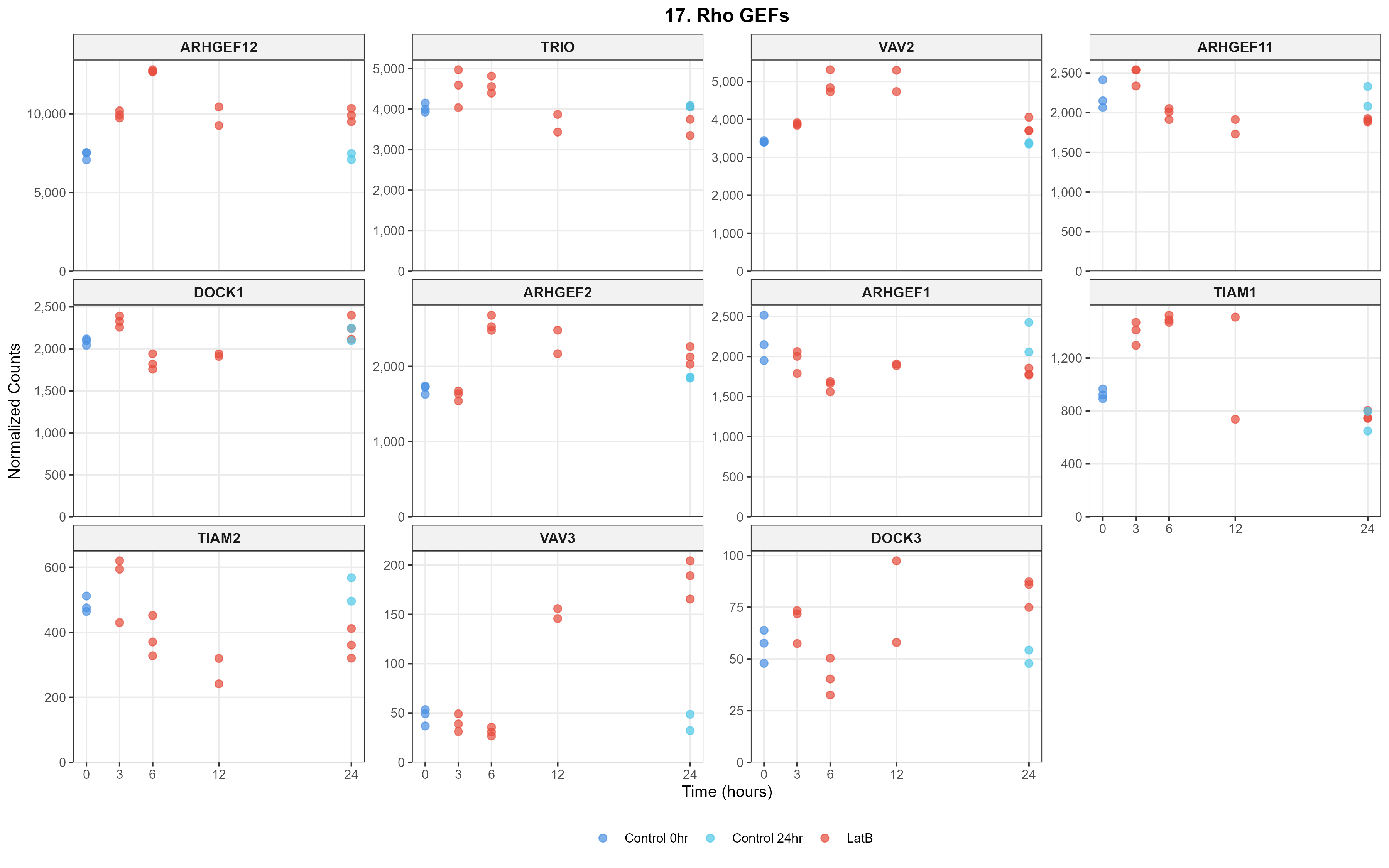

### _Rho_GTPases.png

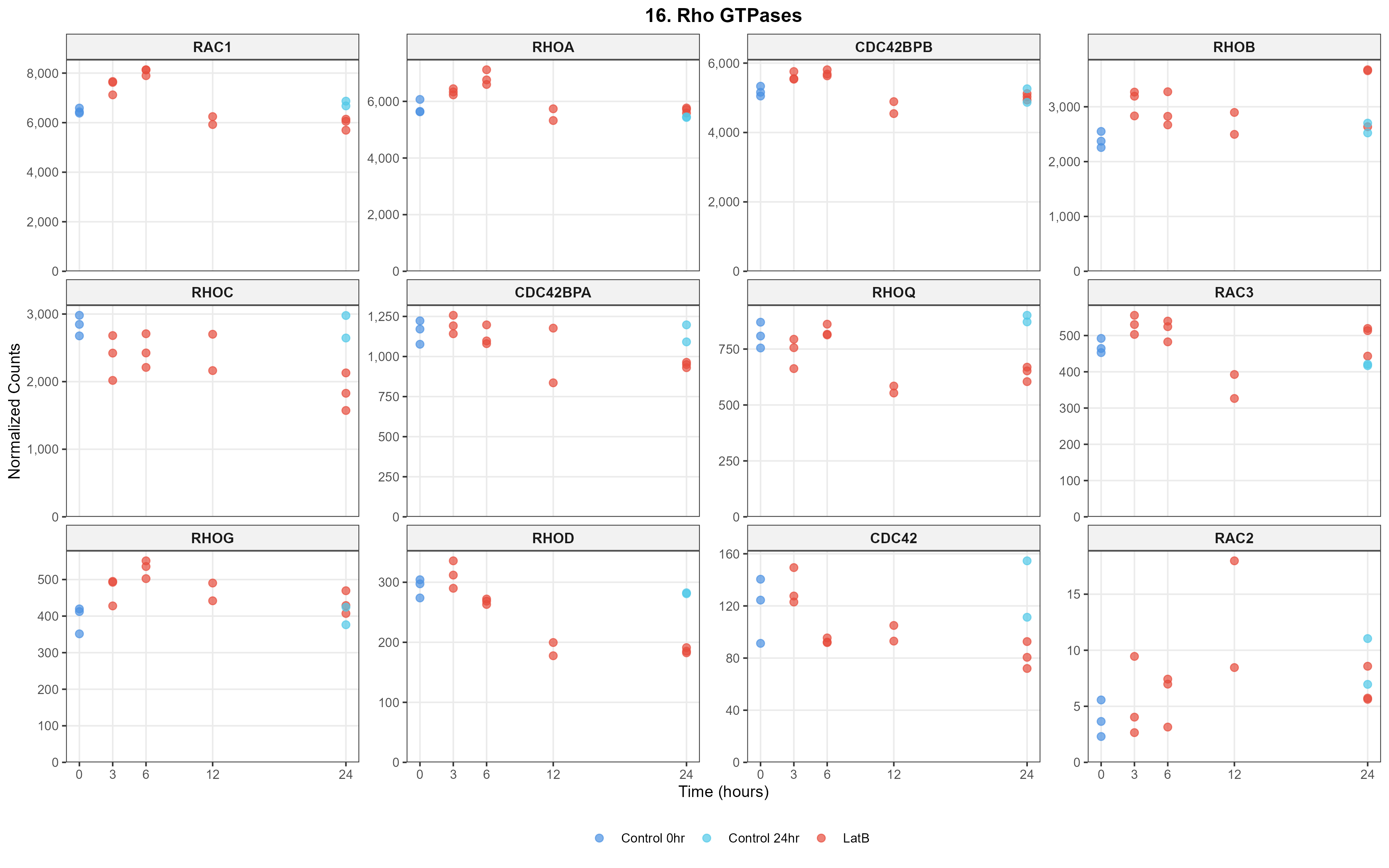

### _Thymosin.png

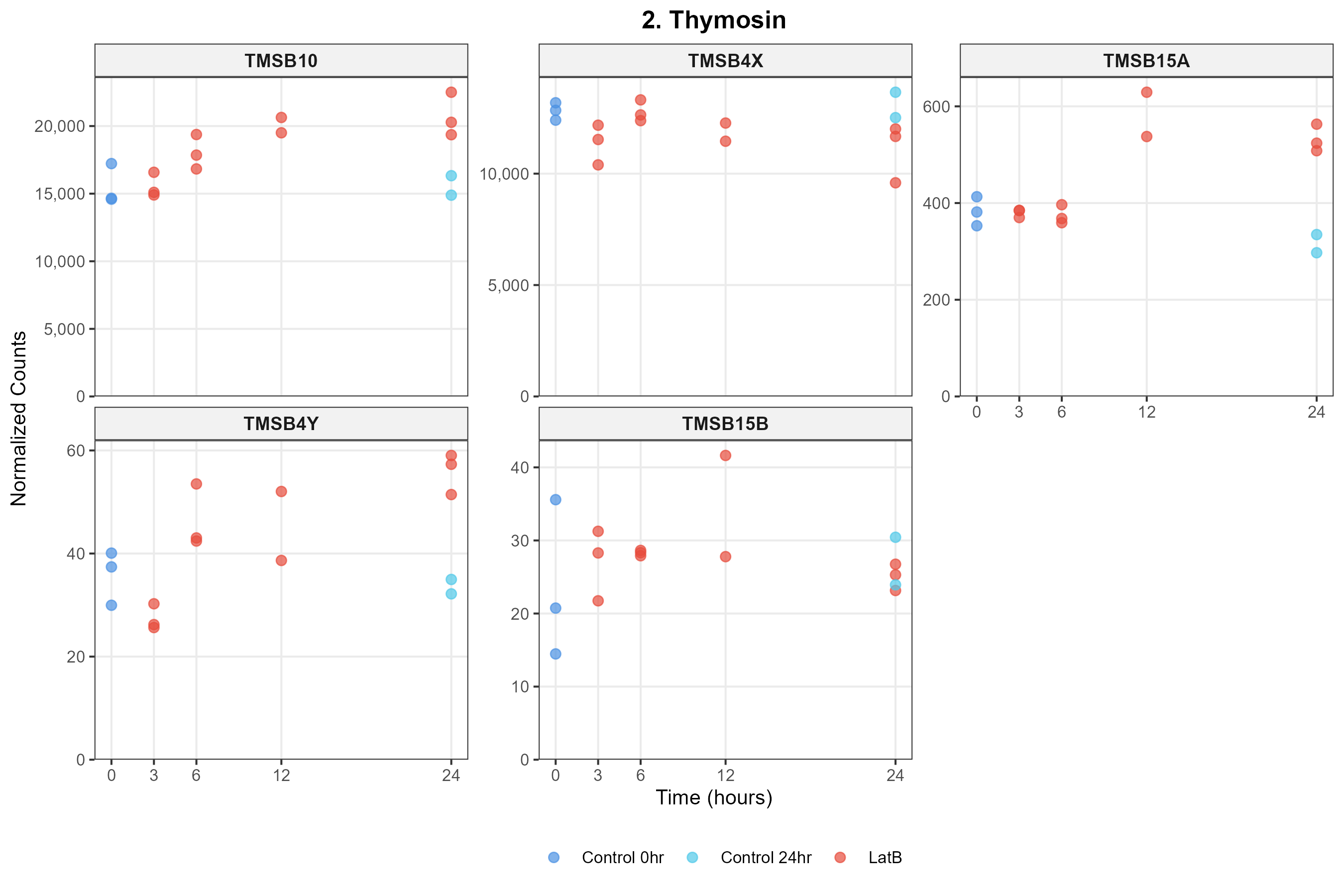

### _Tropomyosin_Tropomodulin.png

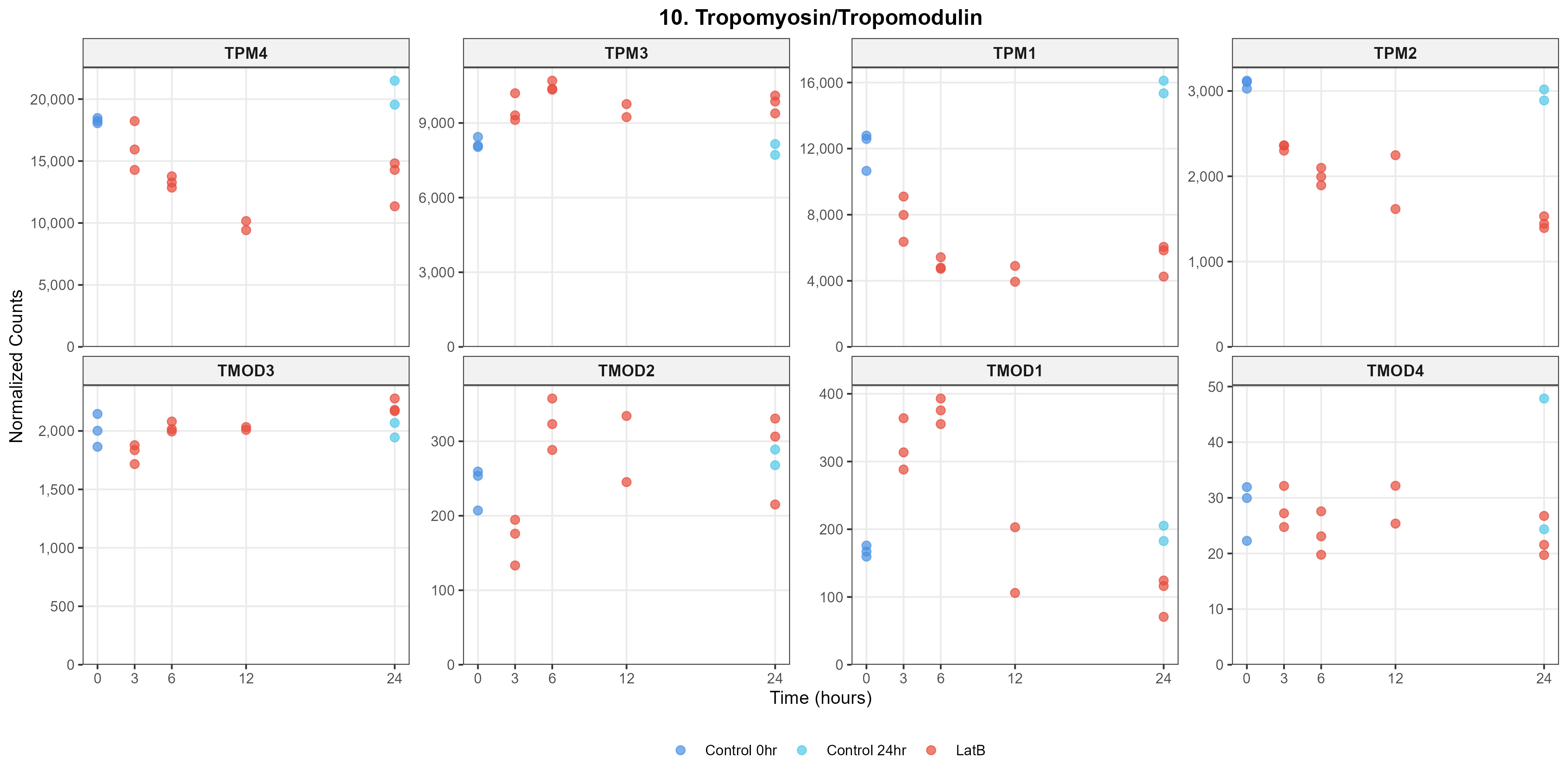

### IF_genes.png

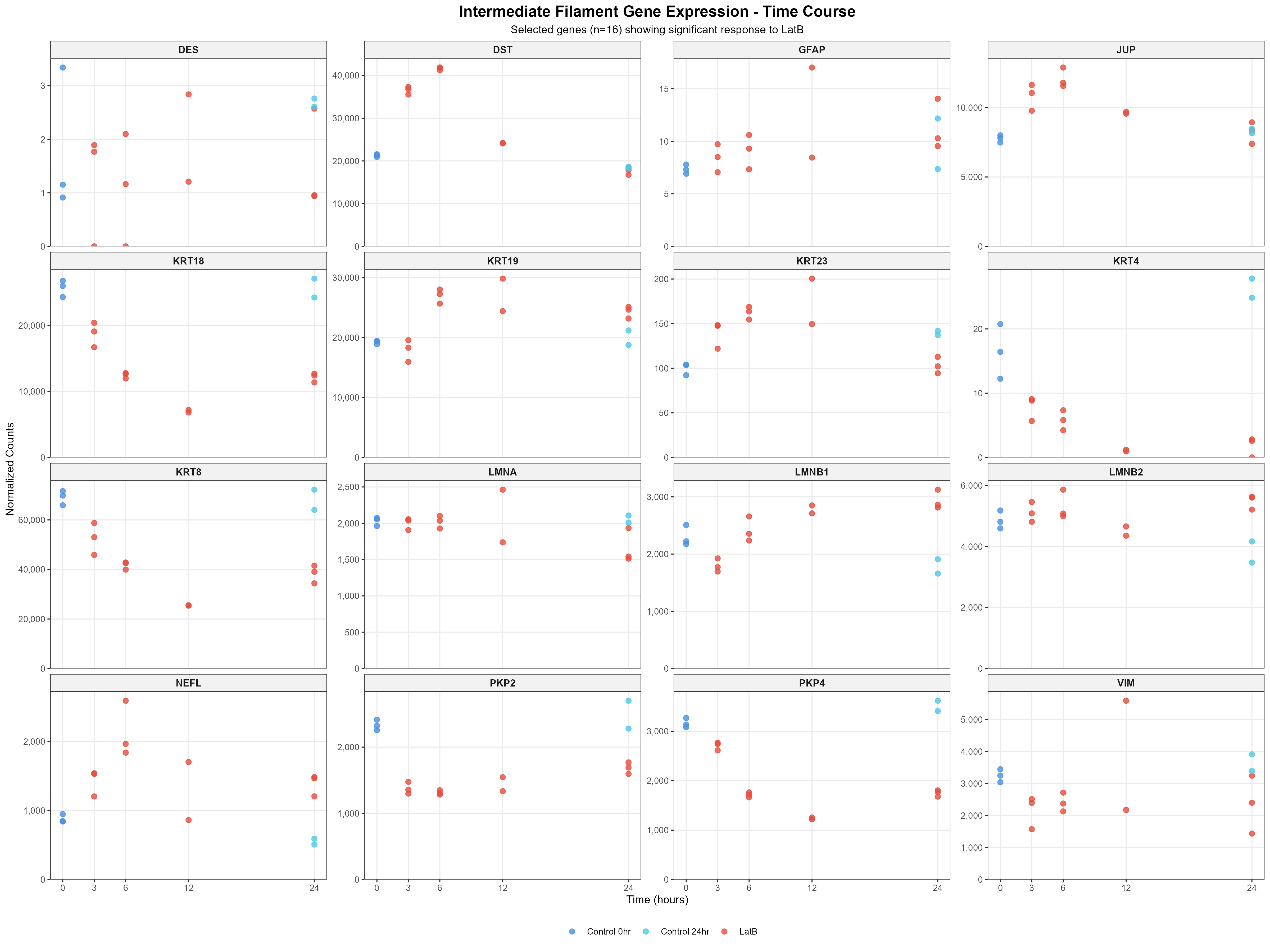

### Microtubule_genes.png

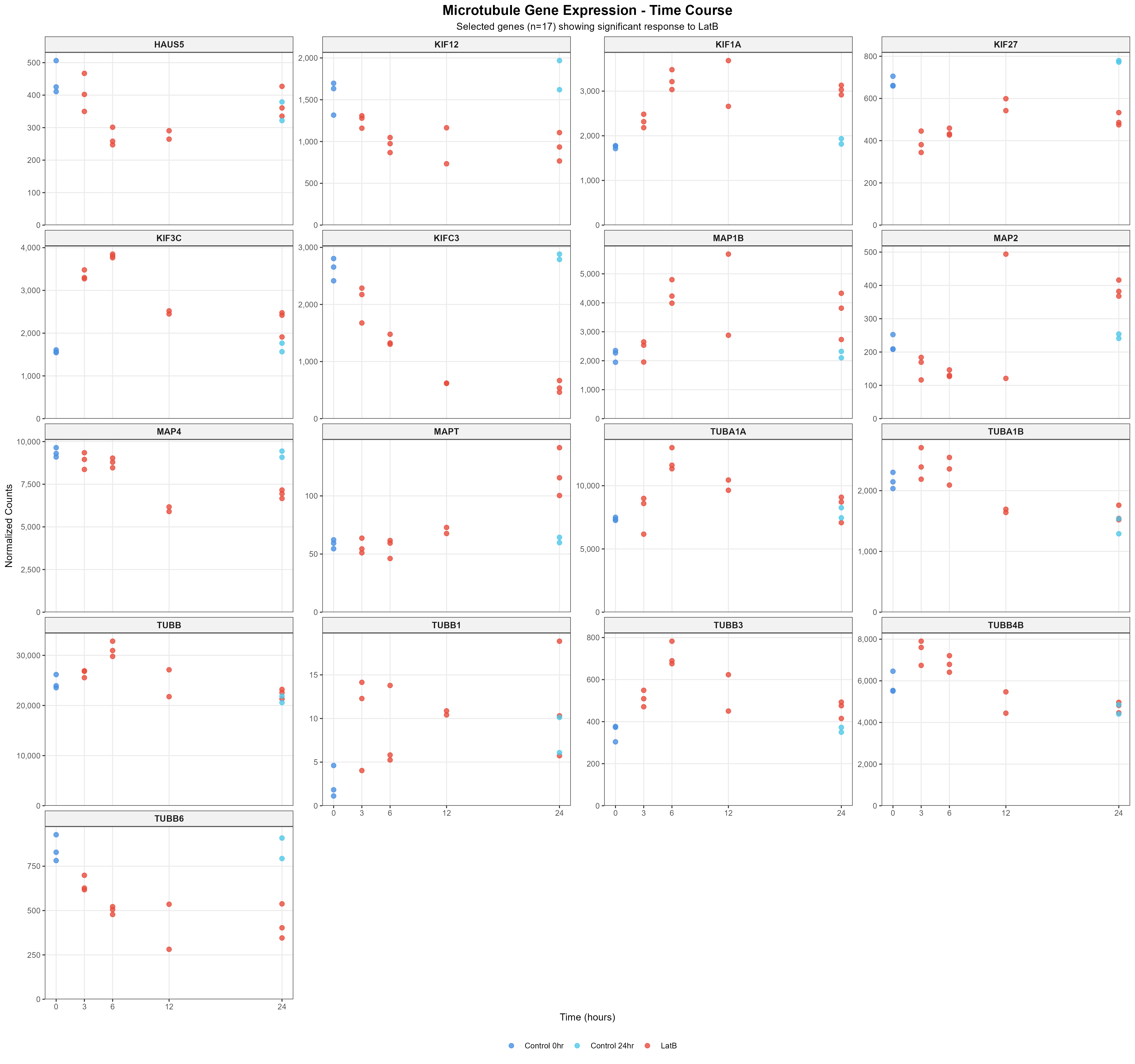
