## Supplementary material for "Analysis of gene expression changes after Latrunculin B-induced actin depolymerization": Scripts: README.md

# LatB RNA-seq — GSE275775 Analysis Code

Computational analysis pipeline characterizing the transcriptional response of human
pancreatic progenitor cells to acute actin depolymerization by latrunculin B (LatB),
in the Vavylonis Lab (Lehigh University). The analysis covers genome-wide differential
expression, cytoskeletal gene family responses (actin, microtubule, and intermediate
filament systems), transcription factor activity inference, and comparison against
prior latrunculin studies.

This repository accompanies the manuscript in preparation. All computational analysis
was performed by Georgios Kargas.

## Repository structure

- `scripts/R/` — Data processing scripts: ChEA3 transcription factor enrichment
  queries and formatting, differential expression extraction (actin genes and
  genome-wide), decoupleR transcription factor activity inference, the DESeq2
  pipeline (per-timepoint and pooled comparisons), gene functional categorization,
  MRTF/SRF target gene extraction, and time-course summary chart data.

Manuscript figure-generation scripts will be added to this structure separately.

## Reference dataset

GEO accession [GSE275775](https://www.ncbi.nlm.nih.gov/geo/query/acc.cgi?acc=GSE275775).
16 samples across latrunculin B treatment (3, 6, 12, 24 hr) and matched controls
(0, 24 hr).

## Environment

- R, with DESeq2, decoupleR, tidyverse, ggplot2
- ChEA3 (transcription factor enrichment) queried via its public API

Each script contains its own header describing its specific analysis and inputs.

## AI assistance

Parts of the code, analysis scripts and documentation in this repository were
developed with the assistance of Claude (Anthropic). The authors ran the analyses
and verified the results.

## License

All code in this repository is released under the MIT License (see `LICENSE`).

## Contact

Georgios Kargas, Vavylonis Lab, Lehigh University.
