## Supplementary material for "Analysis of gene expression changes after Latrunculin B-induced actin depolymerization": Scripts: chea3_input_top300.R

```
library(dplyr)

# Run with the working directory set to Latb_rnaseq/scripts. Only SYMBOL,
# log2FC_LatB_vs_Control and padj_LatB_vs_Control are needed, so this reads
# directly from the shipped pooled-comparison file (Supplementary Data File 5)
# rather than requiring deseq2_pooled.R to be run first.
SDF <- "../Supplementary_Data_Files"
PIPE <- "Pipeline_Outputs"
dir.create(PIPE, showWarnings = FALSE, recursive = TRUE)

de <- read.csv(file.path(SDF, "S5_all_genes_DE_Pooled_Fig1A.csv"),
               stringsAsFactors = FALSE)

# Strict filter: only keep clean standard gene symbols
# Must start with a letter, contain only letters/numbers/hyphens/dots
is_valid_symbol <- function(x) {
  !is.na(x) &
    x != "" &
    nchar(x) > 0 &
    grepl("^[A-Za-z][A-Za-z0-9\\-\\.]*$", x) &
    !grepl("^LINC|^MIR|^SNOR|^RNA|^MT-|^RPS|^RPL|-AS[0-9]|-DT$|-IT[0-9]|HG$|P[0-9]$", x)
}

up_genes <- de %>%
  filter(!is.na(padj_LatB_vs_Control),
         padj_LatB_vs_Control < 0.05,
         log2FC_LatB_vs_Control > 0.5,
         is_valid_symbol(SYMBOL)) %>%
  arrange(desc(log2FC_LatB_vs_Control)) %>%
  pull(SYMBOL) %>%
  unique() %>%
  head(300)

down_genes <- de %>%
  filter(!is.na(padj_LatB_vs_Control),
         padj_LatB_vs_Control < 0.05,
         log2FC_LatB_vs_Control < -0.5,
         is_valid_symbol(SYMBOL)) %>%
  arrange(log2FC_LatB_vs_Control) %>%
  pull(SYMBOL) %>%
  unique() %>%
  head(300)

cat("Upregulated:", length(up_genes), "\n")
cat("Downregulated:", length(down_genes), "\n")

# Write with no blank lines — explicit newline control
writeLines(up_genes[nchar(up_genes) > 0],
           file.path(PIPE, "ChEA3_top300_up_clean.txt"))
writeLines(down_genes[nchar(down_genes) > 0],
           file.path(PIPE, "ChEA3_top300_down_clean.txt"))

# Verify no blank lines in output
up_check   <- readLines(file.path(PIPE, "ChEA3_top300_up_clean.txt"))
down_check <- readLines(file.path(PIPE, "ChEA3_top300_down_clean.txt"))

cat("\nBlank lines in up file:",   sum(up_check == ""), "\n")
cat("Blank lines in down file:", sum(down_check == ""), "\n")

cat("\nPreview upregulated (first 10):\n")
cat(head(up_genes, 10), sep = "\n")
cat("\n\nPreview downregulated (first 10):\n")
cat(head(down_genes, 10), sep = "\n")
```
