## Supplementary material for "Analysis of gene expression changes after Latrunculin B-induced actin depolymerization": Scripts: chea3_query.R

```
# Load required packages
if (!require("httr")) install.packages("httr", repos = "https://cloud.r-project.org")
if (!require("jsonlite")) install.packages("jsonlite", repos = "https://cloud.r-project.org")
if (!require("dplyr")) install.packages("dplyr", repos = "https://cloud.r-project.org")
if (!require("ggplot2")) install.packages("ggplot2", repos = "https://cloud.r-project.org")
if (!require("tidyr")) install.packages("tidyr", repos = "https://cloud.r-project.org")
if (!require("gridExtra")) install.packages("gridExtra", repos = "https://cloud.r-project.org")

library(httr)
library(jsonlite)
library(dplyr)
library(ggplot2)
library(tidyr)
library(gridExtra)

# ============================================================================
# STEP 1: Load Data
# ============================================================================

cat("=== STEP 1: Loading RNA-seq data ===\n\n")

# Run with the working directory set to Latb_rnaseq/scripts. Only SYMBOL and
# the per-timepoint log2FC/padj columns are needed (no "category" column), so
# this reads directly from the shipped normalized-counts+DE table
# (Supplementary Data File 8) rather than requiring gene_categorization.R.
SDF <- "../Supplementary_Data_Files"
PIPE <- "Pipeline_Outputs"
dir.create(PIPE, showWarnings = FALSE, recursive = TRUE)

deg_data <- read.csv(file.path(SDF, "S8_All_Genes_DE_Normalized_Counts.csv"),
                     stringsAsFactors = FALSE)

cat("Total genes in dataset:", nrow(deg_data), "\n\n")

# ============================================================================
# STEP 2: Define significance criteria
# ============================================================================

padj_threshold <- 0.05
log2fc_threshold <- 0.5  # |log2FC| > 0.5

# Check significance at ANY timepoint (3hr, 6hr, 12hr, 24hr vs Control_0hr)
deg_data <- deg_data %>%
  filter(SYMBOL != "") %>%  # Remove empty gene symbols
  mutate(
    # Significant at any timepoint
    sig_3hr = !is.na(padj_LatB_3hr_vs_Control_0hr) & padj_LatB_3hr_vs_Control_0hr < padj_threshold,
    sig_6hr = !is.na(padj_LatB_6hr_vs_Control_0hr) & padj_LatB_6hr_vs_Control_0hr < padj_threshold,
    sig_12hr = !is.na(padj_LatB_12hr_vs_Control_0hr) & padj_LatB_12hr_vs_Control_0hr < padj_threshold,
    sig_24hr = !is.na(padj_LatB_24hr_vs_Control_0hr) & padj_LatB_24hr_vs_Control_0hr < padj_threshold,
    
    # Direction at peak response (use 6hr as representative - peak for many genes)
    direction = case_when(
      log2FC_LatB_6hr_vs_Control_0hr < -log2fc_threshold ~ "down",
      log2FC_LatB_6hr_vs_Control_0hr > log2fc_threshold ~ "up",
      TRUE ~ "unchanged"
    ),
    
    # Minimum log2FC across timepoints (most negative)
    min_log2fc = pmin(log2FC_LatB_3hr_vs_Control_0hr, 
                      log2FC_LatB_6hr_vs_Control_0hr,
                      log2FC_LatB_12hr_vs_Control_0hr,
                      log2FC_LatB_24hr_vs_Control_0hr, na.rm = TRUE),
    
    # Maximum log2FC across timepoints (most positive)
    max_log2fc = pmax(log2FC_LatB_3hr_vs_Control_0hr, 
                      log2FC_LatB_6hr_vs_Control_0hr,
                      log2FC_LatB_12hr_vs_Control_0hr,
                      log2FC_LatB_24hr_vs_Control_0hr, na.rm = TRUE),
    
    # Ever significant
    ever_sig = sig_3hr | sig_6hr | sig_12hr | sig_24hr,
    
    # Significantly downregulated (sig + negative FC)
    sig_down = ever_sig & min_log2fc < -log2fc_threshold,
    
    # Significantly upregulated (sig + positive FC)
    sig_up = ever_sig & max_log2fc > log2fc_threshold
  )

# ============================================================================
# STEP 3: Extract gene lists
# ============================================================================

cat("=== STEP 2: Extracting significant DEGs ===\n\n")

# Downregulated genes
down_genes <- deg_data %>%
  filter(sig_down) %>%
  pull(SYMBOL) %>%
  unique()

# Upregulated genes
up_genes <- deg_data %>%
  filter(sig_up) %>%
  pull(SYMBOL) %>%
  unique()

# All significant genes
all_sig_genes <- deg_data %>%
  filter(ever_sig & (abs(min_log2fc) > log2fc_threshold | abs(max_log2fc) > log2fc_threshold)) %>%
  pull(SYMBOL) %>%
  unique()

cat("Significantly DOWNREGULATED genes:", length(down_genes), "\n")
cat("Significantly UPREGULATED genes:", length(up_genes), "\n")
cat("Total significant DEGs:", length(all_sig_genes), "\n\n")

# Save gene lists
writeLines(down_genes, file.path(PIPE, "ChEA3_Downregulated_Genes.txt"))
writeLines(up_genes, file.path(PIPE, "ChEA3_Upregulated_Genes.txt"))
writeLines(all_sig_genes, file.path(PIPE, "ChEA3_All_Significant_Genes.txt"))

cat("Saved gene lists to text files\n\n")

# ============================================================================
# STEP 4: ChEA3 Query Function
# ============================================================================

query_chea3 <- function(gene_list, description = "query") {
  
  url <- "https://maayanlab.cloud/chea3/api/enrich/"
  
  payload <- list(
    query_name = description,
    gene_set = gene_list
  )
  
  cat("Querying ChEA3 with", length(gene_list), "genes...\n")
  
  response <- tryCatch({
    POST(
      url,
      body = toJSON(payload, auto_unbox = TRUE),
      content_type("application/json"),
      timeout(120)
    )
  }, error = function(e) {
    cat("Error:", e$message, "\n")
    return(NULL)
  })
  
  if (is.null(response)) return(NULL)
  
  if (status_code(response) != 200) {
    cat("API error. Status:", status_code(response), "\n")
    return(NULL)
  }
  
  result <- fromJSON(content(response, "text", encoding = "UTF-8"))
  cat("Success!\n\n")
  return(result)
}

# ============================================================================
# STEP 5: Run ChEA3 on both gene lists
# ============================================================================

cat("=== STEP 3: Running ChEA3 Enrichment ===\n\n")

# Query downregulated genes
cat("--- Analyzing DOWNREGULATED genes ---\n")
chea3_down <- query_chea3(down_genes, "LatB_Downregulated")

# Query upregulated genes
cat("--- Analyzing UPREGULATED genes ---\n")
chea3_up <- query_chea3(up_genes, "LatB_Upregulated")

# ============================================================================
# STEP 6: Process Results
# ============================================================================

cat("=== STEP 4: Processing Results ===\n\n")

# Function to extract integrated results
extract_results <- function(chea3_result, direction) {
  if (is.null(chea3_result)) return(NULL)
  
  if ("Integrated--meanRank" %in% names(chea3_result)) {
    df <- as.data.frame(chea3_result[["Integrated--meanRank"]])
    df$Direction <- direction
    df$Rank <- as.numeric(df$Rank)
    df$Score <- as.numeric(df$Score)
    return(df %>% arrange(Rank))
  }
  return(NULL)
}

results_down <- extract_results(chea3_down, "Downregulated")
results_up <- extract_results(chea3_up, "Upregulated")

# ============================================================================
# STEP 7: Find MRTF/SRF and YAP pathway TFs
# ============================================================================

cat("=== STEP 5: Finding MRTF/SRF and YAP/TEAD TFs ===\n\n")

# Key TFs to look for
mrtf_srf_tfs <- c("SRF", "MRTFA", "MRTFB", "MKL1", "MKL2", "MYOCD")
yap_tead_tfs <- c("YAP1", "WWTR1", "TAZ", "TEAD1", "TEAD2", "TEAD3", "TEAD4")
all_target_tfs <- c(mrtf_srf_tfs, yap_tead_tfs)

# Search in downregulated results
cat("--- In DOWNREGULATED genes ---\n")
if (!is.null(results_down)) {
  for (tf in all_target_tfs) {
    match <- results_down %>% filter(grepl(paste0("^", tf, "$"), TF, ignore.case = TRUE))
    if (nrow(match) > 0) {
      cat(sprintf("  %s: Rank #%d (Score: %.2f)\n", match$TF[1], match$Rank[1], match$Score[1]))
    }
  }
}

cat("\n--- In UPREGULATED genes ---\n")
if (!is.null(results_up)) {
  for (tf in all_target_tfs) {
    match <- results_up %>% filter(grepl(paste0("^", tf, "$"), TF, ignore.case = TRUE))
    if (nrow(match) > 0) {
      cat(sprintf("  %s: Rank #%d (Score: %.2f)\n", match$TF[1], match$Rank[1], match$Score[1]))
    }
  }
}

# ============================================================================
# STEP 8: Create Summary Tables
# ============================================================================

cat("\n=== STEP 6: Top 30 TFs ===\n\n")

cat("--- Top 30 TFs for DOWNREGULATED genes ---\n")
if (!is.null(results_down)) {
  print(head(results_down[, c("Rank", "TF", "Score")], 30))
}

cat("\n--- Top 30 TFs for UPREGULATED genes ---\n")
if (!is.null(results_up)) {
  print(head(results_up[, c("Rank", "TF", "Score")], 30))
}

# ============================================================================
# STEP 9: Save Full Results
# ============================================================================

cat("\n=== STEP 7: Saving Results ===\n\n")

if (!is.null(results_down)) {
  write.csv(results_down, file.path(PIPE, "ChEA3_Results_Downregulated_Full.csv"), row.names = FALSE)
  cat("Saved: ChEA3_Results_Downregulated_Full.csv\n")
}

if (!is.null(results_up)) {
  write.csv(results_up, file.path(PIPE, "ChEA3_Results_Upregulated_Full.csv"), row.names = FALSE)
  cat("Saved: ChEA3_Results_Upregulated_Full.csv\n")
}

# ============================================================================
# STEP 10: Create Visualizations
# ============================================================================

cat("\n=== STEP 8: Creating Visualizations ===\n\n")

# Define TF categories for coloring
categorize_tf <- function(tf) {
  case_when(
    grepl("^SRF$|^MRTF|^MKL|^MYOCD", tf, ignore.case = TRUE) ~ "MRTF/SRF",
    grepl("^YAP|^WWTR1|^TAZ$|^TEAD", tf, ignore.case = TRUE) ~ "YAP/TEAD",
    grepl("^FOX", tf, ignore.case = TRUE) ~ "FOX family",
    grepl("^SMAD", tf, ignore.case = TRUE) ~ "SMAD/TGFβ",
    grepl("^STAT", tf, ignore.case = TRUE) ~ "STAT/JAK",
    grepl("^NF.?B|^REL", tf, ignore.case = TRUE) ~ "NF-κB",
    grepl("^AP.?1|^FOS|^JUN", tf, ignore.case = TRUE) ~ "AP-1",
    TRUE ~ "Other"
  )
}

# Color palette
category_colors <- c(
  "MRTF/SRF" = "#E41A1C",
  "YAP/TEAD" = "#FF7F00",
  "FOX family" = "#4DAF4A",
  "SMAD/TGFβ" = "#984EA3",
  "STAT/JAK" = "#377EB8",
  "NF-κB" = "#A65628",
  "AP-1" = "#F781BF",
  "Other" = "#999999"
)

# Plot function
create_tf_plot <- function(results_df, title, n_top = 30) {
  if (is.null(results_df)) return(NULL)
  
  plot_data <- head(results_df, n_top) %>%
    mutate(
      Category = categorize_tf(TF),
      Category = factor(Category, levels = names(category_colors))
    )
  
  ggplot(plot_data, aes(x = reorder(TF, -Rank), y = Score, fill = Category)) +
    geom_bar(stat = "identity", width = 0.8) +
    scale_fill_manual(values = category_colors, drop = FALSE) +
    coord_flip() +
    labs(
      title = title,
      subtitle = paste("Top", n_top, "transcription factors by ChEA3 enrichment"),
      x = NULL,
      y = "Mean Rank Score (lower = stronger enrichment)"
    ) +
    theme_minimal(base_size = 11) +
    theme(
      plot.title = element_text(size = 14, face = "bold"),
      plot.subtitle = element_text(size = 10, color = "gray40"),
      axis.text.y = element_text(size = 9),
      legend.position = "right",
      legend.title = element_text(size = 10, face = "bold")
    )
}

# Create plots
p_down <- create_tf_plot(results_down, "TF Enrichment: DOWNREGULATED Genes")
p_up <- create_tf_plot(results_up, "TF Enrichment: UPREGULATED Genes")

# Save individual plots
if (!is.null(p_down)) {
  ggsave(file.path(PIPE, "ChEA3_Downregulated_Top30_TFs.png"), p_down, width = 12, height = 10, dpi = 300)
  ggsave(file.path(PIPE, "ChEA3_Downregulated_Top30_TFs.pdf"), p_down, width = 12, height = 10)
  cat("Saved: ChEA3_Downregulated_Top30_TFs.png/pdf\n")
}

if (!is.null(p_up)) {
  ggsave(file.path(PIPE, "ChEA3_Upregulated_Top30_TFs.png"), p_up, width = 12, height = 10, dpi = 300)
  ggsave(file.path(PIPE, "ChEA3_Upregulated_Top30_TFs.pdf"), p_up, width = 12, height = 10)
  cat("Saved: ChEA3_Upregulated_Top30_TFs.png/pdf\n")
}

# Combined figure
if (!is.null(p_down) && !is.null(p_up)) {
  combined <- grid.arrange(p_down, p_up, ncol = 2)
  ggsave(file.path(PIPE, "ChEA3_Combined_TF_Analysis.png"), combined, width = 20, height = 10, dpi = 300)
  ggsave(file.path(PIPE, "ChEA3_Combined_TF_Analysis.pdf"), combined, width = 20, height = 10)
  cat("Saved: ChEA3_Combined_TF_Analysis.png/pdf\n")
}

# ============================================================================
# STEP 11: Summary Report
# ============================================================================

cat("\n")
cat("============================================================\n")
cat("SUMMARY: UNBIASED TF PATHWAY ANALYSIS\n")
cat("============================================================\n")
cat("\n")
cat("Input:\n")
cat("  - Downregulated genes:", length(down_genes), "\n")
cat("  - Upregulated genes:", length(up_genes), "\n")
cat("\n")
cat("KEY FINDINGS - MRTF/SRF Pathway:\n")
if (!is.null(results_down)) {
  srf_rank <- results_down %>% filter(TF == "SRF") %>% pull(Rank)
  if (length(srf_rank) > 0) {
    cat("  - SRF rank in DOWNREGULATED:", srf_rank[1], "\n")
  }
}
cat("\n")
cat("KEY FINDINGS - YAP/TEAD Pathway:\n")
if (!is.null(results_down)) {
  for (tf in c("TEAD1", "TEAD2", "TEAD3", "TEAD4")) {
    tf_rank <- results_down %>% filter(TF == tf) %>% pull(Rank)
    if (length(tf_rank) > 0) {
      cat(sprintf("  - %s rank in DOWNREGULATED: %d\n", tf, tf_rank[1]))
    }
  }
}
cat("\n")
cat("Output files:\n")
cat("  - ChEA3_Results_Downregulated_Full.csv\n")
cat("  - ChEA3_Results_Upregulated_Full.csv\n")
cat("  - ChEA3_Downregulated_Top30_TFs.png/pdf\n")
cat("  - ChEA3_Upregulated_Top30_TFs.png/pdf\n")
cat("  - ChEA3_Combined_TF_Analysis.png/pdf\n")
cat("============================================================\n")
```
