## Supplementary material for "Analysis of gene expression changes after Latrunculin B-induced actin depolymerization": Scripts: chea3_temporal_analysis.R

```
# Load required packages
if (!require("httr")) install.packages("httr", repos = "https://cloud.r-project.org")
if (!require("jsonlite")) install.packages("jsonlite", repos = "https://cloud.r-project.org")
if (!require("dplyr")) install.packages("dplyr", repos = "https://cloud.r-project.org")
if (!require("ggplot2")) install.packages("ggplot2", repos = "https://cloud.r-project.org")
if (!require("tidyr")) install.packages("tidyr", repos = "https://cloud.r-project.org")
if (!require("gridExtra")) install.packages("gridExtra", repos = "https://cloud.r-project.org")
if (!require("pheatmap")) install.packages("pheatmap", repos = "https://cloud.r-project.org")
if (!require("tibble")) install.packages("tibble", repos = "https://cloud.r-project.org")

library(httr)
library(jsonlite)
library(dplyr)
library(ggplot2)
library(tidyr)
library(gridExtra)
library(pheatmap)
library(tibble)

# Recovered from an archived copy of the original analysis. This is the
# upstream step chea3_wide_format.R needs: per-timepoint and per-response-
# pattern ChEA3 enrichment, plus the temporal pattern classification itself.
# Run with the working directory set to Latb_rnaseq/scripts.
SDF <- "../Supplementary_Data_Files"
PIPE <- "Pipeline_Outputs"
dir.create(PIPE, showWarnings = FALSE, recursive = TRUE)

cat("=============================================================\n")
cat("IMPROVED ChEA3 TEMPORAL ANALYSIS - CORRECTED VERSION\n")
cat("=============================================================\n\n")

#-------------------------------------------------------------------------------
# GLOBAL DEFINITIONS (moved to top scope)
#-------------------------------------------------------------------------------

# TF categorization function
categorize_tf <- function(tf) {
  case_when(
    grepl("^SRF$|^MRTF|^MKL|^MYOCD", tf, ignore.case = TRUE) ~ "MRTF/SRF",
    grepl("^YAP|^WWTR1|^TAZ$|^TEAD", tf, ignore.case = TRUE) ~ "YAP/TEAD",
    grepl("^FOX", tf, ignore.case = TRUE) ~ "FOX family",
    grepl("^SMAD", tf, ignore.case = TRUE) ~ "SMAD/TGFb",
    grepl("^STAT", tf, ignore.case = TRUE) ~ "STAT/JAK",
    grepl("^NF.?B|^REL", tf, ignore.case = TRUE) ~ "NF-kB",
    grepl("^AP.?1|^FOS|^JUN", tf, ignore.case = TRUE) ~ "AP-1",
    TRUE ~ "Other"
  )
}

# Color palette for categories
category_colors <- c(
  "MRTF/SRF" = "#E41A1C",
  "YAP/TEAD" = "#FF7F00",
  "FOX family" = "#4DAF4A",
  "SMAD/TGFb" = "#984EA3",
  "STAT/JAK" = "#377EB8",
  "NF-kB" = "#A65628",
  "AP-1" = "#F781BF",
  "Other" = "#999999"
)

# Key TFs to track
target_tfs <- c("SRF", "MRTFA", "MKL1", "MKL2", "MYOCD",
                "TEAD1", "TEAD2", "TEAD3", "TEAD4", "YAP1", "WWTR1")

# Parameters
padj_threshold <- 0.05
log2fc_threshold <- 0.5

#-------------------------------------------------------------------------------
# STEP 1: Load Data
#-------------------------------------------------------------------------------
cat("=== STEP 1: Loading RNA-seq data ===\n\n")

# Only SYMBOL, ensembl_id and the per-comparison log2FC/padj columns are
# needed, so this reads directly from the shipped normalized-counts+DE table
# (Supplementary Data File 8) rather than a hand-versioned intermediate file.
deg_data <- read.csv(file.path(SDF, "S8_All_Genes_DE_Normalized_Counts.csv"),
                     stringsAsFactors = FALSE)

cat("Total genes in dataset:", nrow(deg_data), "\n")

#-------------------------------------------------------------------------------
# STEP 2: Validate Required Columns
#-------------------------------------------------------------------------------
cat("\n=== STEP 2: Validating required columns ===\n\n")

# Check for 24hr control columns
required_24hr_cols <- c("padj_LatB_24hr_vs_Control_24hr", "log2FC_LatB_24hr_vs_Control_24hr")
required_early_cols <- c(
  "padj_LatB_3hr_vs_Control_0hr", "log2FC_LatB_3hr_vs_Control_0hr",
  "padj_LatB_6hr_vs_Control_0hr", "log2FC_LatB_6hr_vs_Control_0hr",
  "padj_LatB_12hr_vs_Control_0hr", "log2FC_LatB_12hr_vs_Control_0hr",
  "padj_LatB_24hr_vs_Control_0hr", "log2FC_LatB_24hr_vs_Control_0hr"
)

all_required <- c(required_24hr_cols, required_early_cols)
missing_cols <- setdiff(all_required, colnames(deg_data))

if (length(missing_cols) > 0) {
  cat("WARNING: Missing columns:", paste(missing_cols, collapse = ", "), "\n")
  if (any(required_24hr_cols %in% missing_cols)) {
    stop("ERROR: Missing critical 24hr control comparison columns. Cannot proceed.")
  }
} else {
  cat("All required columns present.\n")
}

# Check for proper 24hr control availability
has_24hr_control <- all(required_24hr_cols %in% colnames(deg_data))
cat("Using proper 24hr control (LatB_24hr vs Control_24hr):", has_24hr_control, "\n")

#-------------------------------------------------------------------------------
# STEP 3: Calculate Significance at Each Timepoint
#-------------------------------------------------------------------------------
cat("\n=== STEP 3: Calculating temporal significance ===\n\n")

deg_data <- deg_data %>%
  filter(SYMBOL != "" & !is.na(SYMBOL)) %>%
  mutate(
    # EARLY TIMEPOINTS: Compare to 0hr control
    sig_3hr = !is.na(padj_LatB_3hr_vs_Control_0hr) &
      padj_LatB_3hr_vs_Control_0hr < padj_threshold &
      abs(log2FC_LatB_3hr_vs_Control_0hr) > log2fc_threshold,

    sig_6hr = !is.na(padj_LatB_6hr_vs_Control_0hr) &
      padj_LatB_6hr_vs_Control_0hr < padj_threshold &
      abs(log2FC_LatB_6hr_vs_Control_0hr) > log2fc_threshold,

    sig_12hr = !is.na(padj_LatB_12hr_vs_Control_0hr) &
      padj_LatB_12hr_vs_Control_0hr < padj_threshold &
      abs(log2FC_LatB_12hr_vs_Control_0hr) > log2fc_threshold,

    # 24HR: Use PROPER control (LatB_24hr vs Control_24hr)
    sig_24hr = !is.na(padj_LatB_24hr_vs_Control_24hr) &
      padj_LatB_24hr_vs_Control_24hr < padj_threshold &
      abs(log2FC_LatB_24hr_vs_Control_24hr) > log2fc_threshold,

    # Direction at each timepoint
    dir_3hr = case_when(
      sig_3hr & log2FC_LatB_3hr_vs_Control_0hr < 0 ~ "down",
      sig_3hr & log2FC_LatB_3hr_vs_Control_0hr > 0 ~ "up",
      TRUE ~ "ns"
    ),
    dir_6hr = case_when(
      sig_6hr & log2FC_LatB_6hr_vs_Control_0hr < 0 ~ "down",
      sig_6hr & log2FC_LatB_6hr_vs_Control_0hr > 0 ~ "up",
      TRUE ~ "ns"
    ),
    dir_12hr = case_when(
      sig_12hr & log2FC_LatB_12hr_vs_Control_0hr < 0 ~ "down",
      sig_12hr & log2FC_LatB_12hr_vs_Control_0hr > 0 ~ "up",
      TRUE ~ "ns"
    ),
    dir_24hr = case_when(
      sig_24hr & log2FC_LatB_24hr_vs_Control_24hr < 0 ~ "down",
      sig_24hr & log2FC_LatB_24hr_vs_Control_24hr > 0 ~ "up",
      TRUE ~ "ns"
    )
  )

cat("Genes with valid symbols:", nrow(deg_data), "\n")

#-------------------------------------------------------------------------------
# STEP 4: Peak Response Detection
#-------------------------------------------------------------------------------
cat("\n=== STEP 4: Detecting peak response for each gene ===\n\n")

deg_data <- deg_data %>%
  mutate(
    # Find peak absolute fold change across timepoints
    abs_fc_3hr = abs(log2FC_LatB_3hr_vs_Control_0hr),
    abs_fc_6hr = abs(log2FC_LatB_6hr_vs_Control_0hr),
    abs_fc_12hr = abs(log2FC_LatB_12hr_vs_Control_0hr),
    abs_fc_24hr = abs(log2FC_LatB_24hr_vs_Control_24hr),

    peak_abs_fc = pmax(abs_fc_3hr, abs_fc_6hr, abs_fc_12hr, abs_fc_24hr, na.rm = TRUE),

    # Determine which timepoint has peak
    peak_time = case_when(
      abs_fc_3hr == peak_abs_fc ~ "3hr",
      abs_fc_6hr == peak_abs_fc ~ "6hr",
      abs_fc_12hr == peak_abs_fc ~ "12hr",
      abs_fc_24hr == peak_abs_fc ~ "24hr",
      TRUE ~ NA_character_
    ),

    # Fold change at peak time
    peak_log2fc = case_when(
      peak_time == "3hr" ~ log2FC_LatB_3hr_vs_Control_0hr,
      peak_time == "6hr" ~ log2FC_LatB_6hr_vs_Control_0hr,
      peak_time == "12hr" ~ log2FC_LatB_12hr_vs_Control_0hr,
      peak_time == "24hr" ~ log2FC_LatB_24hr_vs_Control_24hr
    ),

    # padj at peak time
    peak_padj = case_when(
      peak_time == "3hr" ~ padj_LatB_3hr_vs_Control_0hr,
      peak_time == "6hr" ~ padj_LatB_6hr_vs_Control_0hr,
      peak_time == "12hr" ~ padj_LatB_12hr_vs_Control_0hr,
      peak_time == "24hr" ~ padj_LatB_24hr_vs_Control_24hr
    ),

    # Significant at peak
    sig_at_peak = !is.na(peak_padj) & peak_padj < padj_threshold &
      abs(peak_log2fc) > log2fc_threshold,

    # Direction at peak
    peak_direction = case_when(
      sig_at_peak & peak_log2fc < 0 ~ "down",
      sig_at_peak & peak_log2fc > 0 ~ "up",
      TRUE ~ "unchanged"
    )
  )

#-------------------------------------------------------------------------------
# STEP 5: Temporal Pattern Classification (WITH MIXED DETECTION)
#-------------------------------------------------------------------------------
cat("=== STEP 5: Classifying temporal patterns ===\n\n")

# Helper function to detect mixed direction responses
is_mixed_direction <- function(dir_3hr, dir_6hr, dir_12hr, dir_24hr) {
  dirs <- c(dir_3hr, dir_6hr, dir_12hr, dir_24hr)
  dirs <- dirs[dirs != "ns"]  # Remove non-significant
  if (length(dirs) < 2) return(FALSE)
  length(unique(dirs)) > 1
}

deg_data <- deg_data %>%
  rowwise() %>%
  mutate(
    is_mixed = is_mixed_direction(dir_3hr, dir_6hr, dir_12hr, dir_24hr)
  ) %>%
  ungroup() %>%
  mutate(
    # Temporal patterns
    temporal_pattern = case_when(
      # First check for mixed responses (up at some timepoints, down at others)
      is_mixed ~ "mixed_response",

      # Early transient DOWN: significant early, not at 24hr
      (dir_3hr == "down" | dir_6hr == "down") & dir_24hr != "down" ~ "early_transient_down",

      # Early transient UP: significant early, not at 24hr
      (dir_3hr == "up" | dir_6hr == "up") & dir_24hr != "up" ~ "early_transient_up",

      # Sustained DOWN: down at 24hr AND at least one earlier timepoint
      dir_24hr == "down" & (dir_3hr == "down" | dir_6hr == "down" | dir_12hr == "down") ~ "sustained_down",

      # Sustained UP: up at 24hr AND at least one earlier timepoint
      dir_24hr == "up" & (dir_3hr == "up" | dir_6hr == "up" | dir_12hr == "up") ~ "sustained_up",

      # Late DOWN: only significant at 24hr
      dir_24hr == "down" & dir_3hr == "ns" & dir_6hr == "ns" & dir_12hr == "ns" ~ "late_only_down",

      # Late UP: only significant at 24hr
      dir_24hr == "up" & dir_3hr == "ns" & dir_6hr == "ns" & dir_12hr == "ns" ~ "late_only_up",

      # 12hr only patterns
      dir_12hr == "down" & dir_3hr == "ns" & dir_6hr == "ns" & dir_24hr == "ns" ~ "mid_only_down",
      dir_12hr == "up" & dir_3hr == "ns" & dir_6hr == "ns" & dir_24hr == "ns" ~ "mid_only_up",

      # Ever significant but doesn't fit above
      sig_3hr | sig_6hr | sig_12hr | sig_24hr ~ "other_pattern",

      TRUE ~ "not_significant"
    )
  )

# Count patterns
pattern_counts <- deg_data %>%
  count(temporal_pattern) %>%
  arrange(desc(n))

cat("Temporal pattern distribution:\n")
print(pattern_counts)

#-------------------------------------------------------------------------------
# STEP 6: Create Gene Lists (using named lists - FIXED)
#-------------------------------------------------------------------------------
cat("\n=== STEP 6: Creating gene lists ===\n\n")

# Store all gene lists in a named list (FIXED: avoids get() bug)
gene_lists <- list()

# A. TIMEPOINT-SPECIFIC LISTS
gene_lists$down_3hr <- deg_data %>% filter(dir_3hr == "down") %>% pull(SYMBOL) %>% unique()
gene_lists$up_3hr <- deg_data %>% filter(dir_3hr == "up") %>% pull(SYMBOL) %>% unique()

gene_lists$down_6hr <- deg_data %>% filter(dir_6hr == "down") %>% pull(SYMBOL) %>% unique()
gene_lists$up_6hr <- deg_data %>% filter(dir_6hr == "up") %>% pull(SYMBOL) %>% unique()

gene_lists$down_12hr <- deg_data %>% filter(dir_12hr == "down") %>% pull(SYMBOL) %>% unique()
gene_lists$up_12hr <- deg_data %>% filter(dir_12hr == "up") %>% pull(SYMBOL) %>% unique()

gene_lists$down_24hr <- deg_data %>% filter(dir_24hr == "down") %>% pull(SYMBOL) %>% unique()
gene_lists$up_24hr <- deg_data %>% filter(dir_24hr == "up") %>% pull(SYMBOL) %>% unique()

# B. PEAK RESPONSE LISTS
gene_lists$peak_down <- deg_data %>% filter(peak_direction == "down") %>% pull(SYMBOL) %>% unique()
gene_lists$peak_up <- deg_data %>% filter(peak_direction == "up") %>% pull(SYMBOL) %>% unique()

# C. TEMPORAL PATTERN LISTS
gene_lists$early_transient_down <- deg_data %>% filter(temporal_pattern == "early_transient_down") %>% pull(SYMBOL) %>% unique()
gene_lists$sustained_down <- deg_data %>% filter(temporal_pattern == "sustained_down") %>% pull(SYMBOL) %>% unique()
gene_lists$late_only_down <- deg_data %>% filter(temporal_pattern == "late_only_down") %>% pull(SYMBOL) %>% unique()

gene_lists$early_transient_up <- deg_data %>% filter(temporal_pattern == "early_transient_up") %>% pull(SYMBOL) %>% unique()
gene_lists$sustained_up <- deg_data %>% filter(temporal_pattern == "sustained_up") %>% pull(SYMBOL) %>% unique()
gene_lists$late_only_up <- deg_data %>% filter(temporal_pattern == "late_only_up") %>% pull(SYMBOL) %>% unique()

gene_lists$mixed_response <- deg_data %>% filter(temporal_pattern == "mixed_response") %>% pull(SYMBOL) %>% unique()

cat("Gene list sizes:\n")
cat("\n--- TIMEPOINT-SPECIFIC ---\n")
cat("3hr DOWN:", length(gene_lists$down_3hr), "| 3hr UP:", length(gene_lists$up_3hr), "\n")
cat("6hr DOWN:", length(gene_lists$down_6hr), "| 6hr UP:", length(gene_lists$up_6hr), "\n")
cat("12hr DOWN:", length(gene_lists$down_12hr), "| 12hr UP:", length(gene_lists$up_12hr), "\n")
cat("24hr DOWN:", length(gene_lists$down_24hr), "| 24hr UP:", length(gene_lists$up_24hr), "(vs 24hr Control)\n")

cat("\n--- PEAK RESPONSE ---\n")
cat("Peak DOWN:", length(gene_lists$peak_down), "| Peak UP:", length(gene_lists$peak_up), "\n")

cat("\n--- TEMPORAL PATTERNS ---\n")
cat("Early transient DOWN:", length(gene_lists$early_transient_down), "| UP:", length(gene_lists$early_transient_up), "\n")
cat("Sustained DOWN:", length(gene_lists$sustained_down), "| UP:", length(gene_lists$sustained_up), "\n")
cat("Late only DOWN:", length(gene_lists$late_only_down), "| UP:", length(gene_lists$late_only_up), "\n")
cat("Mixed response:", length(gene_lists$mixed_response), "\n")

#-------------------------------------------------------------------------------
# STEP 7: ChEA3 Query Functions
#-------------------------------------------------------------------------------

query_chea3 <- function(gene_list, description = "query") {
  if (length(gene_list) < 10) {
    cat("  Skipping", description, "- too few genes (", length(gene_list), ")\n")
    return(NULL)
  }

  url <- "https://maayanlab.cloud/chea3/api/enrich/"

  payload <- list(
    query_name = description,
    gene_set = gene_list
  )

  cat("  Querying ChEA3:", description, "(", length(gene_list), "genes)...")

  response <- tryCatch({
    POST(
      url,
      body = toJSON(payload, auto_unbox = TRUE),
      content_type("application/json"),
      timeout(120)
    )
  }, error = function(e) {
    cat(" Error:", e$message, "\n")
    return(NULL)
  })

  if (is.null(response)) return(NULL)

  if (status_code(response) != 200) {
    cat(" API error. Status:", status_code(response), "\n")
    return(NULL)
  }

  result <- fromJSON(content(response, "text", encoding = "UTF-8"))
  cat(" Success!\n")
  return(result)
}

extract_results <- function(chea3_result, description) {
  if (is.null(chea3_result)) return(NULL)

  if ("Integrated--meanRank" %in% names(chea3_result)) {
    df <- as.data.frame(chea3_result[["Integrated--meanRank"]])
    df$Analysis <- description
    df$Rank <- as.numeric(df$Rank)
    df$Score <- as.numeric(df$Score)
    df <- df %>% arrange(Rank)
    return(df)
  }
  return(NULL)
}

#-------------------------------------------------------------------------------
# STEP 8: Run ChEA3 Analyses (WITH PROGRESS TRACKING)
#-------------------------------------------------------------------------------
cat("\n=== STEP 7: Running ChEA3 enrichment analyses ===\n\n")

all_results <- list()

# Define all analyses to run
analyses_to_run <- list(
  # Timepoint-specific DOWN
  list(key = "down_3hr", desc = "Down_3hr"),
  list(key = "down_6hr", desc = "Down_6hr"),
  list(key = "down_12hr", desc = "Down_12hr"),
  list(key = "down_24hr", desc = "Down_24hr"),
  # Timepoint-specific UP
  list(key = "up_3hr", desc = "Up_3hr"),
  list(key = "up_6hr", desc = "Up_6hr"),
  list(key = "up_12hr", desc = "Up_12hr"),
  list(key = "up_24hr", desc = "Up_24hr"),
  # Peak response
  list(key = "peak_down", desc = "Peak_Down"),
  list(key = "peak_up", desc = "Peak_Up"),
  # Temporal patterns
  list(key = "early_transient_down", desc = "Early_Transient_Down"),
  list(key = "sustained_down", desc = "Sustained_Down"),
  list(key = "late_only_down", desc = "Late_Only_Down"),
  list(key = "early_transient_up", desc = "Early_Transient_Up"),
  list(key = "sustained_up", desc = "Sustained_Up"),
  list(key = "late_only_up", desc = "Late_Only_Up"),
  list(key = "mixed_response", desc = "Mixed_Response")
)

# Progress tracking
total_analyses <- length(analyses_to_run)
cat("Running", total_analyses, "ChEA3 analyses...\n\n")

for (i in seq_along(analyses_to_run)) {
  analysis <- analyses_to_run[[i]]

  cat(sprintf("[%d/%d] ", i, total_analyses))

  genes <- gene_lists[[analysis$key]]

  if (!is.null(genes) && length(genes) >= 10) {
    result <- query_chea3(genes, analysis$desc)
    all_results[[analysis$key]] <- extract_results(result, analysis$desc)
  } else {
    cat("  Skipping", analysis$desc, "- insufficient genes\n")
  }

  Sys.sleep(1.5)  # Be nice to the API
}

cat("\nCompleted", sum(!sapply(all_results, is.null)), "analyses successfully.\n")

#-------------------------------------------------------------------------------
# STEP 9: Extract Key TF Rankings
#-------------------------------------------------------------------------------
cat("\n=== STEP 8: Extracting MRTF/SRF and YAP/TEAD rankings ===\n\n")

get_tf_ranks <- function(results_df, tf_list) {
  if (is.null(results_df)) return(NULL)
  results_df %>%
    filter(TF %in% tf_list) %>%
    select(TF, Rank, Score) %>%
    arrange(Rank)
}

# Create summary table
tf_summary <- data.frame()

for (analysis_name in names(all_results)) {
  if (!is.null(all_results[[analysis_name]])) {
    tf_ranks <- get_tf_ranks(all_results[[analysis_name]], target_tfs)
    if (!is.null(tf_ranks) && nrow(tf_ranks) > 0) {
      tf_ranks$Analysis <- analysis_name
      tf_summary <- bind_rows(tf_summary, tf_ranks)
    }
  }
}

# Pivot for easy comparison
if (nrow(tf_summary) > 0) {
  tf_comparison <- tf_summary %>%
    select(TF, Rank, Analysis) %>%
    pivot_wider(names_from = Analysis, values_from = Rank)

  cat("TF Rankings across analyses:\n")
  print(tf_comparison)
}

#-------------------------------------------------------------------------------
# STEP 10: Create Heatmap Visualization
#-------------------------------------------------------------------------------
cat("\n=== STEP 9: Creating visualizations ===\n\n")

if (nrow(tf_summary) > 0) {
  # Prepare heatmap data for all TFs that appear in multiple analyses
  heatmap_data <- tf_summary %>%
    select(TF, Rank, Analysis) %>%
    pivot_wider(names_from = Analysis, values_from = Rank, values_fill = NA) %>%
    column_to_rownames("TF")

  # Keep TFs that appear in at least 2 analyses
  heatmap_data <- heatmap_data[rowSums(!is.na(heatmap_data)) >= 2, , drop = FALSE]

  if (nrow(heatmap_data) > 0 && ncol(heatmap_data) > 1) {
    # Create heatmap
    png(file.path(PIPE, "ChEA3_TF_Temporal_Heatmap.png"), width = 12, height = 8, units = "in", res = 300)
    pheatmap(as.matrix(heatmap_data),
             cluster_rows = TRUE,
             cluster_cols = TRUE,
             color = colorRampPalette(c("darkblue", "blue", "white", "orange", "red"))(100),
             main = "TF Ranks Across Temporal Analyses\n(Lower rank = stronger enrichment)",
             fontsize = 10,
             display_numbers = TRUE,
             number_format = "%.0f",
             na_col = "grey90")
    dev.off()
    cat("Saved: ChEA3_TF_Temporal_Heatmap.png\n")
  }
}

#-------------------------------------------------------------------------------
# STEP 11: Create Bar Plot Comparisons
#-------------------------------------------------------------------------------

# Plot function
create_tf_plot <- function(results_df, title, n_top = 25) {
  if (is.null(results_df) || nrow(results_df) == 0) return(NULL)

  plot_data <- head(results_df, n_top) %>%
    mutate(
      Category = categorize_tf(TF),
      Category = factor(Category, levels = names(category_colors))
    )

  ggplot(plot_data, aes(x = reorder(TF, -Rank), y = Score, fill = Category)) +
    geom_bar(stat = "identity", width = 0.8) +
    scale_fill_manual(values = category_colors, drop = FALSE) +
    coord_flip() +
    labs(
      title = title,
      x = NULL,
      y = "Mean Rank Score (lower = stronger enrichment)"
    ) +
    theme_minimal(base_size = 10) +
    theme(
      plot.title = element_text(size = 11, face = "bold"),
      legend.position = "right",
      legend.title = element_text(size = 9)
    )
}

# Compare 3hr vs 24hr for downregulated
if (!is.null(all_results$down_3hr) && !is.null(all_results$down_24hr)) {
  p_3hr <- create_tf_plot(all_results$down_3hr, "Downregulated @ 3hr\n(Immediate response)")
  p_24hr <- create_tf_plot(all_results$down_24hr, "Downregulated @ 24hr\n(vs 24hr Control)")

  combined <- grid.arrange(p_3hr, p_24hr, ncol = 2)

  ggsave(file.path(PIPE, "ChEA3_Temporal_Comparison_3hr_vs_24hr_Down.png"), combined, width = 18, height = 10, dpi = 300)
  ggsave(file.path(PIPE, "ChEA3_Temporal_Comparison_3hr_vs_24hr_Down.pdf"), combined, width = 18, height = 10)
  cat("Saved: ChEA3_Temporal_Comparison_3hr_vs_24hr_Down.png/pdf\n")
}

# Compare temporal patterns for downregulated
pattern_analyses <- c("early_transient_down", "sustained_down", "late_only_down")
pattern_plots <- list()

for (pattern in pattern_analyses) {
  if (!is.null(all_results[[pattern]]) && nrow(all_results[[pattern]]) > 0) {
    title <- gsub("_", " ", tools::toTitleCase(pattern))
    pattern_plots[[pattern]] <- create_tf_plot(all_results[[pattern]], title, n_top = 20) +
      theme(legend.position = "none")
  }
}

if (length(pattern_plots) >= 2) {
  combined_patterns <- do.call(grid.arrange, c(pattern_plots, ncol = length(pattern_plots)))
  ggsave(file.path(PIPE, "ChEA3_Temporal_Patterns_Down.png"), combined_patterns,
         width = 6 * length(pattern_plots), height = 8, dpi = 300)
  ggsave(file.path(PIPE, "ChEA3_Temporal_Patterns_Down.pdf"), combined_patterns,
         width = 6 * length(pattern_plots), height = 8)
  cat("Saved: ChEA3_Temporal_Patterns_Down.png/pdf\n")
}

#-------------------------------------------------------------------------------
# STEP 12: Save Results
#-------------------------------------------------------------------------------
cat("\n=== STEP 10: Saving results ===\n\n")

# Save all ChEA3 results
for (name in names(all_results)) {
  if (!is.null(all_results[[name]])) {
    filename <- file.path(PIPE, paste0("ChEA3_", name, "_Full.csv"))
    write.csv(all_results[[name]], filename, row.names = FALSE)
    cat("Saved:", filename, "\n")
  }
}

# Save TF summary
if (nrow(tf_summary) > 0) {
  write.csv(tf_summary, file.path(PIPE, "ChEA3_TF_Summary_All_Analyses.csv"), row.names = FALSE)
  cat("Saved: ChEA3_TF_Summary_All_Analyses.csv\n")

  # Also save the wide format comparison
  write.csv(tf_comparison, file.path(PIPE, "ChEA3_TF_Comparison_Wide.csv"), row.names = FALSE)
  cat("Saved: ChEA3_TF_Comparison_Wide.csv\n")
}

# Save gene lists
for (name in names(gene_lists)) {
  if (length(gene_lists[[name]]) > 0) {
    writeLines(gene_lists[[name]], file.path(PIPE, paste0("Genes_", name, ".txt")))
  }
}
cat("Saved gene list files\n")

# Save temporal pattern data
deg_data_summary <- deg_data %>%
  select(SYMBOL, ensembl_id, temporal_pattern, peak_time, peak_log2fc, peak_direction,
         dir_3hr, dir_6hr, dir_12hr, dir_24hr, is_mixed)
write.csv(deg_data_summary, file.path(PIPE, "Gene_Temporal_Patterns.csv"), row.names = FALSE)
cat("Saved: Gene_Temporal_Patterns.csv\n")

#-------------------------------------------------------------------------------
# STEP 13: Summary Report
#-------------------------------------------------------------------------------
cat("\n")
cat("============================================================\n")
cat("SUMMARY: IMPROVED TEMPORAL ChEA3 ANALYSIS\n")
cat("============================================================\n\n")

cat("KEY IMPROVEMENTS APPLIED:\n")
cat("1. Used proper 24hr control (LatB_24hr vs Control_24hr)\n")
cat("2. Analyzed each timepoint separately\n")
cat("3. Classified temporal patterns (early, sustained, late, mixed)\n")
cat("4. Detected peak response for each gene\n")
cat("5. Fixed all identified bugs\n\n")

cat("GENE COUNTS BY TEMPORAL PATTERN:\n")
print(pattern_counts)

cat("\n")
if (exists("tf_comparison") && nrow(tf_comparison) > 0) {
  cat("KEY TF RANKINGS COMPARISON:\n\n")

  # Focus on key TFs
  key_tfs <- c("SRF", "TEAD1", "TEAD3")
  key_comparison <- tf_comparison[tf_comparison$TF %in% key_tfs, ]

  if (nrow(key_comparison) > 0) {
    print(key_comparison)
  }

  cat("\nINTERPRETATION:\n")
  cat("- Compare SRF ranks at 3hr vs 24hr to test if MRTF/SRF is early-peaking\n")
  cat("- Compare TEAD ranks at 3hr vs 24hr to test if YAP/TEAD is sustained\n")
  cat("- Look for TFs enriched in 'sustained_down' for core mechanosensing program\n")
}

cat("\n============================================================\n")
cat("Analysis complete!\n")
cat("============================================================\n")
```
