## Supplementary material for "Analysis of gene expression changes after Latrunculin B-induced actin depolymerization": Scripts: chea3_wide_format.R

```
# Run with the working directory set to Latb_rnaseq/scripts. Reads the outputs
# of chea3_temporal_analysis.R (run that first) from the local Pipeline_Outputs
# folder - that script produces the 17 per-timepoint/pattern ChEA3 result
# files, gene lists and Gene_Temporal_Patterns.csv this one needs.
library(dplyr)
library(tidyr)

PIPE <- "Pipeline_Outputs"

cat("=== Loading ALL ChEA3 Results Files ===\n\n")

# -----------------------------------------------------------------------------
# LOAD ALL 17 ChEA3 RESULTS FILES
# -----------------------------------------------------------------------------

# Create a list to store all results
all_results <- list()

# Timepoint-specific - Downregulated
all_results$down_3hr <- read.csv(file.path(PIPE, "ChEA3_down_3hr_Full.csv"), stringsAsFactors = FALSE) %>% 
  select(TF, Rank, Score) %>% mutate(Analysis = "down_3hr")
all_results$down_6hr <- read.csv(file.path(PIPE, "ChEA3_down_6hr_Full.csv"), stringsAsFactors = FALSE) %>% 
  select(TF, Rank, Score) %>% mutate(Analysis = "down_6hr")
all_results$down_12hr <- read.csv(file.path(PIPE, "ChEA3_down_12hr_Full.csv"), stringsAsFactors = FALSE) %>% 
  select(TF, Rank, Score) %>% mutate(Analysis = "down_12hr")
all_results$down_24hr <- read.csv(file.path(PIPE, "ChEA3_down_24hr_Full.csv"), stringsAsFactors = FALSE) %>% 
  select(TF, Rank, Score) %>% mutate(Analysis = "down_24hr")

# Timepoint-specific - Upregulated
all_results$up_3hr <- read.csv(file.path(PIPE, "ChEA3_up_3hr_Full.csv"), stringsAsFactors = FALSE) %>% 
  select(TF, Rank, Score) %>% mutate(Analysis = "up_3hr")
all_results$up_6hr <- read.csv(file.path(PIPE, "ChEA3_up_6hr_Full.csv"), stringsAsFactors = FALSE) %>% 
  select(TF, Rank, Score) %>% mutate(Analysis = "up_6hr")
all_results$up_12hr <- read.csv(file.path(PIPE, "ChEA3_up_12hr_Full.csv"), stringsAsFactors = FALSE) %>% 
  select(TF, Rank, Score) %>% mutate(Analysis = "up_12hr")
all_results$up_24hr <- read.csv(file.path(PIPE, "ChEA3_up_24hr_Full.csv"), stringsAsFactors = FALSE) %>% 
  select(TF, Rank, Score) %>% mutate(Analysis = "up_24hr")

# Peak response
all_results$peak_down <- read.csv(file.path(PIPE, "ChEA3_peak_down_Full.csv"), stringsAsFactors = FALSE) %>% 
  select(TF, Rank, Score) %>% mutate(Analysis = "peak_down")
all_results$peak_up <- read.csv(file.path(PIPE, "ChEA3_peak_up_Full.csv"), stringsAsFactors = FALSE) %>% 
  select(TF, Rank, Score) %>% mutate(Analysis = "peak_up")

# Temporal patterns
all_results$early_transient_down <- read.csv(file.path(PIPE, "ChEA3_early_transient_down_Full.csv"), stringsAsFactors = FALSE) %>% 
  select(TF, Rank, Score) %>% mutate(Analysis = "early_transient_down")
all_results$early_transient_up <- read.csv(file.path(PIPE, "ChEA3_early_transient_up_Full.csv"), stringsAsFactors = FALSE) %>% 
  select(TF, Rank, Score) %>% mutate(Analysis = "early_transient_up")
all_results$sustained_down <- read.csv(file.path(PIPE, "ChEA3_sustained_down_Full.csv"), stringsAsFactors = FALSE) %>% 
  select(TF, Rank, Score) %>% mutate(Analysis = "sustained_down")
all_results$sustained_up <- read.csv(file.path(PIPE, "ChEA3_sustained_up_Full.csv"), stringsAsFactors = FALSE) %>% 
  select(TF, Rank, Score) %>% mutate(Analysis = "sustained_up")
all_results$late_only_down <- read.csv(file.path(PIPE, "ChEA3_late_only_down_Full.csv"), stringsAsFactors = FALSE) %>% 
  select(TF, Rank, Score) %>% mutate(Analysis = "late_only_down")
all_results$late_only_up <- read.csv(file.path(PIPE, "ChEA3_late_only_up_Full.csv"), stringsAsFactors = FALSE) %>% 
  select(TF, Rank, Score) %>% mutate(Analysis = "late_only_up")
all_results$mixed_response <- read.csv(file.path(PIPE, "ChEA3_mixed_response_Full.csv"), stringsAsFactors = FALSE) %>% 
  select(TF, Rank, Score) %>% mutate(Analysis = "mixed_response")

cat("Loaded 17 ChEA3 result files.\n\n")

# -----------------------------------------------------------------------------
# COMBINE ALL RESULTS INTO ONE LONG DATA FRAME
# -----------------------------------------------------------------------------

combined_long <- bind_rows(all_results)

cat("Total rows in combined data:", nrow(combined_long), "\n")
cat("Unique TFs:", length(unique(combined_long$TF)), "\n")
cat("Analyses:", length(unique(combined_long$Analysis)), "\n\n")

# -----------------------------------------------------------------------------
# CREATE WIDE FORMAT TABLE - RANKS
# -----------------------------------------------------------------------------

cat("=== Creating Wide Format Tables ===\n\n")

# Pivot to wide format - RANKS
ranks_wide <- combined_long %>%
  select(TF, Analysis, Rank) %>%
  pivot_wider(
    names_from = Analysis,
    values_from = Rank,
    values_fill = NA
  )

# Reorder columns in logical order
col_order <- c("TF",
               # Downregulated by timepoint
               "down_3hr", "down_6hr", "down_12hr", "down_24hr",
               # Upregulated by timepoint
               "up_3hr", "up_6hr", "up_12hr", "up_24hr",
               # Peak response
               "peak_down", "peak_up",
               # Temporal patterns - down
               "early_transient_down", "sustained_down", "late_only_down",
               # Temporal patterns - up
               "early_transient_up", "sustained_up", "late_only_up",
               # Mixed
               "mixed_response")

ranks_wide <- ranks_wide %>%
  select(all_of(col_order))

# -----------------------------------------------------------------------------
# CREATE WIDE FORMAT TABLE - SCORES
# -----------------------------------------------------------------------------

scores_wide <- combined_long %>%
  select(TF, Analysis, Score) %>%
  pivot_wider(
    names_from = Analysis,
    values_from = Score,
    values_fill = NA
  ) %>%
  select(all_of(col_order))

# -----------------------------------------------------------------------------
# ADD TF CATEGORIZATION
# -----------------------------------------------------------------------------

categorize_tf <- function(tf) {
  case_when(
    grepl("^SRF$|^MRTF|^MKL|^MYOCD", tf, ignore.case = TRUE) ~ "MRTF/SRF",
    grepl("^YAP|^WWTR1|^TAZ$|^TEAD", tf, ignore.case = TRUE) ~ "YAP/TEAD",
    grepl("^FOX", tf, ignore.case = TRUE) ~ "FOX family",
    grepl("^SMAD", tf, ignore.case = TRUE) ~ "SMAD/TGFb",
    grepl("^STAT", tf, ignore.case = TRUE) ~ "STAT/JAK",
    grepl("^NF.?B|^REL", tf, ignore.case = TRUE) ~ "NF-kB",
    grepl("^FOS|^JUN", tf, ignore.case = TRUE) ~ "AP-1",
    grepl("^EGR", tf, ignore.case = TRUE) ~ "Immediate-Early",
    grepl("^ATF", tf, ignore.case = TRUE) ~ "ATF/Stress",
    grepl("^NR4A", tf, ignore.case = TRUE) ~ "NR4A/Orphan",
    grepl("^KLF", tf, ignore.case = TRUE) ~ "KLF family",
    grepl("^SP[0-9]", tf, ignore.case = TRUE) ~ "SP family",
    grepl("^IRF", tf, ignore.case = TRUE) ~ "IRF/Interferon",
    grepl("^GATA", tf, ignore.case = TRUE) ~ "GATA family",
    grepl("^HNF", tf, ignore.case = TRUE) ~ "HNF family",
    grepl("^ZNF|^ZFP|^ZSCAN|^ZBTB", tf, ignore.case = TRUE) ~ "Zinc Finger",
    grepl("^HOX", tf, ignore.case = TRUE) ~ "HOX family",
    grepl("^SOX", tf, ignore.case = TRUE) ~ "SOX family",
    grepl("^PAX", tf, ignore.case = TRUE) ~ "PAX family",
    grepl("^E2F", tf, ignore.case = TRUE) ~ "E2F family",
    grepl("^MYC|^MAX", tf, ignore.case = TRUE) ~ "MYC family",
    grepl("^TP53|^TP63|^TP73", tf, ignore.case = TRUE) ~ "p53 family",
    grepl("^CEBP", tf, ignore.case = TRUE) ~ "C/EBP family",
    grepl("^NRF|^NFE2", tf, ignore.case = TRUE) ~ "NRF/Oxidative",
    grepl("^HIF|^EPAS", tf, ignore.case = TRUE) ~ "HIF/Hypoxia",
    grepl("^GLI", tf, ignore.case = TRUE) ~ "GLI/Hedgehog",
    grepl("^NOTCH|^RBPJ", tf, ignore.case = TRUE) ~ "Notch pathway",
    grepl("^TCF|^LEF", tf, ignore.case = TRUE) ~ "TCF/LEF/Wnt",
    grepl("^ETS|^ELK|^ELF|^ERG", tf, ignore.case = TRUE) ~ "ETS family",
    TRUE ~ "Other"
  )
}

# Add category column
ranks_wide <- ranks_wide %>%
  mutate(Category = sapply(TF, categorize_tf)) %>%
  select(TF, Category, everything())

scores_wide <- scores_wide %>%
  mutate(Category = sapply(TF, categorize_tf)) %>%
  select(TF, Category, everything())

# -----------------------------------------------------------------------------
# ADD SUMMARY STATISTICS
# -----------------------------------------------------------------------------

# Calculate best (minimum) rank across downregulated analyses
ranks_wide <- ranks_wide %>%
  rowwise() %>%
  mutate(
    Best_Rank_Down = min(c(down_3hr, down_6hr, down_12hr, down_24hr, 
                           peak_down, early_transient_down, sustained_down, late_only_down), 
                         na.rm = TRUE),
    Best_Rank_Up = min(c(up_3hr, up_6hr, up_12hr, up_24hr,
                         peak_up, early_transient_up, sustained_up, late_only_up),
                       na.rm = TRUE),
    Best_Rank_Overall = min(c_across(down_3hr:mixed_response), na.rm = TRUE)
  ) %>%
  ungroup() %>%
  mutate(
    Best_Rank_Down = ifelse(is.infinite(Best_Rank_Down), NA, Best_Rank_Down),
    Best_Rank_Up = ifelse(is.infinite(Best_Rank_Up), NA, Best_Rank_Up),
    Best_Rank_Overall = ifelse(is.infinite(Best_Rank_Overall), NA, Best_Rank_Overall)
  )

# Sort by best overall rank
ranks_wide <- ranks_wide %>%
  arrange(Best_Rank_Overall)

# -----------------------------------------------------------------------------
# SAVE OUTPUT FILES
# -----------------------------------------------------------------------------

cat("=== Saving Output Files ===\n\n")

# 1. Complete ranks table - ALL 1632 TFs
write.csv(ranks_wide, file.path(PIPE, "ChEA3_Complete_TF_Ranks_Wide.csv"), row.names = FALSE)
cat("Saved: ChEA3_Complete_TF_Ranks_Wide.csv (", nrow(ranks_wide), " TFs x ", ncol(ranks_wide), " columns)\n")

# 2. Complete scores table - ALL 1632 TFs
write.csv(scores_wide, file.path(PIPE, "ChEA3_Complete_TF_Scores_Wide.csv"), row.names = FALSE)
cat("Saved: ChEA3_Complete_TF_Scores_Wide.csv\n")

# 3. Long format for alternative viewing
write.csv(combined_long, file.path(PIPE, "ChEA3_Complete_TF_Long_Format.csv"), row.names = FALSE)
cat("Saved: ChEA3_Complete_TF_Long_Format.csv\n")

# -----------------------------------------------------------------------------
# CREATE FILTERED TABLES FOR KEY TF CATEGORIES
# -----------------------------------------------------------------------------

cat("\n=== Creating Category-Specific Tables ===\n\n")

# Mechanosensing TFs (TEAD + SRF)
mechanosensing_tfs <- ranks_wide %>%
  filter(Category %in% c("YAP/TEAD", "MRTF/SRF"))
write.csv(mechanosensing_tfs, file.path(PIPE, "ChEA3_Mechanosensing_TFs.csv"), row.names = FALSE)
cat("Saved: ChEA3_Mechanosensing_TFs.csv (", nrow(mechanosensing_tfs), " TFs)\n")

# Immediate-early / Stress response TFs
ieg_tfs <- ranks_wide %>%
  filter(Category %in% c("AP-1", "Immediate-Early", "ATF/Stress", "NR4A/Orphan"))
write.csv(ieg_tfs, file.path(PIPE, "ChEA3_IEG_Stress_TFs.csv"), row.names = FALSE)
cat("Saved: ChEA3_IEG_Stress_TFs.csv (", nrow(ieg_tfs), " TFs)\n")

# FOX family TFs
fox_tfs <- ranks_wide %>%
  filter(Category == "FOX family")
write.csv(fox_tfs, file.path(PIPE, "ChEA3_FOX_Family_TFs.csv"), row.names = FALSE)
cat("Saved: ChEA3_FOX_Family_TFs.csv (", nrow(fox_tfs), " TFs)\n")

# Top 100 TFs (by best overall rank)
top100_tfs <- ranks_wide %>%
  head(100)
write.csv(top100_tfs, file.path(PIPE, "ChEA3_Top100_TFs.csv"), row.names = FALSE)
cat("Saved: ChEA3_Top100_TFs.csv\n")

# -----------------------------------------------------------------------------
# CREATE GENE COUNT SUMMARY
# -----------------------------------------------------------------------------

gene_patterns <- read.csv(file.path(PIPE, "Gene_Temporal_Patterns.csv"), stringsAsFactors = FALSE)

pattern_summary <- gene_patterns %>%
  count(temporal_pattern) %>%
  arrange(desc(n)) %>%
  rename(Pattern = temporal_pattern, Gene_Count = n)

# Add gene list file info
gene_list_files <- data.frame(
  Analysis = c("down_3hr", "down_6hr", "down_12hr", "down_24hr",
               "up_3hr", "up_6hr", "up_12hr", "up_24hr",
               "peak_down", "peak_up",
               "early_transient_down", "early_transient_up",
               "sustained_down", "sustained_up",
               "late_only_down", "late_only_up",
               "mixed_response"),
  Gene_Count = c(
    nrow(all_results$down_3hr %>% head(1) %>% select(TF)),  # placeholder
    length(readLines(file.path(PIPE, "Genes_down_3hr.txt"))),
    length(readLines(file.path(PIPE, "Genes_down_6hr.txt"))),
    length(readLines(file.path(PIPE, "Genes_down_12hr.txt"))),
    length(readLines(file.path(PIPE, "Genes_down_24hr.txt"))),
    length(readLines(file.path(PIPE, "Genes_up_3hr.txt"))),
    length(readLines(file.path(PIPE, "Genes_up_6hr.txt"))),
    length(readLines(file.path(PIPE, "Genes_up_12hr.txt"))),
    length(readLines(file.path(PIPE, "Genes_up_24hr.txt"))),
    length(readLines(file.path(PIPE, "Genes_peak_down.txt"))),
    length(readLines(file.path(PIPE, "Genes_peak_up.txt"))),
    length(readLines(file.path(PIPE, "Genes_early_transient_down.txt"))),
    length(readLines(file.path(PIPE, "Genes_early_transient_up.txt"))),
    length(readLines(file.path(PIPE, "Genes_sustained_down.txt"))),
    length(readLines(file.path(PIPE, "Genes_sustained_up.txt"))),
    length(readLines(file.path(PIPE, "Genes_late_only_down.txt"))),
    length(readLines(file.path(PIPE, "Genes_late_only_up.txt"))),
    length(readLines(file.path(PIPE, "Genes_mixed_response.txt")))
  )[-1]  # Remove placeholder
)

# Actually read gene counts properly
gene_counts <- data.frame(
  Analysis = c("down_3hr", "down_6hr", "down_12hr", "down_24hr",
               "up_3hr", "up_6hr", "up_12hr", "up_24hr",
               "peak_down", "peak_up",
               "early_transient_down", "early_transient_up",
               "sustained_down", "sustained_up",
               "late_only_down", "late_only_up",
               "mixed_response"),
  stringsAsFactors = FALSE
)

gene_counts$Gene_Count <- sapply(gene_counts$Analysis, function(a) {
  file <- file.path(PIPE, paste0("Genes_", a, ".txt"))
  if (file.exists(file)) {
    return(length(readLines(file)))
  } else {
    return(NA)
  }
})

write.csv(gene_counts, file.path(PIPE, "ChEA3_Gene_Counts_Per_Analysis.csv"), row.names = FALSE)
cat("Saved: ChEA3_Gene_Counts_Per_Analysis.csv\n")

# -----------------------------------------------------------------------------
# PRINT SUMMARY
# -----------------------------------------------------------------------------

cat("\n")
cat("================================================================================\n")
cat("COMPLETE OUTPUT GENERATED!\n")
cat("================================================================================\n\n")

cat("MAIN FILES:\n")
cat("  1. ChEA3_Complete_TF_Ranks_Wide.csv   - ALL 1632 TFs, ranks across 17 analyses\n")
cat("  2. ChEA3_Complete_TF_Scores_Wide.csv  - ALL 1632 TFs, scores across 17 analyses\n")
cat("  3. ChEA3_Complete_TF_Long_Format.csv  - Long format (TF, Rank, Score, Analysis)\n\n")

cat("FILTERED TABLES:\n")
cat("  4. ChEA3_Mechanosensing_TFs.csv       - YAP/TEAD + MRTF/SRF only\n")
cat("  5. ChEA3_IEG_Stress_TFs.csv           - AP-1, EGR, ATF, NR4A families\n")
cat("  6. ChEA3_FOX_Family_TFs.csv           - All FOX TFs\n")
cat("  7. ChEA3_Top100_TFs.csv               - Top 100 by best overall rank\n")
cat("  8. ChEA3_Gene_Counts_Per_Analysis.csv - Gene counts for each analysis\n\n")

cat("================================================================================\n")
cat("COLUMN DESCRIPTIONS:\n")
cat("================================================================================\n")
cat("TF              - Transcription factor name\n")
cat("Category        - TF family/pathway classification\n")
cat("down_3hr        - Rank in downregulated genes at 3hr\n")
cat("down_6hr        - Rank in downregulated genes at 6hr\n")
cat("down_12hr       - Rank in downregulated genes at 12hr\n")
cat("down_24hr       - Rank in downregulated genes at 24hr (vs 24hr control)\n")
cat("up_3hr          - Rank in upregulated genes at 3hr\n")
cat("up_6hr          - Rank in upregulated genes at 6hr\n")
cat("up_12hr         - Rank in upregulated genes at 12hr\n")
cat("up_24hr         - Rank in upregulated genes at 24hr\n")
cat("peak_down       - Rank in genes with peak downregulation\n")
cat("peak_up         - Rank in genes with peak upregulation\n")
cat("early_transient_down - Rank in early transient downregulated\n")
cat("early_transient_up   - Rank in early transient upregulated\n")
cat("sustained_down  - Rank in sustained downregulated (core program)\n")
cat("sustained_up    - Rank in sustained upregulated\n")
cat("late_only_down  - Rank in late-only downregulated\n")
cat("late_only_up    - Rank in late-only upregulated\n")
cat("mixed_response  - Rank in mixed response genes\n")
cat("Best_Rank_Down  - Best (lowest) rank across all DOWN analyses\n")
cat("Best_Rank_Up    - Best (lowest) rank across all UP analyses\n")
cat("Best_Rank_Overall - Best (lowest) rank across ALL analyses\n")
cat("================================================================================\n")
```
