## Supplementary material for "Analysis of gene expression changes after Latrunculin B-induced actin depolymerization": Scripts: de_actin_genes.R

```
library(tidyverse)

# Run with the working directory set to Latb_rnaseq/scripts. Reads the outputs
# of deseq2_pipeline.R (run that first) from the local Pipeline_Outputs folder.
PIPE <- "Pipeline_Outputs"

cat("Creating 'core actin genes DE.csv'...\n\n")

################################################################################
# STEP 1: Define the 23 core actin genes
################################################################################

actin_genes <- data.frame(
  SYMBOL = c("ACTA1", "ACTA2", "ACTB", "ACTG1", "ACTN1", "ARPC2", "ARPC3", 
             "CAPZA1", "CAPZB", "CFL1", "DIAPH1", "FLNA", "FLNB", "FLNC", 
             "MYH10", "MYH9", "PFN1", "RAC1", "RHOA", "TMSB4X", "TWF1", 
             "VCL", "WAS"),
  ensembl_id = c("ENSG00000143632", "ENSG00000107796", "ENSG00000075624", 
                 "ENSG00000184009", "ENSG00000072110", "ENSG00000163466", 
                 "ENSG00000111229", "ENSG00000116489", "ENSG00000077549", 
                 "ENSG00000172757", "ENSG00000131504", "ENSG00000196924", 
                 "ENSG00000136068", "ENSG00000128591", "ENSG00000133026", 
                 "ENSG00000100345", "ENSG00000108518", "ENSG00000136238", 
                 "ENSG00000067560", "ENSG00000205542", "ENSG00000151239", 
                 "ENSG00000035403", "ENSG00000015285"),
  stringsAsFactors = FALSE
)

cat("Target genes:", nrow(actin_genes), "\n")

################################################################################
# STEP 2: Load normalized counts (all 16 samples)
################################################################################

cat("Loading normalized counts...\n")
normalized_counts <- read.csv(file.path(PIPE, "DESeq2_normalized_counts.csv"), row.names = 1)

# Filter for actin genes
actin_norm <- normalized_counts[rownames(normalized_counts) %in% actin_genes$ensembl_id, ]

cat("Found", nrow(actin_norm), "out of", nrow(actin_genes), "genes\n")

# Add gene symbols
gene_map <- setNames(actin_genes$SYMBOL, actin_genes$ensembl_id)
actin_norm <- data.frame(
  SYMBOL = gene_map[rownames(actin_norm)],
  ensembl_id = rownames(actin_norm),
  actin_norm,
  stringsAsFactors = FALSE
)

################################################################################
# STEP 3: Load all DESeq2 comparison results
################################################################################

cat("Loading DESeq2 results...\n")

comparisons <- c(
  "DESeq2_LatB_3hr_vs_Control_0hr.csv",
  "DESeq2_LatB_6hr_vs_Control_0hr.csv",
  "DESeq2_LatB_12hr_vs_Control_0hr.csv",
  "DESeq2_LatB_24hr_vs_Control_0hr.csv",
  "DESeq2_Control_24hr_vs_Control_0hr.csv",
  "DESeq2_LatB_24hr_vs_Control_24hr.csv"
)

# Initialize storage for DESeq2 results
log2fc_list <- list()
padj_list <- list()
basemean_list <- list()

for(comp_file in comparisons) {
  # Extract comparison name
  comp_name <- gsub("DESeq2_", "", gsub(".csv", "", comp_file))
  
  # Load results
  res <- read.csv(file.path(PIPE, comp_file), row.names = 1)

  # Filter for actin genes
  res_actin <- res[rownames(res) %in% actin_genes$ensembl_id, ]
  
  # Store log2FC
  log2fc_list[[paste0("log2FC_", comp_name)]] <- 
    res_actin[rownames(actin_norm)[actin_norm$ensembl_id %in% rownames(res_actin)], "log2FoldChange"]
  
  # Store padj
  padj_list[[paste0("padj_", comp_name)]] <- 
    res_actin[rownames(actin_norm)[actin_norm$ensembl_id %in% rownames(res_actin)], "padj"]
  
  # Store baseMean
  basemean_list[[paste0("baseMean_", comp_name)]] <- 
    res_actin[rownames(actin_norm)[actin_norm$ensembl_id %in% rownames(res_actin)], "baseMean"]
  
  cat("  Processed:", comp_name, "\n")
}

# Convert lists to data frames
log2fc_df <- data.frame(log2fc_list, row.names = actin_norm$ensembl_id)
padj_df <- data.frame(padj_list, row.names = actin_norm$ensembl_id)
basemean_df <- data.frame(basemean_list, row.names = actin_norm$ensembl_id)

################################################################################
# STEP 4: Combine everything into one comprehensive file
################################################################################

cat("\nCombining all data...\n")

# Start with normalized counts (has SYMBOL and ensembl_id)
comprehensive <- actin_norm

# Add log2FC columns
comprehensive <- cbind(comprehensive, log2fc_df)

# Add padj columns
comprehensive <- cbind(comprehensive, padj_df)

# Add baseMean columns
comprehensive <- cbind(comprehensive, basemean_df)

# Sort by gene symbol
comprehensive <- comprehensive[order(comprehensive$SYMBOL), ]

################################################################################
# STEP 5: Save the file
################################################################################

output_file <- file.path(PIPE, "core actin genes DE.csv")
write.csv(comprehensive, output_file, row.names = FALSE)

cat("\n")
cat("================================================================================\n")
cat("✓ FILE CREATED SUCCESSFULLY!\n")
cat("================================================================================\n")
cat("\nFile name:", output_file, "\n")
cat("Location:", getwd(), "\n")
cat("\nDimensions:", nrow(comprehensive), "genes ×", ncol(comprehensive), "columns\n")

cat("\nColumn structure:\n")
cat("  • Identifiers: SYMBOL, ensembl_id (2 columns)\n")
cat("  • Individual samples: 16 columns\n")
cat("  • log2 Fold Changes: 6 columns\n")
cat("  • Adjusted p-values: 6 columns\n")
cat("  • Base means: 6 columns\n")
cat("  • TOTAL:", ncol(comprehensive), "columns\n")

cat("\nGenes included:\n")
cat(" ", paste(comprehensive$SYMBOL, collapse = ", "), "\n")

cat("\n✓ DONE!\n")

################################################################################
# OPTIONAL: Preview the data
################################################################################

cat("\nPreview (first 5 genes, selected columns):\n")
preview_cols <- c("SYMBOL", "ensembl_id", "SRR30416730", "SRR30416723", 
                  "log2FC_LatB_3hr_vs_Control_0hr", "padj_LatB_3hr_vs_Control_0hr")
print(comprehensive[1:5, preview_cols])

cat("\n================================================================================\n")
```
