## Supplementary material for "Analysis of gene expression changes after Latrunculin B-induced actin depolymerization": Scripts: de_all_genes.R

```
library(tidyverse)

# Install biomaRt if not already installed
if (!requireNamespace("BiocManager", quietly = TRUE))
  install.packages("BiocManager")
if (!requireNamespace("biomaRt", quietly = TRUE))
  BiocManager::install("biomaRt")

library(biomaRt)

# Run with the working directory set to Latb_rnaseq/scripts. Reads the outputs
# of deseq2_pipeline.R (run that first) from the local Pipeline_Outputs folder.
PIPE <- "Pipeline_Outputs"

cat("\n")
cat("================================================================================\n")
cat("CREATING COMPREHENSIVE FILE FOR ALL GENES\n")
cat("================================================================================\n\n")

################################################################################
# STEP 1: Load normalized counts (all genes, all 16 samples)
################################################################################

cat("STEP 1: Loading normalized counts...\n")
normalized_counts <- read.csv(file.path(PIPE, "DESeq2_normalized_counts.csv"), row.names = 1)

cat("  Loaded:", nrow(normalized_counts), "genes ×", ncol(normalized_counts), "samples\n")
cat("  First few Ensembl IDs:", paste(head(rownames(normalized_counts), 3), collapse = ", "), "...\n\n")

################################################################################
# STEP 2: Get gene symbols from Ensembl using biomaRt
################################################################################

cat("STEP 2: Querying Ensembl database for gene symbols...\n")
cat("  This may take 2-5 minutes for ~28,000 genes...\n\n")

# Get all Ensembl IDs
ensembl_ids <- rownames(normalized_counts)

# Connect to Ensembl
cat("  Connecting to Ensembl database...\n")
ensembl <- useEnsembl(biomart = "genes", dataset = "hsapiens_gene_ensembl")

cat("  Querying gene annotations...\n")
# Query for gene symbols, gene names, and biotypes
gene_annotations <- getBM(
  attributes = c('ensembl_gene_id', 'external_gene_name', 'description', 'gene_biotype'),
  filters = 'ensembl_gene_id',
  values = ensembl_ids,
  mart = ensembl
)

cat("  Retrieved annotations for", nrow(gene_annotations), "genes\n")

# Create a mapping from ensembl_id to gene_symbol
gene_map <- setNames(gene_annotations$external_gene_name, gene_annotations$ensembl_gene_id)

# For genes without symbols, use the Ensembl ID
missing_symbols <- ensembl_ids[!ensembl_ids %in% names(gene_map)]
if(length(missing_symbols) > 0) {
  cat("  Warning:", length(missing_symbols), "genes without gene symbols (will use Ensembl ID)\n")
  gene_map_missing <- setNames(missing_symbols, missing_symbols)
  gene_map <- c(gene_map, gene_map_missing)
}

cat("  Total genes mapped:", length(gene_map), "\n\n")

################################################################################
# STEP 3: Create base dataframe with gene info and normalized counts
################################################################################

cat("STEP 3: Creating base dataframe...\n")

# Create dataframe with gene identifiers and normalized counts
all_genes_data <- data.frame(
  ensembl_id = rownames(normalized_counts),
  SYMBOL = gene_map[rownames(normalized_counts)],
  normalized_counts,
  stringsAsFactors = FALSE,
  check.names = FALSE
)

# Add gene description if available
description_map <- setNames(gene_annotations$description, gene_annotations$ensembl_gene_id)
all_genes_data$description <- description_map[all_genes_data$ensembl_id]

# Add gene biotype if available
biotype_map <- setNames(gene_annotations$gene_biotype, gene_annotations$ensembl_gene_id)
all_genes_data$gene_biotype <- biotype_map[all_genes_data$ensembl_id]

cat("  Base dataframe created:", nrow(all_genes_data), "genes\n\n")

################################################################################
# STEP 4: Load all DESeq2 comparison results
################################################################################

cat("STEP 4: Loading DESeq2 results for all comparisons...\n")

comparisons <- c(
  "DESeq2_LatB_3hr_vs_Control_0hr.csv",
  "DESeq2_LatB_6hr_vs_Control_0hr.csv",
  "DESeq2_LatB_12hr_vs_Control_0hr.csv",
  "DESeq2_LatB_24hr_vs_Control_0hr.csv",
  "DESeq2_Control_24hr_vs_Control_0hr.csv",
  "DESeq2_LatB_24hr_vs_Control_24hr.csv"
)

# Initialize lists to store results
all_log2fc <- list()
all_padj <- list()
all_basemean <- list()

for(comp_file in comparisons) {
  comp_name <- gsub("DESeq2_", "", gsub(".csv", "", comp_file))
  
  cat("  Loading:", comp_name, "\n")
  
  # Load results
  res <- read.csv(file.path(PIPE, comp_file), row.names = 1)

  # Match to our gene order
  res_matched <- res[match(all_genes_data$ensembl_id, rownames(res)), ]
  
  # Store results
  all_log2fc[[comp_name]] <- res_matched$log2FoldChange
  all_padj[[comp_name]] <- res_matched$padj
  all_basemean[[comp_name]] <- res_matched$baseMean
}

cat("  All comparisons loaded\n\n")

################################################################################
# STEP 5: Combine everything into comprehensive dataframe
################################################################################

cat("STEP 5: Combining all data...\n")

# Convert lists to dataframes
log2fc_df <- as.data.frame(all_log2fc)
colnames(log2fc_df) <- paste0("log2FC_", names(all_log2fc))

padj_df <- as.data.frame(all_padj)
colnames(padj_df) <- paste0("padj_", names(all_padj))

basemean_df <- as.data.frame(all_basemean)
colnames(basemean_df) <- paste0("baseMean_", names(all_basemean))

# Combine everything
comprehensive <- cbind(all_genes_data, log2fc_df, padj_df, basemean_df)

# Reorder columns for better organization
sample_cols <- colnames(normalized_counts)
identifier_cols <- c("SYMBOL", "ensembl_id", "description", "gene_biotype")
log2fc_cols <- colnames(log2fc_df)
padj_cols <- colnames(padj_df)
basemean_cols <- colnames(basemean_df)

comprehensive <- comprehensive[, c(identifier_cols, sample_cols, log2fc_cols, padj_cols, basemean_cols)]

cat("  Combined dataframe:", nrow(comprehensive), "genes ×", ncol(comprehensive), "columns\n\n")

################################################################################
# STEP 6: Save the comprehensive file
################################################################################

cat("STEP 6: Saving file...\n")

output_file <- file.path(PIPE, "all genes DE.csv")
write.csv(comprehensive, output_file, row.names = FALSE)

cat("  File saved:", output_file, "\n\n")

################################################################################
# STEP 7: Create summary statistics
################################################################################

cat("================================================================================\n")
cat("FILE CREATION SUMMARY\n")
cat("================================================================================\n\n")

cat("Output file:", output_file, "\n")
cat("Location:", getwd(), "\n\n")

cat("Dimensions:\n")
cat("  Genes:", nrow(comprehensive), "\n")
cat("  Columns:", ncol(comprehensive), "\n\n")

cat("Column breakdown:\n")
cat("  • Gene identifiers: 4 columns (SYMBOL, ensembl_id, description, gene_biotype)\n")
cat("  • Normalized counts: 16 columns (all samples)\n")
cat("  • log2 Fold Changes: 6 columns (all comparisons)\n")
cat("  • Adjusted p-values: 6 columns (all comparisons)\n")
cat("  • Base means: 6 columns (all comparisons)\n")
cat("  • TOTAL:", ncol(comprehensive), "columns\n\n")

# Count genes with symbols
genes_with_symbols <- sum(!is.na(comprehensive$SYMBOL) & comprehensive$SYMBOL != "" & 
                            comprehensive$SYMBOL != comprehensive$ensembl_id)
cat("Gene annotations:\n")
cat("  Genes with symbols:", genes_with_symbols, 
    sprintf("(%.1f%%)\n", 100 * genes_with_symbols / nrow(comprehensive)))
cat("  Genes without symbols:", nrow(comprehensive) - genes_with_symbols, 
    "(Ensembl ID used)\n\n")

# Sample comparisons included
cat("Comparisons included:\n")
for(comp in names(all_log2fc)) {
  cat("  •", comp, "\n")
}

################################################################################
# STEP 8: Create preview and summary statistics
################################################################################

cat("\n================================================================================\n")
cat("DATA PREVIEW (First 5 genes)\n")
cat("================================================================================\n\n")

preview_cols <- c("SYMBOL", "ensembl_id", "SRR30416730", "SRR30416723",
                  "log2FC_LatB_3hr_vs_Control_0hr", "padj_LatB_3hr_vs_Control_0hr")
print(comprehensive[1:5, preview_cols])

cat("\n================================================================================\n")
cat("SIGNIFICANCE SUMMARY ACROSS ALL GENES\n")
cat("================================================================================\n\n")

for(comp in names(all_log2fc)) {
  padj_col <- paste0("padj_", comp)
  log2fc_col <- paste0("log2FC_", comp)
  
  n_sig <- sum(comprehensive[[padj_col]] < 0.05, na.rm = TRUE)
  n_up <- sum(comprehensive[[padj_col]] < 0.05 & comprehensive[[log2fc_col]] > 0, na.rm = TRUE)
  n_down <- sum(comprehensive[[padj_col]] < 0.05 & comprehensive[[log2fc_col]] < 0, na.rm = TRUE)
  
  cat(sprintf("%-40s %5d DEGs (%d UP, %d DOWN)\n", comp, n_sig, n_up, n_down))
}

cat("\n================================================================================\n")
cat("✓ FILE CREATED SUCCESSFULLY!\n")
cat("================================================================================\n\n")

cat("Next steps:\n")
cat("  1. Open 'all genes DE.csv' in Excel or R\n")
cat("  2. Filter by SYMBOL to find genes of interest\n")
cat("  3. Filter by padj < 0.05 for significant genes\n")
cat("  4. Use log2FC columns for expression changes\n\n")

cat("File contains complete differential expression data for all", nrow(comprehensive), "genes!\n\n")

################################################################################
# OPTIONAL: Save a filtered version with only protein-coding genes
################################################################################

cat("================================================================================\n")
cat("BONUS: Creating filtered version (protein-coding genes only)\n")
cat("================================================================================\n\n")

if("gene_biotype" %in% colnames(comprehensive)) {
  protein_coding <- comprehensive[comprehensive$gene_biotype == "protein_coding", ]
  
  if(nrow(protein_coding) > 0) {
    output_file_pc <- file.path(PIPE, "all genes DE - protein coding only.csv")
    write.csv(protein_coding, output_file_pc, row.names = FALSE)
    
    cat("✓ Created:", output_file_pc, "\n")
    cat("  Contains:", nrow(protein_coding), "protein-coding genes\n\n")
  }
}

cat("================================================================================\n")
cat("✓ ALL DONE!\n")
cat("================================================================================\n\n")

# Print session info for reproducibility
cat("R session info:\n")
print(sessionInfo())
```
