## Supplementary material for "Analysis of gene expression changes after Latrunculin B-induced actin depolymerization": Scripts: decoupler_tf_activity.R

```
# =============================================================================
# DECOUPLER TF ACTIVITY INFERENCE ANALYSIS (FINAL FIXED VERSION)
# GSE275775 LatB RNA-seq Time Course
# Using CollecTRI/DoRothEA regulons with decoupleR
# =============================================================================
#
# FIXES:
# 1. ENSEMBL to gene symbol conversion
# 2. Explicit dplyr:: namespace to avoid AnnotationDbi conflict
#
# =============================================================================

# -----------------------------------------------------------------------------
# INSTALL REQUIRED PACKAGES
# -----------------------------------------------------------------------------

if (!requireNamespace("BiocManager", quietly = TRUE)) {
  install.packages("BiocManager")
}

if (!requireNamespace("decoupleR", quietly = TRUE)) {
  BiocManager::install("decoupleR")
}

if (!requireNamespace("OmnipathR", quietly = TRUE)) {
  BiocManager::install("OmnipathR")
}

if (!requireNamespace("org.Hs.eg.db", quietly = TRUE)) {
  BiocManager::install("org.Hs.eg.db")
}

if (!requireNamespace("AnnotationDbi", quietly = TRUE)) {
  BiocManager::install("AnnotationDbi")
}

required_packages <- c("dplyr", "tidyr", "ggplot2", "pheatmap", 
                       "ggrepel", "tibble", "RColorBrewer")

for (pkg in required_packages) {
  if (!requireNamespace(pkg, quietly = TRUE)) {
    install.packages(pkg)
  }
}

# -----------------------------------------------------------------------------
# LOAD LIBRARIES
# -----------------------------------------------------------------------------

library(decoupleR)
library(OmnipathR)
library(dplyr)
library(tidyr)
library(tibble)
library(ggplot2)
library(pheatmap)
library(ggrepel)
library(RColorBrewer)
library(org.Hs.eg.db)
library(AnnotationDbi)

# -----------------------------------------------------------------------------
# SET WORKING DIRECTORY AND LOAD DATA
# -----------------------------------------------------------------------------

# Run with the working directory set to Latb_rnaseq/scripts. Reads the outputs
# of deseq2_pipeline.R (run that first) from the local Pipeline_Outputs folder.
PIPE <- "Pipeline_Outputs"

cat("=============================================================================\n")
cat("DECOUPLER TF ACTIVITY INFERENCE ANALYSIS\n")
cat("GSE275775 LatB RNA-seq Time Course\n")
cat("=============================================================================\n\n")

# Load DESeq2 results - ALL COMPARED TO CONTROL_0HR
cat("Loading DESeq2 results (all vs Control_0hr)...\n")

deseq_3hr <- read.csv(file.path(PIPE, "DESeq2_LatB_3hr_vs_Control_0hr.csv"), row.names = 1)
deseq_6hr <- read.csv(file.path(PIPE, "DESeq2_LatB_6hr_vs_Control_0hr.csv"), row.names = 1)
deseq_12hr <- read.csv(file.path(PIPE, "DESeq2_LatB_12hr_vs_Control_0hr.csv"), row.names = 1)
deseq_24hr <- read.csv(file.path(PIPE, "DESeq2_LatB_24hr_vs_Control_0hr.csv"), row.names = 1)

cat("  - 3hr: ", nrow(deseq_3hr), " genes\n")
cat("  - 6hr: ", nrow(deseq_6hr), " genes\n")
cat("  - 12hr: ", nrow(deseq_12hr), " genes\n")
cat("  - 24hr: ", nrow(deseq_24hr), " genes\n")

# -----------------------------------------------------------------------------
# CONVERT ENSEMBL IDs TO GENE SYMBOLS
# -----------------------------------------------------------------------------

cat("\n=== Converting ENSEMBL IDs to Gene Symbols ===\n\n")

all_ensembl_ids <- rownames(deseq_3hr)
cat("Total ENSEMBL IDs: ", length(all_ensembl_ids), "\n")

# Remove version numbers
ensembl_clean <- gsub("\\..*", "", all_ensembl_ids)

# Map ENSEMBL to Symbol
cat("Mapping ENSEMBL IDs to gene symbols...\n")
gene_mapping <- AnnotationDbi::select(org.Hs.eg.db,
                                      keys = ensembl_clean,
                                      columns = c("SYMBOL"),
                                      keytype = "ENSEMBL")

# Remove duplicates
gene_mapping <- gene_mapping[!duplicated(gene_mapping$ENSEMBL), ]

# Create lookup vector
ensembl_to_symbol <- setNames(gene_mapping$SYMBOL, gene_mapping$ENSEMBL)

n_mapped <- sum(!is.na(ensembl_to_symbol))
cat("  Successfully mapped: ", n_mapped, " genes (", 
    round(100 * n_mapped / length(ensembl_clean), 1), "%)\n")

# Function to convert DESeq2 results to use gene symbols
convert_to_symbols <- function(deseq_result, ensembl_to_symbol) {
  ensembl_clean <- gsub("\\..*", "", rownames(deseq_result))
  symbols <- ensembl_to_symbol[ensembl_clean]
  
  deseq_result$ENSEMBL <- rownames(deseq_result)
  deseq_result$SYMBOL <- symbols
  
  # Remove unmapped
  
  deseq_result <- deseq_result[!is.na(deseq_result$SYMBOL), ]
  
  # Handle duplicates - use dplyr:: explicitly to avoid AnnotationDbi conflict
  deseq_result <- deseq_result %>%
    dplyr::group_by(SYMBOL) %>%
    dplyr::slice_max(order_by = baseMean, n = 1, with_ties = FALSE) %>%
    dplyr::ungroup() %>%
    as.data.frame()
  
  rownames(deseq_result) <- deseq_result$SYMBOL
  return(deseq_result)
}

# Convert all timepoints
cat("Converting DESeq2 results to gene symbols...\n")
deseq_3hr_sym <- convert_to_symbols(deseq_3hr, ensembl_to_symbol)
deseq_6hr_sym <- convert_to_symbols(deseq_6hr, ensembl_to_symbol)
deseq_12hr_sym <- convert_to_symbols(deseq_12hr, ensembl_to_symbol)
deseq_24hr_sym <- convert_to_symbols(deseq_24hr, ensembl_to_symbol)

cat("  - 3hr with symbols: ", nrow(deseq_3hr_sym), " genes\n")
cat("  - 6hr with symbols: ", nrow(deseq_6hr_sym), " genes\n")
cat("  - 12hr with symbols: ", nrow(deseq_12hr_sym), " genes\n")
cat("  - 24hr with symbols: ", nrow(deseq_24hr_sym), " genes\n")

# Verify key genes
cat("\nVerifying key genes are mapped:\n")
key_genes <- c("ACTB", "GAPDH", "TEAD1", "SRF", "JUN", "FOS", "EGR1", "ACTA2", "MYH9")
for (gene in key_genes) {
  present <- gene %in% rownames(deseq_3hr_sym)
  cat("  ", gene, ": ", ifelse(present, "✓ Present", "✗ NOT FOUND"), "\n")
}

# -----------------------------------------------------------------------------
# TF CATEGORIZATION FUNCTION
# -----------------------------------------------------------------------------

categorize_tf <- function(tf) {
  dplyr::case_when(
    tf %in% c("SRF") ~ "MRTF/SRF",
    grepl("^TEAD[1-4]$", tf) ~ "YAP/TEAD",
    grepl("^FOX", tf) ~ "FOX family",
    grepl("^JUN|^FOS", tf) ~ "AP-1",
    grepl("^EGR", tf) ~ "Immediate-Early",
    grepl("^ATF", tf) ~ "ATF/Stress",
    grepl("^NR4A", tf) ~ "NR4A/Orphan",
    grepl("^STAT", tf) ~ "STAT/JAK",
    grepl("^SMAD", tf) ~ "SMAD/TGFb",
    grepl("^NFKB|^REL", tf) ~ "NF-kB",
    grepl("^KLF", tf) ~ "KLF family",
    grepl("^SP[0-9]", tf) ~ "SP family",
    grepl("^ZNF", tf) ~ "Zinc Finger",
    grepl("^IRF", tf) ~ "IRF/Interferon",
    grepl("^E2F", tf) ~ "E2F family",
    grepl("^MYC|^MAX", tf) ~ "MYC family",
    grepl("^TP53|^TP63|^TP73", tf) ~ "p53 family",
    grepl("^HIF|^EPAS", tf) ~ "HIF/Hypoxia",
    grepl("^GLI", tf) ~ "GLI/Hedgehog",
    grepl("^CEBP", tf) ~ "C/EBP family",
    TRUE ~ "Other"
  )
}

# -----------------------------------------------------------------------------
# RETRIEVE TF REGULON NETWORKS
# -----------------------------------------------------------------------------

cat("\n=== Retrieving TF Regulon Networks from OmniPath ===\n\n")

cat("Downloading CollecTRI network...\n")
collectri <- decoupleR::get_collectri(organism = 'human', split_complexes = FALSE)
cat("  CollecTRI: ", length(unique(collectri$source)), " TFs, ", 
    nrow(collectri), " interactions\n")

key_tfs_to_check <- c("TEAD1", "TEAD2", "TEAD3", "TEAD4", "SRF", 
                      "MYOCD", "JUN", "FOS", "EGR1", "ATF3")

cat("\nKey TFs in CollecTRI network:\n")
for (tf in key_tfs_to_check) {
  present <- tf %in% collectri$source
  n_targets <- sum(collectri$source == tf)
  status <- ifelse(present, paste0("✓ Present (", n_targets, " targets)"), "✗ NOT FOUND")
  cat("  ", tf, ": ", status, "\n")
}

cat("\nDownloading DoRothEA network (confidence A, B, C)...\n")
dorothea <- decoupleR::get_dorothea(organism = 'human', levels = c('A', 'B', 'C'))
cat("  DoRothEA ABC: ", length(unique(dorothea$source)), " TFs, ", 
    nrow(dorothea), " interactions\n")

# Save networks
write.csv(collectri, file.path(PIPE, "CollecTRI_Network.csv"), row.names = FALSE)
write.csv(dorothea, file.path(PIPE, "DoRothEA_ABC_Network.csv"), row.names = FALSE)

# -----------------------------------------------------------------------------
# PREPARE INPUT MATRICES (NOW WITH GENE SYMBOLS)
# -----------------------------------------------------------------------------

cat("\n=== Preparing Input Matrices (Gene Symbols) ===\n\n")

# FIXED: Use base R instead of dplyr::select to avoid namespace issues
prepare_stat_matrix <- function(deseq_result, contrast_name) {
  # Remove NA stat values
  deseq_clean <- deseq_result[!is.na(deseq_result$stat), ]
  # Extract just the stat column as a data frame
  stat_df <- data.frame(stat = deseq_clean$stat, row.names = rownames(deseq_clean))
  colnames(stat_df) <- contrast_name
  return(stat_df)
}

stat_3hr <- prepare_stat_matrix(deseq_3hr_sym, "LatB_3hr")
stat_6hr <- prepare_stat_matrix(deseq_6hr_sym, "LatB_6hr")
stat_12hr <- prepare_stat_matrix(deseq_12hr_sym, "LatB_12hr")
stat_24hr <- prepare_stat_matrix(deseq_24hr_sym, "LatB_24hr")

cat("Stat matrices created:\n")
cat("  - 3hr: ", nrow(stat_3hr), " genes\n")
cat("  - 6hr: ", nrow(stat_6hr), " genes\n")
cat("  - 12hr: ", nrow(stat_12hr), " genes\n")
cat("  - 24hr: ", nrow(stat_24hr), " genes\n")

# Find common genes
common_genes <- Reduce(intersect, list(rownames(stat_3hr), rownames(stat_6hr), 
                                       rownames(stat_12hr), rownames(stat_24hr)))
cat("\nCommon genes across all timepoints: ", length(common_genes), "\n")

# Verify these are gene symbols not ENSEMBL IDs
cat("Sample gene names: ", paste(head(common_genes, 10), collapse = ", "), "\n")

# Create combined stat matrix
stat_matrix <- cbind(
  stat_3hr[common_genes, , drop = FALSE],
  stat_6hr[common_genes, , drop = FALSE],
  stat_12hr[common_genes, , drop = FALSE],
  stat_24hr[common_genes, , drop = FALSE]
)
stat_matrix <- as.matrix(stat_matrix)
cat("\nStat matrix dimensions: ", nrow(stat_matrix), " genes x ", ncol(stat_matrix), " contrasts\n")

# Check overlap with network
network_targets <- unique(collectri$target)
overlap <- sum(rownames(stat_matrix) %in% network_targets)
cat("Genes overlapping with CollecTRI network: ", overlap, " (", 
    round(100 * overlap / nrow(stat_matrix), 1), "% of our genes)\n")

if (overlap == 0) {
  stop("ERROR: No overlap between your genes and the network! Check gene symbol conversion.")
}

# -----------------------------------------------------------------------------
# RUN DECOUPLER - TF ACTIVITY INFERENCE
# -----------------------------------------------------------------------------

cat("\n=== Running decoupleR TF Activity Inference ===\n\n")

# Run ULM with CollecTRI
cat("Running ULM with CollecTRI network...\n")
tf_activities_collectri <- decoupleR::run_ulm(
  mat = stat_matrix,
  net = collectri,
  .source = 'source',
  .target = 'target',
  .mor = 'mor',
  minsize = 5
)
cat("  Results: ", nrow(tf_activities_collectri), " TF-contrast pairs\n")
cat("  TFs analyzed: ", length(unique(tf_activities_collectri$source)), "\n")

# Run ULM with DoRothEA
cat("\nRunning ULM with DoRothEA network...\n")
tf_activities_dorothea <- decoupleR::run_ulm(
  mat = stat_matrix,
  net = dorothea,
  .source = 'source',
  .target = 'target',
  .mor = 'mor',
  minsize = 5
)
cat("  Results: ", nrow(tf_activities_dorothea), " TF-contrast pairs\n")
cat("  TFs analyzed: ", length(unique(tf_activities_dorothea$source)), "\n")

# -----------------------------------------------------------------------------
# PROCESS AND FORMAT RESULTS
# -----------------------------------------------------------------------------

cat("\n=== Processing Results ===\n\n")

# Pivot to wide format - CollecTRI
tf_scores_wide <- tf_activities_collectri %>%
  dplyr::filter(statistic == "ulm") %>%
  dplyr::select(source, condition, score) %>%
  tidyr::pivot_wider(names_from = condition, values_from = score) %>%
  dplyr::rename(TF = source)

tf_pvals_wide <- tf_activities_collectri %>%
  dplyr::filter(statistic == "ulm") %>%
  dplyr::select(source, condition, p_value) %>%
  tidyr::pivot_wider(names_from = condition, values_from = p_value) %>%
  dplyr::rename(TF = source)

# Add summary statistics
tf_scores_wide <- tf_scores_wide %>%
  dplyr::rowwise() %>%
  dplyr::mutate(
    Mean_Score = mean(c(LatB_3hr, LatB_6hr, LatB_12hr, LatB_24hr), na.rm = TRUE),
    Min_Score = min(c(LatB_3hr, LatB_6hr, LatB_12hr, LatB_24hr), na.rm = TRUE),
    Max_Score = max(c(LatB_3hr, LatB_6hr, LatB_12hr, LatB_24hr), na.rm = TRUE),
    Score_Range = Max_Score - Min_Score
  ) %>%
  dplyr::ungroup() %>%
  dplyr::arrange(Mean_Score)

# Add categories
tf_scores_wide$Category <- sapply(tf_scores_wide$TF, categorize_tf)

# Reorder columns
tf_scores_wide <- tf_scores_wide %>%
  dplyr::select(TF, Category, LatB_3hr, LatB_6hr, LatB_12hr, LatB_24hr, 
                Mean_Score, Min_Score, Max_Score, Score_Range)

cat("Processed TF activity scores for ", nrow(tf_scores_wide), " TFs\n")

# Save results
write.csv(tf_scores_wide, file.path(PIPE, "DecoupleR_TF_Activities_CollecTRI.csv"), row.names = FALSE)
cat("Saved: DecoupleR_TF_Activities_CollecTRI.csv\n")

write.csv(tf_pvals_wide, file.path(PIPE, "DecoupleR_TF_Pvalues_CollecTRI.csv"), row.names = FALSE)
cat("Saved: DecoupleR_TF_Pvalues_CollecTRI.csv\n")

# DoRothEA results
tf_scores_dorothea <- tf_activities_dorothea %>%
  dplyr::filter(statistic == "ulm") %>%
  dplyr::select(source, condition, score) %>%
  tidyr::pivot_wider(names_from = condition, values_from = score) %>%
  dplyr::rename(TF = source) %>%
  dplyr::rowwise() %>%
  dplyr::mutate(Mean_Score = mean(c(LatB_3hr, LatB_6hr, LatB_12hr, LatB_24hr), na.rm = TRUE)) %>%
  dplyr::ungroup() %>%
  dplyr::arrange(Mean_Score)

tf_scores_dorothea$Category <- sapply(tf_scores_dorothea$TF, categorize_tf)
write.csv(tf_scores_dorothea, file.path(PIPE, "DecoupleR_TF_Activities_DoRothEA.csv"), row.names = FALSE)
cat("Saved: DecoupleR_TF_Activities_DoRothEA.csv\n")

# -----------------------------------------------------------------------------
# VISUALIZATIONS
# -----------------------------------------------------------------------------

cat("\n=== Creating Visualizations ===\n\n")

# Define category colors
category_colors <- c(
  "YAP/TEAD" = "#E41A1C",
  "MRTF/SRF" = "#FF7F00",
  "FOX family" = "#4DAF4A",
  "AP-1" = "#F781BF",
  "Immediate-Early" = "#984EA3",
  "ATF/Stress" = "#A65628",
  "NR4A/Orphan" = "#FFFF33",
  "STAT/JAK" = "#1B9E77",
  "SMAD/TGFb" = "#66C2A5",
  "NF-kB" = "#FC8D62",
  "KLF family" = "#D95F02",
  "SP family" = "#7570B3",
  "Zinc Finger" = "#377EB8",
  "IRF/Interferon" = "#8DA0CB",
  "E2F family" = "#E78AC3",
  "MYC family" = "#A6D854",
  "p53 family" = "#FFD92F",
  "HIF/Hypoxia" = "#E5C494",
  "GLI/Hedgehog" = "#B3B3B3",
  "C/EBP family" = "#CBD5E8",
  "Other" = "#999999"
)

# 1. BAR PLOT: Top Inactivated and Activated TFs
# -----------------------------------------------------------------
cat("Creating bar plot of top TFs...\n")

top_inactivated <- tf_scores_wide %>% dplyr::arrange(Mean_Score) %>% head(20)
top_activated <- tf_scores_wide %>% dplyr::arrange(dplyr::desc(Mean_Score)) %>% head(20)

top_tfs <- dplyr::bind_rows(top_inactivated, top_activated) %>%
  dplyr::distinct(TF, .keep_all = TRUE) %>%
  dplyr::mutate(TF = factor(TF, levels = TF[order(Mean_Score)]))

p1 <- ggplot(top_tfs, aes(x = TF, y = Mean_Score, fill = Category)) +
  geom_bar(stat = "identity") +
  geom_hline(yintercept = 0, linetype = "dashed", color = "gray50") +
  coord_flip() +
  scale_fill_manual(values = category_colors) +
  labs(
    title = "TF Activity Scores (decoupleR/CollecTRI)",
    subtitle = "Mean activity across LatB timepoints vs Control_0hr",
    x = NULL,
    y = "Activity Score (negative = inactivated, positive = activated)",
    fill = "TF Family"
  ) +
  theme_minimal(base_size = 11) +
  theme(plot.title = element_text(face = "bold"), legend.position = "right")

ggsave(file.path(PIPE, "DecoupleR_Top_TFs_BarPlot.png"), p1, width = 12, height = 10, dpi = 300)
ggsave(file.path(PIPE, "DecoupleR_Top_TFs_BarPlot.pdf"), p1, width = 12, height = 10)
cat("Saved: DecoupleR_Top_TFs_BarPlot.png/pdf\n")

# 2. HEATMAP: TF Activities Across Timepoints
# -----------------------------------------------------------------
cat("Creating heatmap of TF activities...\n")

top_variable <- tf_scores_wide %>% 
  dplyr::arrange(dplyr::desc(Score_Range)) %>% 
  head(50)

heatmap_data <- top_variable %>%
  dplyr::select(TF, LatB_3hr, LatB_6hr, LatB_12hr, LatB_24hr) %>%
  tibble::column_to_rownames("TF") %>%
  as.matrix()

row_annotation <- data.frame(
  Category = top_variable$Category,
  row.names = top_variable$TF
)

max_abs <- max(abs(heatmap_data), na.rm = TRUE)
breaks <- seq(-max_abs, max_abs, length.out = 101)

png(file.path(PIPE, "DecoupleR_TF_Heatmap_Top50.png"), width = 10, height = 14, units = "in", res = 300)
pheatmap(heatmap_data,
         color = colorRampPalette(c("blue", "white", "red"))(100),
         breaks = breaks,
         cluster_rows = TRUE,
         cluster_cols = FALSE,
         show_rownames = TRUE,
         show_colnames = TRUE,
         annotation_row = row_annotation,
         annotation_colors = list(Category = category_colors),
         fontsize_row = 8,
         fontsize_col = 10,
         main = "TF Activity Scores Across LatB Timepoints (vs Control_0hr)\n(Blue = Inactivated, Red = Activated)")
dev.off()

pdf(file.path(PIPE, "DecoupleR_TF_Heatmap_Top50.pdf"), width = 10, height = 14)
pheatmap(heatmap_data,
         color = colorRampPalette(c("blue", "white", "red"))(100),
         breaks = breaks,
         cluster_rows = TRUE,
         cluster_cols = FALSE,
         show_rownames = TRUE,
         show_colnames = TRUE,
         annotation_row = row_annotation,
         annotation_colors = list(Category = category_colors),
         fontsize_row = 8,
         fontsize_col = 10,
         main = "TF Activity Scores Across LatB Timepoints (vs Control_0hr)\n(Blue = Inactivated, Red = Activated)")
dev.off()
cat("Saved: DecoupleR_TF_Heatmap_Top50.png/pdf\n")

# 3. MECHANOSENSING TFs: TEAD and SRF
# -----------------------------------------------------------------
cat("Creating mechanosensing TFs plot...\n")

mechano_tfs_available <- c("TEAD1", "TEAD2", "TEAD3", "TEAD4", "SRF")
mechano_tfs_present <- mechano_tfs_available[mechano_tfs_available %in% tf_scores_wide$TF]
cat("  Mechanosensing TFs found: ", paste(mechano_tfs_present, collapse = ", "), "\n")

if (length(mechano_tfs_present) > 0) {
  mechano_data <- tf_scores_wide %>%
    dplyr::filter(TF %in% mechano_tfs_present) %>%
    dplyr::select(TF, Category, LatB_3hr, LatB_6hr, LatB_12hr, LatB_24hr) %>%
    tidyr::pivot_longer(cols = starts_with("LatB"), names_to = "Timepoint", values_to = "Activity") %>%
    dplyr::mutate(
      Timepoint = gsub("LatB_", "", Timepoint),
      Timepoint = factor(Timepoint, levels = c("3hr", "6hr", "12hr", "24hr"))
    )
  
  p2 <- ggplot(mechano_data, aes(x = Timepoint, y = Activity, color = TF, group = TF)) +
    geom_line(linewidth = 1.5) +
    geom_point(size = 4) +
    geom_hline(yintercept = 0, linetype = "dashed", color = "gray50") +
    scale_color_manual(values = c("TEAD1" = "#E41A1C", "TEAD2" = "#FF7F00", 
                                  "TEAD3" = "#4DAF4A", "TEAD4" = "#984EA3",
                                  "SRF" = "#377EB8")) +
    labs(
      title = "Mechanosensing TF Activities Over Time",
      subtitle = "YAP/TEAD and SRF (DNA-binding TFs only)",
      x = "Timepoint (vs Control_0hr)",
      y = "Activity Score",
      color = "TF"
    ) +
    theme_minimal(base_size = 12) +
    theme(plot.title = element_text(face = "bold"), legend.position = "right")
  
  ggsave(file.path(PIPE, "DecoupleR_Mechanosensing_TFs_Temporal.png"), p2, width = 10, height = 6, dpi = 300)
  ggsave(file.path(PIPE, "DecoupleR_Mechanosensing_TFs_Temporal.pdf"), p2, width = 10, height = 6)
  cat("Saved: DecoupleR_Mechanosensing_TFs_Temporal.png/pdf\n")
} else {
  cat("WARNING: No mechanosensing TFs found in results!\n")
}

# 4. VOLCANO-STYLE PLOT: Activity vs Significance (6hr)
# -----------------------------------------------------------------
cat("Creating volcano-style plot (6hr)...\n")

volcano_data <- tf_activities_collectri %>%
  dplyr::filter(statistic == "ulm", condition == "LatB_6hr") %>%
  dplyr::mutate(
    neg_log10_pval = -log10(p_value),
    significant = p_value < 0.05,
    Category = sapply(source, categorize_tf),
    label = ifelse(abs(score) > 3 | (significant & abs(score) > 2), source, "")
  )

p3 <- ggplot(volcano_data, aes(x = score, y = neg_log10_pval, color = Category)) +
  geom_point(alpha = 0.7, size = 2) +
  geom_hline(yintercept = -log10(0.05), linetype = "dashed", color = "gray50") +
  geom_vline(xintercept = c(-2, 2), linetype = "dashed", color = "gray50") +
  geom_text_repel(aes(label = label), size = 3, max.overlaps = 20) +
  scale_color_manual(values = category_colors) +
  labs(
    title = "TF Activity Volcano Plot - LatB 6hr vs Control_0hr",
    subtitle = "Activity score vs statistical significance",
    x = "Activity Score (negative = inactivated)",
    y = "-log10(p-value)",
    color = "TF Family"
  ) +
  theme_minimal(base_size = 11) +
  theme(plot.title = element_text(face = "bold"), legend.position = "right")

ggsave(file.path(PIPE, "DecoupleR_Volcano_6hr.png"), p3, width = 12, height = 10, dpi = 300)
ggsave(file.path(PIPE, "DecoupleR_Volcano_6hr.pdf"), p3, width = 12, height = 10)
cat("Saved: DecoupleR_Volcano_6hr.png/pdf\n")

# 5. TF FAMILY SUMMARY
# -----------------------------------------------------------------
cat("Creating TF family summary...\n")

family_summary <- tf_scores_wide %>%
  dplyr::group_by(Category) %>%
  dplyr::summarise(
    n_TFs = dplyr::n(),
    Mean_Activity = mean(Mean_Score, na.rm = TRUE),
    SD_Activity = sd(Mean_Score, na.rm = TRUE),
    .groups = "drop"
  ) %>%
  dplyr::arrange(Mean_Activity)

p4 <- ggplot(family_summary, aes(x = reorder(Category, Mean_Activity), y = Mean_Activity, fill = Category)) +
  geom_bar(stat = "identity") +
  geom_errorbar(aes(ymin = Mean_Activity - SD_Activity, ymax = Mean_Activity + SD_Activity), 
                width = 0.3) +
  geom_hline(yintercept = 0, linetype = "dashed") +
  coord_flip() +
  scale_fill_manual(values = category_colors) +
  labs(
    title = "TF Family Activity Summary",
    subtitle = "Mean activity score across all TFs in each family",
    x = NULL,
    y = "Mean Activity Score ± SD"
  ) +
  theme_minimal(base_size = 12) +
  theme(legend.position = "none")

ggsave(file.path(PIPE, "DecoupleR_Family_Summary.png"), p4, width = 10, height = 8, dpi = 300)
ggsave(file.path(PIPE, "DecoupleR_Family_Summary.pdf"), p4, width = 10, height = 8)
cat("Saved: DecoupleR_Family_Summary.png/pdf\n")

# -----------------------------------------------------------------------------
# KEY TF RESULTS TABLE
# -----------------------------------------------------------------------------

cat("\n=== Key TF Results ===\n\n")

key_tfs_list <- c("TEAD1", "TEAD2", "TEAD3", "TEAD4",
                  "SRF",
                  "JUN", "JUNB", "JUND", "FOS", "FOSB", "FOSL1", "FOSL2",
                  "EGR1", "EGR2", "EGR3", "EGR4",
                  "ATF1", "ATF2", "ATF3", "ATF4",
                  "NR4A1", "NR4A2", "NR4A3",
                  "STAT1", "STAT2", "STAT3",
                  "SMAD2", "SMAD3", "SMAD4",
                  "FOXO1", "FOXO3", "FOXF1", "FOXA1",
                  "KLF4", "KLF6", "SP1", "SP3")

key_tf_results <- tf_scores_wide %>%
  dplyr::filter(TF %in% key_tfs_list) %>%
  dplyr::arrange(Mean_Score)

cat("\nKey TF Activity Scores (decoupleR/CollecTRI):\n")
cat("==============================================\n")
print(key_tf_results, n = 50)

write.csv(key_tf_results, file.path(PIPE, "DecoupleR_Key_TFs_Results.csv"), row.names = FALSE)
cat("\nSaved: DecoupleR_Key_TFs_Results.csv\n")

# -----------------------------------------------------------------------------
# PRINT TOP RESULTS SUMMARY
# -----------------------------------------------------------------------------

cat("\n=== TOP 10 MOST INACTIVATED TFs (LatB Treatment) ===\n")
print(tf_scores_wide %>% dplyr::arrange(Mean_Score) %>% head(10) %>% 
        dplyr::select(TF, Category, Mean_Score, LatB_3hr, LatB_6hr, LatB_12hr, LatB_24hr))

cat("\n=== TOP 10 MOST ACTIVATED TFs (LatB Treatment) ===\n")
print(tf_scores_wide %>% dplyr::arrange(dplyr::desc(Mean_Score)) %>% head(10) %>% 
        dplyr::select(TF, Category, Mean_Score, LatB_3hr, LatB_6hr, LatB_12hr, LatB_24hr))

# -----------------------------------------------------------------------------
# FINAL SUMMARY
# -----------------------------------------------------------------------------

cat("\n")
cat("=============================================================================\n")
cat("ANALYSIS COMPLETE\n")
cat("=============================================================================\n\n")

cat("OUTPUT FILES:\n")
cat("  - DecoupleR_TF_Activities_CollecTRI.csv (main results)\n")
cat("  - DecoupleR_TF_Pvalues_CollecTRI.csv\n")
cat("  - DecoupleR_TF_Activities_DoRothEA.csv\n")
cat("  - DecoupleR_Key_TFs_Results.csv\n")
cat("  - DecoupleR_Top_TFs_BarPlot.png/pdf\n")
cat("  - DecoupleR_TF_Heatmap_Top50.png/pdf\n")
cat("  - DecoupleR_Mechanosensing_TFs_Temporal.png/pdf\n")
cat("  - DecoupleR_Volcano_6hr.png/pdf\n")
cat("  - DecoupleR_Family_Summary.png/pdf\n")
```
