## Supplementary material for "Analysis of gene expression changes after Latrunculin B-induced actin depolymerization": Scripts: deseq2_pipeline.R

```
# Load required libraries
library(DESeq2)
library(tidyverse)
library(pheatmap)
library(RColorBrewer)
library(ggplot2)

# Run with the working directory set to Latb_rnaseq/scripts (matching the
# suppfig_*.R convention). Raw per-replicate counts come from Supplementary
# Data File 1 (S1) - the original private counts matrix used condition-blind
# SRR accessions as column names; S1's count columns are labeled by
# condition/timepoint/replicate instead, so sample metadata below is parsed
# from those column names rather than hardcoded by SRR position.
SDF <- "../Supplementary_Data_Files"
dir.create("Pipeline_Outputs", showWarnings = FALSE, recursive = TRUE)

################################################################################
# STEP 1: LOAD RAW COUNT DATA
################################################################################

cat("\n")
cat("═══════════════════════════════════════════════════════════════════\n")
cat("STARTING DESEQ2 ANALYSIS\n")
cat("═══════════════════════════════════════════════════════════════════\n\n")

s1 <- read.csv(file.path(SDF, "S1_All_Genes_DE_Raw_Counts.csv"),
               row.names = "ensembl_id", check.names = FALSE)
count_cols <- grep("^(Control|LatB)_[0-9]+hr_rep[0-9]+$", colnames(s1), value = TRUE)
raw_counts <- s1[, count_cols]

# Verify data is integers
cat("✓ Checking data integrity...\n")
if(all(raw_counts == floor(raw_counts))) {
  cat("  ✓ Counts are integers - CORRECT\n")
} else {
  stop("  ✗ ERROR: Counts are not integers!")
}

cat("  ✓ Matrix dimensions:", nrow(raw_counts), "genes x", ncol(raw_counts), "samples\n\n")

################################################################################
# STEP 2: CREATE SAMPLE METADATA - CORRECTED
################################################################################

cn <- colnames(raw_counts)
sample_info <- data.frame(
  sample_id = cn,
  condition = sub("_.*$", "", cn),
  timepoint = as.numeric(sub("^.*_([0-9]+)hr_rep[0-9]+$", "\\1", cn)),
  replicate = as.numeric(sub("^.*_rep([0-9]+)$", "\\1", cn)),
  stringsAsFactors = FALSE
)

# Restore the original SRR accessions as column/sample names (verified 1:1
# against S1's condition/timepoint/replicate labels via Pearson r = 1.000,
# same check as suppfig_2.R) so downstream scripts that expect SRR-named
# sample columns (S5, S8, de_actin_genes.R, figure_2.R, figure_3.R, ...)
# still line up.
srr_map <- c(
  "Control_0hr_rep1" = "SRR30416730", "Control_0hr_rep2" = "SRR30416729", "Control_0hr_rep3" = "SRR30416728",
  "LatB_3hr_rep1" = "SRR30416725", "LatB_3hr_rep2" = "SRR30416724", "LatB_3hr_rep3" = "SRR30416723",
  "LatB_6hr_rep1" = "SRR30416722", "LatB_6hr_rep2" = "SRR30416721", "LatB_6hr_rep3" = "SRR30416720",
  "LatB_12hr_rep1" = "SRR30416719", "LatB_12hr_rep2" = "SRR30416718",
  "LatB_24hr_rep1" = "SRR30416716", "LatB_24hr_rep2" = "SRR30416717", "LatB_24hr_rep3" = "SRR30416715",
  "Control_24hr_rep1" = "SRR30416727", "Control_24hr_rep2" = "SRR30416726"
)
sample_info$sample_id <- unname(srr_map[cn])
colnames(raw_counts) <- unname(srr_map[cn])

rownames(sample_info) <- sample_info$sample_id

# CRITICAL FIX: Use numeric values for factor levels
sample_info$condition <- factor(sample_info$condition, 
                                levels = c("Control", "LatB"))
sample_info$timepoint <- factor(sample_info$timepoint,
                               levels = c(0, 3, 6, 12, 24))  # FIXED: numeric not character

# Combined factor for flexible comparisons
sample_info$group <- factor(paste(sample_info$condition, 
                                 sample_info$timepoint, 
                                 sep = "_"))

# CRITICAL CHECKS
cat("✓ Performing data integrity checks...\n")
if(!all(colnames(raw_counts) %in% rownames(sample_info))) {
  stop("  ✗ ERROR: Sample mismatch!")
}
if(!all(rownames(sample_info) == colnames(raw_counts))) {
  stop("  ✗ ERROR: Sample order mismatch!")
}
cat("  ✓ All samples verified\n")
cat("  ✓ Sample order correct\n\n")

cat("Sample metadata:\n")
print(sample_info)
cat("\n")

################################################################################
# STEP 3: PRE-FILTERING
################################################################################

cat("✓ Pre-filtering genes...\n")
cat("  Genes before filtering:", nrow(raw_counts), "\n")

# Filter: keep genes with at least 10 reads total
keep <- rowSums(raw_counts) >= 10
filtered_counts <- raw_counts[keep, ]

cat("  Genes after filtering:", nrow(filtered_counts), "\n")
cat("  Genes removed:", nrow(raw_counts) - nrow(filtered_counts), "\n\n")

################################################################################
# STEP 4: CREATE DESEQ2 OBJECT WITH OPTIMAL DESIGN
################################################################################

cat("✓ Creating DESeqDataSet...\n")

dds <- DESeqDataSetFromMatrix(
  countData = filtered_counts,
  colData = sample_info,
  design = ~ group
)

cat("  ✓ DESeqDataSet created\n")
cat("  Design formula: ~ group\n\n")

################################################################################
# STEP 5: QUALITY CONTROL BEFORE ANALYSIS
################################################################################

cat("✓ Running quality control checks...\n\n")

# QC1: Check library sizes
lib_sizes <- colSums(counts(dds))
cat("  Library sizes (millions of reads):\n")
print(round(lib_sizes / 1e6, 2))
cat("\n")

# QC2: Visualize library sizes
pdf("Pipeline_Outputs/QC_library_sizes.pdf", width = 10, height = 6)
par(mar = c(8, 4, 2, 1))
barplot(lib_sizes / 1e6, 
        las = 2, 
        col = ifelse(sample_info$condition == "Control", "steelblue", "coral"),
        main = "Library Sizes",
        ylab = "Millions of reads",
        names.arg = paste(sample_info$condition, sample_info$timepoint, sep = "_"))
legend("topright", 
       legend = c("Control", "LatB"), 
       fill = c("steelblue", "coral"))
dev.off()
cat("  ✓ Library size plot saved\n")

# QC3: Estimate size factors (for inspection)
dds <- estimateSizeFactors(dds)
cat("\n  Size factors (normalization factors):\n")
print(round(sizeFactors(dds), 3))
cat("\n")

# Check for outliers
if(any(sizeFactors(dds) < 0.5 | sizeFactors(dds) > 2)) {
  cat("  ⚠ WARNING: Some size factors are outside normal range (0.5-2.0)\n")
  cat("    This may indicate quality issues or very different library compositions\n\n")
}

################################################################################
# STEP 6: RUN DESEQ2 ANALYSIS
################################################################################

cat("═══════════════════════════════════════════════════════════════════\n")
cat("RUNNING DESEQ2 DIFFERENTIAL EXPRESSION ANALYSIS\n")
cat("═══════════════════════════════════════════════════════════════════\n\n")

cat("This will take 2-5 minutes...\n\n")

dds <- DESeq(dds)

cat("✓ DESeq2 analysis complete!\n\n")

# Check what comparisons are available
cat("Available result names:\n")
print(resultsNames(dds))
cat("\n")

################################################################################
# STEP 7: EXTRACT AND SAVE RESULTS - CORRECTED APPROACH
################################################################################

cat("═══════════════════════════════════════════════════════════════════\n")
cat("EXTRACTING RESULTS FOR ALL COMPARISONS\n")
cat("═══════════════════════════════════════════════════════════════════\n\n")

# Create output directory
dir.create("Pipeline_Outputs", showWarnings = FALSE, recursive = TRUE)

# Define all comparisons
comparisons <- list(
  list(name = "LatB_3hr_vs_Control_0hr", 
       contrast = c("group", "LatB_3", "Control_0")),
  list(name = "LatB_6hr_vs_Control_0hr", 
       contrast = c("group", "LatB_6", "Control_0")),
  list(name = "LatB_12hr_vs_Control_0hr", 
       contrast = c("group", "LatB_12", "Control_0")),
  list(name = "LatB_24hr_vs_Control_0hr", 
       contrast = c("group", "LatB_24", "Control_0")),
  list(name = "Control_24hr_vs_Control_0hr", 
       contrast = c("group", "Control_24", "Control_0")),
  list(name = "LatB_24hr_vs_Control_24hr", 
       contrast = c("group", "LatB_24", "Control_24"))
)

# Extract results for each comparison
all_results <- list()

for(comp in comparisons) {
  cat("Processing:", comp$name, "\n")
  
  # Extract results with proper settings
  res <- results(dds, 
                 contrast = comp$contrast,
                 alpha = 0.05,  # FDR threshold
                 pAdjustMethod = "BH")  # Benjamini-Hochberg
  
  # Order by adjusted p-value
  res <- res[order(res$padj), ]
  
  # Print summary
  cat("\nSummary for", comp$name, ":\n")
  print(summary(res))
  cat("\n")
  
  # Count significant genes
  sig_up <- sum(res$padj < 0.05 & res$log2FoldChange > 0, na.rm = TRUE)
  sig_down <- sum(res$padj < 0.05 & res$log2FoldChange < 0, na.rm = TRUE)
  cat(sprintf("  Significant genes (padj < 0.05): %d UP, %d DOWN\n", sig_up, sig_down))
  cat(sprintf("  Total significant: %d\n\n", sig_up + sig_down))
  
  # Save results
  write.csv(as.data.frame(res), 
            paste0("Pipeline_Outputs/DESeq2_", comp$name, ".csv"))
  
  # Store for later
  all_results[[comp$name]] <- res
}

################################################################################
# STEP 8: SAVE ADDITIONAL OUTPUTS
################################################################################

cat("═══════════════════════════════════════════════════════════════════\n")
cat("SAVING ADDITIONAL OUTPUTS\n")
cat("═══════════════════════════════════════════════════════════════════\n\n")

# Save DESeq2 object
saveRDS(dds, "Pipeline_Outputs/dds_object.rds")
cat("✓ DESeq2 object saved\n")

# Save normalized counts
normalized_counts <- counts(dds, normalized = TRUE)
write.csv(normalized_counts, "Pipeline_Outputs/DESeq2_normalized_counts.csv")
cat("✓ Normalized counts saved\n")

# Save size factors
size_factors_df <- data.frame(
  sample = names(sizeFactors(dds)), 
  size_factor = sizeFactors(dds),
  condition = sample_info$condition,
  timepoint = sample_info$timepoint
)
write.csv(size_factors_df, 
          "Pipeline_Outputs/DESeq2_size_factors.csv", 
          row.names = FALSE)
cat("✓ Size factors saved\n")

# Save dispersion estimates
write.csv(data.frame(
  gene = rownames(dds),
  dispersion = dispersions(dds)
), "Pipeline_Outputs/DESeq2_dispersions.csv", row.names = FALSE)
cat("✓ Dispersion estimates saved\n\n")

################################################################################
# STEP 9: QUALITY CONTROL VISUALIZATIONS
################################################################################

cat("═══════════════════════════════════════════════════════════════════\n")
cat("GENERATING QUALITY CONTROL VISUALIZATIONS\n")
cat("═══════════════════════════════════════════════════════════════════\n\n")

# QC4: Dispersion plot
pdf("Pipeline_Outputs/QC_dispersion_plot.pdf", width = 10, height = 8)
plotDispEsts(dds, main = "Dispersion Estimates")
dev.off()
cat("✓ Dispersion plot saved\n")

# QC5: PCA plot
vsd <- vst(dds, blind = FALSE)
pdf("Pipeline_Outputs/QC_PCA_plot.pdf", width = 10, height = 8)
plotPCA(vsd, intgroup = c("condition", "timepoint")) + 
  theme_bw() +
  ggtitle("PCA - Variance Stabilized Data") +
  theme(plot.title = element_text(hjust = 0.5))
dev.off()
cat("✓ PCA plot saved\n")

# QC6: Sample distance heatmap
sampleDists <- dist(t(assay(vsd)))
sampleDistMatrix <- as.matrix(sampleDists)
rownames(sampleDistMatrix) <- paste(vsd$condition, vsd$timepoint, sep = "_")
colnames(sampleDistMatrix) <- paste(vsd$condition, vsd$timepoint, sep = "_")
colors <- colorRampPalette(rev(brewer.pal(9, "Blues")))(255)

pdf("Pipeline_Outputs/QC_sample_distances.pdf", width = 10, height = 10)
pheatmap(sampleDistMatrix,
         clustering_distance_rows = sampleDists,
         clustering_distance_cols = sampleDists,
         col = colors,
         main = "Sample-to-Sample Distances")
dev.off()
cat("✓ Sample distance heatmap saved\n")

# QC7: MA plots for each comparison
for(comp in comparisons) {
  pdf(paste0("Pipeline_Outputs/MA_plot_", comp$name, ".pdf"), 
      width = 10, height = 8)
  plotMA(all_results[[comp$name]], 
         alpha = 0.05,
         main = paste("MA Plot:", comp$name),
         ylim = c(-5, 5))
  dev.off()
}
cat("✓ MA plots saved (6 comparisons)\n\n")

################################################################################
# STEP 10: CREATE SUMMARY REPORT
################################################################################

cat("═══════════════════════════════════════════════════════════════════\n")
cat("CREATING SUMMARY REPORT\n")
cat("═══════════════════════════════════════════════════════════════════\n\n")

# Create comprehensive summary
summary_df <- data.frame(
  Comparison = character(),
  Total_Genes = integer(),
  Sig_Total = integer(),
  Sig_Up = integer(),
  Sig_Down = integer(),
  stringsAsFactors = FALSE
)

for(comp in comparisons) {
  res <- all_results[[comp$name]]
  sig_up <- sum(res$padj < 0.05 & res$log2FoldChange > 0, na.rm = TRUE)
  sig_down <- sum(res$padj < 0.05 & res$log2FoldChange < 0, na.rm = TRUE)
  
  summary_df <- rbind(summary_df, data.frame(
    Comparison = comp$name,
    Total_Genes = nrow(res),
    Sig_Total = sig_up + sig_down,
    Sig_Up = sig_up,
    Sig_Down = sig_down
  ))
}

write.csv(summary_df, 
          "Pipeline_Outputs/SUMMARY_all_comparisons.csv", 
          row.names = FALSE)

cat("✓ Summary report created\n\n")
print(summary_df)
```
