## Supplementary material for "Analysis of gene expression changes after Latrunculin B-induced actin depolymerization": Scripts: deseq2_pooled.R

```
# =============================================================================
# SIMPLE POOLED DESEQ2 ANALYSIS
# All LatB (3hr+6hr+12hr+24hr) vs All Control (0hr+24hr)
# Professor's suggestion - maximum statistical power
# =============================================================================

library(DESeq2)
library(tidyverse)
library(org.Hs.eg.db)
library(AnnotationDbi)
library(pheatmap)

# Run with the working directory set to Latb_rnaseq/scripts. Raw counts come
# from Supplementary Data File 1 (see deseq2_pipeline.R for the same fix and
# rationale); annotations come from gene_categorization.R's output, which must
# be run first to produce the categorized table below.
SDF <- "../Supplementary_Data_Files"
dir.create("Pipeline_Outputs", showWarnings = FALSE, recursive = TRUE)

cat("═══════════════════════════════════════════════════════════════════\n")
cat("SIMPLE POOLED DESEQ2 ANALYSIS\n")
cat("All LatB (n=11) vs All Control (n=5)\n")
cat("═══════════════════════════════════════════════════════════════════\n\n")

# -----------------------------------------------------------------------------
# STEP 1: LOAD DATA
# -----------------------------------------------------------------------------

cat("Loading raw count data...\n")
s1 <- read.csv(file.path(SDF, "S1_All_Genes_DE_Raw_Counts.csv"),
               row.names = "ensembl_id", check.names = FALSE)
count_cols <- grep("^(Control|LatB)_[0-9]+hr_rep[0-9]+$", colnames(s1), value = TRUE)
raw_counts <- s1[, count_cols]
cat("  Loaded:", nrow(raw_counts), "genes x", ncol(raw_counts), "samples\n")

# Load annotations from the categorized table (output of gene_categorization.R)
cat("Loading annotations from original file...\n")
original_data <- read.csv("Pipeline_Outputs/all genes DE with categories AUTOMATIC.csv",
                          stringsAsFactors = FALSE)

annotations <- original_data %>%
  dplyr::select(SYMBOL, ensembl_id, description, gene_biotype, category) %>%
  distinct()
annotations$ensembl_id_clean <- gsub("\\.\\d+$", "", annotations$ensembl_id)
annotations <- annotations[!duplicated(annotations$ensembl_id_clean), ]
cat("  Loaded annotations for", nrow(annotations), "genes\n\n")

# -----------------------------------------------------------------------------
# STEP 2: SIMPLE SAMPLE METADATA
# -----------------------------------------------------------------------------

cn <- colnames(raw_counts)
sample_info <- data.frame(
  sample_id = cn,
  treatment = sub("_.*$", "", cn),
  timepoint = as.numeric(sub("^.*_([0-9]+)hr_rep[0-9]+$", "\\1", cn)),
  stringsAsFactors = FALSE
)

# Restore the original SRR accessions (see deseq2_pipeline.R for the same fix
# and the verified condition/timepoint/replicate -> SRR mapping) so downstream
# scripts that expect SRR-named sample columns still line up.
srr_map <- c(
  "Control_0hr_rep1" = "SRR30416730", "Control_0hr_rep2" = "SRR30416729", "Control_0hr_rep3" = "SRR30416728",
  "LatB_3hr_rep1" = "SRR30416725", "LatB_3hr_rep2" = "SRR30416724", "LatB_3hr_rep3" = "SRR30416723",
  "LatB_6hr_rep1" = "SRR30416722", "LatB_6hr_rep2" = "SRR30416721", "LatB_6hr_rep3" = "SRR30416720",
  "LatB_12hr_rep1" = "SRR30416719", "LatB_12hr_rep2" = "SRR30416718",
  "LatB_24hr_rep1" = "SRR30416716", "LatB_24hr_rep2" = "SRR30416717", "LatB_24hr_rep3" = "SRR30416715",
  "Control_24hr_rep1" = "SRR30416727", "Control_24hr_rep2" = "SRR30416726"
)
sample_info$sample_id <- unname(srr_map[cn])
colnames(raw_counts) <- unname(srr_map[cn])

rownames(sample_info) <- sample_info$sample_id
sample_info$treatment <- factor(sample_info$treatment, levels = c("Control", "LatB"))

cat("Sample structure:\n")
cat("─────────────────────────────────────────\n")
cat("  Control (0hr + 24hr): ", sum(sample_info$treatment == "Control"), "samples\n")
cat("  LatB (3hr + 6hr + 12hr + 24hr):", sum(sample_info$treatment == "LatB"), "samples\n\n")

# -----------------------------------------------------------------------------
# STEP 3: RUN DESEQ2 - SIMPLE DESIGN
# -----------------------------------------------------------------------------

cat("═══════════════════════════════════════════════════════════════════\n")
cat("RUNNING DESEQ2: ~ treatment\n")
cat("═══════════════════════════════════════════════════════════════════\n\n")

# Filter
keep <- rowSums(raw_counts) >= 10
filtered_counts <- raw_counts[keep, ]
cat("Genes after filtering:", nrow(filtered_counts), "\n")

# DESeq2
dds <- DESeqDataSetFromMatrix(
  countData = filtered_counts,
  colData = sample_info,
  design = ~ treatment
)

cat("Running DESeq2...\n")
dds <- DESeq(dds)

# Results
res <- results(dds, contrast = c("treatment", "LatB", "Control"), alpha = 0.05)

cat("\nResults summary:\n")
summary(res)

# -----------------------------------------------------------------------------
# STEP 4: BUILD COMPREHENSIVE DATAFRAME
# -----------------------------------------------------------------------------

cat("\n═══════════════════════════════════════════════════════════════════\n")
cat("BUILDING COMPREHENSIVE DATAFRAME\n")
cat("═══════════════════════════════════════════════════════════════════\n\n")

# Get normalized counts
normalized_counts <- counts(dds, normalized = TRUE)
norm_df <- as.data.frame(normalized_counts)
norm_df$ensembl_id <- rownames(norm_df)
norm_df$ensembl_id_clean <- gsub("\\.\\d+$", "", norm_df$ensembl_id)

# Get results
res_df <- as.data.frame(res)
res_df$ensembl_id <- rownames(res_df)
res_df$ensembl_id_clean <- gsub("\\.\\d+$", "", res_df$ensembl_id)

# Rename results columns
res_df <- res_df %>%
  dplyr::select(ensembl_id_clean, log2FoldChange, padj, baseMean, pvalue, lfcSE, stat) %>%
  rename(
    log2FC_LatB_vs_Control = log2FoldChange,
    padj_LatB_vs_Control = padj,
    baseMean_LatB_vs_Control = baseMean,
    pvalue_LatB_vs_Control = pvalue,
    lfcSE_LatB_vs_Control = lfcSE,
    stat_LatB_vs_Control = stat
  )

# Merge counts + results
comprehensive <- merge(norm_df, res_df, by = "ensembl_id_clean", all.x = TRUE)

# Merge annotations
annotation_cols <- annotations %>%
  dplyr::select(ensembl_id_clean, SYMBOL, description, gene_biotype, category)
comprehensive <- merge(comprehensive, annotation_cols, by = "ensembl_id_clean", all.x = TRUE)

# Handle missing values
comprehensive$SYMBOL[is.na(comprehensive$SYMBOL)] <- comprehensive$ensembl_id_clean[is.na(comprehensive$SYMBOL)]
comprehensive$category[is.na(comprehensive$category)] <- "Other"

cat("  Total genes:", nrow(comprehensive), "\n")

# -----------------------------------------------------------------------------
# STEP 5: REORDER COLUMNS
# -----------------------------------------------------------------------------

sample_cols <- colnames(raw_counts)

col_order <- c(
  "SYMBOL", "ensembl_id", "description", "gene_biotype",
  sample_cols,
  "log2FC_LatB_vs_Control",
  "padj_LatB_vs_Control",
  "baseMean_LatB_vs_Control",
  "category",
  "pvalue_LatB_vs_Control", "lfcSE_LatB_vs_Control", "stat_LatB_vs_Control"
)

col_order <- col_order[col_order %in% colnames(comprehensive)]
remaining <- setdiff(colnames(comprehensive), col_order)
comprehensive <- comprehensive[, c(col_order, remaining)]

# Sort by padj
comprehensive <- comprehensive[order(comprehensive$padj_LatB_vs_Control), ]

# -----------------------------------------------------------------------------
# STEP 6: SAVE FILES
# -----------------------------------------------------------------------------

cat("\n═══════════════════════════════════════════════════════════════════\n")
cat("SAVING FILES\n")
cat("═══════════════════════════════════════════════════════════════════\n\n")

write.csv(comprehensive,
          "Pipeline_Outputs/all_genes_DE_with_categories_POOLED.csv",
          row.names = FALSE)
cat("✓ Saved: all_genes_DE_with_categories_POOLED.csv\n")

saveRDS(dds, "Pipeline_Outputs/dds_pooled.rds")
cat("✓ Saved: dds_pooled.rds\n")

# -----------------------------------------------------------------------------
# STEP 7: SUMMARY STATISTICS
# -----------------------------------------------------------------------------

cat("\n═══════════════════════════════════════════════════════════════════\n")
cat("RESULTS SUMMARY\n")
cat("═══════════════════════════════════════════════════════════════════\n\n")

sig_total <- sum(comprehensive$padj_LatB_vs_Control < 0.05, na.rm = TRUE)
sig_up <- sum(comprehensive$padj_LatB_vs_Control < 0.05 & 
                comprehensive$log2FC_LatB_vs_Control > 0, na.rm = TRUE)
sig_down <- sum(comprehensive$padj_LatB_vs_Control < 0.05 & 
                  comprehensive$log2FC_LatB_vs_Control < 0, na.rm = TRUE)

cat("POOLED ANALYSIS (All LatB vs All Control):\n")
cat("─────────────────────────────────────────\n")
cat("  Total significant (padj < 0.05):", sig_total, "\n")
cat("    Upregulated:  ", sig_up, "\n")
cat("    Downregulated:", sig_down, "\n")

# Compare to original 24hr
sig_orig_24hr <- sum(original_data$padj_LatB_24hr_vs_Control_0hr < 0.05, na.rm = TRUE)
cat("\nORIGINAL 24hr vs 0hr for comparison:", sig_orig_24hr, "significant\n")
cat("Difference:", sig_total - sig_orig_24hr, "genes\n")

# By category
cat("\n═══════════════════════════════════════════════════════════════════\n")
cat("SIGNIFICANT GENES BY CATEGORY\n")
cat("═══════════════════════════════════════════════════════════════════\n\n")

category_summary <- comprehensive %>%
  filter(padj_LatB_vs_Control < 0.05) %>%
  mutate(direction = ifelse(log2FC_LatB_vs_Control > 0, "Up", "Down")) %>%
  group_by(category) %>%
  summarize(
    Total = n(),
    Up = sum(direction == "Up"),
    Down = sum(direction == "Down"),
    .groups = "drop"
  ) %>%
  arrange(desc(Total))

print(category_summary, n = 20)

write.csv(category_summary, "Pipeline_Outputs/Category_Summary_Pooled.csv", row.names = FALSE)

# -----------------------------------------------------------------------------
# STEP 8: KEY GENES CHECK
# -----------------------------------------------------------------------------

cat("\n═══════════════════════════════════════════════════════════════════\n")
cat("KEY GENES CHECK\n")
cat("═══════════════════════════════════════════════════════════════════\n\n")

key_genes <- c("CCN2", "CNN1", "TAGLN", "ACTA2", "MYL9", "MYLK", "VCL",
               "CCN1", "AREG", "ID1", "ID2", "ID3", "MYC", "FOS", "JUN",
               "YAP1", "WWTR1", "TEAD1")

key_check <- comprehensive %>%
  filter(SYMBOL %in% key_genes) %>%
  dplyr::select(SYMBOL, category, log2FC_LatB_vs_Control, padj_LatB_vs_Control) %>%
  mutate(
    significant = ifelse(padj_LatB_vs_Control < 0.05, "YES", "no"),
    direction = ifelse(log2FC_LatB_vs_Control > 0, "UP", "DOWN")
  ) %>%
  arrange(log2FC_LatB_vs_Control)

print(head(key_check, 20))

write.csv(key_check, "Pipeline_Outputs/Key_Genes_Pooled.csv", row.names = FALSE)

# -----------------------------------------------------------------------------
# STEP 9: VOLCANO PLOT
# -----------------------------------------------------------------------------

cat("\n═══════════════════════════════════════════════════════════════════\n")
cat("CREATING VOLCANO PLOT\n")
cat("═══════════════════════════════════════════════════════════════════\n\n")

volcano_data <- comprehensive %>%
  filter(!is.na(padj_LatB_vs_Control) & !is.na(log2FC_LatB_vs_Control)) %>%
  mutate(
    significance = case_when(
      padj_LatB_vs_Control < 0.05 & log2FC_LatB_vs_Control > 0.5 ~ "Up",
      padj_LatB_vs_Control < 0.05 & log2FC_LatB_vs_Control < -0.5 ~ "Down",
      TRUE ~ "NS"
    )
  )

volcano_plot <- ggplot(volcano_data, aes(x = log2FC_LatB_vs_Control, 
                                         y = -log10(padj_LatB_vs_Control),
                                         color = significance)) +
  geom_point(alpha = 0.5, size = 1) +
  scale_color_manual(values = c("Up" = "red", "Down" = "blue", "NS" = "gray70")) +
  geom_hline(yintercept = -log10(0.05), linetype = "dashed", color = "black") +
  geom_vline(xintercept = c(-0.5, 0.5), linetype = "dashed", color = "black") +
  theme_bw() +
  labs(
    title = "Volcano Plot: All LatB vs All Control (Pooled)",
    subtitle = paste0("Up: ", sum(volcano_data$significance == "Up"), 
                      " | Down: ", sum(volcano_data$significance == "Down")),
    x = "log2 Fold Change",
    y = "-log10(adjusted p-value)"
  ) +
  theme(legend.position = "bottom")

ggsave("Pipeline_Outputs/Volcano_Plot_Pooled.png", volcano_plot, 
       width = 10, height = 8, dpi = 300)
ggsave("Pipeline_Outputs/Volcano_Plot_Pooled.pdf", volcano_plot, 
       width = 10, height = 8)
cat("✓ Saved: Volcano_Plot_Pooled.png/pdf\n")

# -----------------------------------------------------------------------------
# STEP 10: PCA PLOT
# -----------------------------------------------------------------------------

vsd <- vst(dds, blind = FALSE)
pca_data <- plotPCA(vsd, intgroup = "treatment", returnData = TRUE)
percentVar <- round(100 * attr(pca_data, "percentVar"))

# Add timepoint info
pca_data$timepoint <- sample_info[pca_data$name, "timepoint"]

pca_plot <- ggplot(pca_data, aes(x = PC1, y = PC2, color = treatment, shape = factor(timepoint))) +
  geom_point(size = 4) +
  scale_color_manual(values = c("Control" = "gray40", "LatB" = "red3")) +
  theme_bw() +
  labs(
    title = "PCA: Pooled Analysis",
    x = paste0("PC1: ", percentVar[1], "% variance"),
    y = paste0("PC2: ", percentVar[2], "% variance"),
    shape = "Timepoint"
  )

ggsave("Pipeline_Outputs/PCA_Pooled.png", pca_plot, width = 10, height = 8, dpi = 300)
cat("✓ Saved: PCA_Pooled.png\n")

# -----------------------------------------------------------------------------
# DONE
# -----------------------------------------------------------------------------

cat("\n═══════════════════════════════════════════════════════════════════\n")
cat("COMPLETE!\n")
cat("═══════════════════════════════════════════════════════════════════\n\n")

cat("Main output file:\n")
cat("  all_genes_DE_with_categories_POOLED.csv\n\n")

cat("This file has the same structure as your original but with:\n")
cat("  • log2FC_LatB_vs_Control (single pooled comparison)\n")
cat("  • padj_LatB_vs_Control\n")
cat("  • baseMean_LatB_vs_Control\n\n")

cat("Compare with original: Pipeline_Outputs/all genes DE with categories AUTOMATIC.csv\n")
```
