## Supplementary material for "Analysis of gene expression changes after Latrunculin B-induced actin depolymerization": Scripts: gene_categorization.R

```
################################################################################
# AUTOMATIC GENE CATEGORIZATION - DIRECT DATABASE QUERY
# Uses gene symbols to directly query databases for functional annotations
################################################################################

library(tidyverse)
library(org.Hs.eg.db)
library(AnnotationDbi)
library(GO.db)
library(biomaRt)

# ==============================================================================
# SET YOUR FILE PATHS
# ==============================================================================
# Run with the working directory set to Latb_rnaseq/scripts. Only SYMBOL and
# the per-comparison log2FC/padj/baseMean columns are needed, so this reads
# directly from the shipped normalized-counts+DE table (Supplementary Data
# File 8) rather than requiring de_all_genes.R (and its live biomaRt/Ensembl
# query) to be run first.
SDF <- "../Supplementary_Data_Files"
PIPE <- "Pipeline_Outputs"
dir.create(PIPE, showWarnings = FALSE, recursive = TRUE)

input_file <- file.path(SDF, "S8_All_Genes_DE_Normalized_Counts.csv")
output_file <- file.path(PIPE, "all genes DE with categories AUTOMATIC.csv")

cat("================================================================================\n")
cat("AUTOMATIC GENE CATEGORIZATION USING DIRECT DATABASE QUERIES\n")
cat("================================================================================\n\n")

# ==============================================================================
# LOAD DATA
# ==============================================================================
cat("Loading data...\n")
all_genes_data <- read.csv(input_file, stringsAsFactors = FALSE)
gene_symbols <- all_genes_data$SYMBOL

cat("  Total genes:", length(gene_symbols), "\n")
cat("  Unique genes:", length(unique(gene_symbols)), "\n\n")

# ==============================================================================
# FUNCTION: QUERY ALL GO TERMS FOR A GENE
# ==============================================================================
get_gene_GO_terms <- function(gene_symbols, batch_size = 1000) {
  cat("Querying GO terms for all genes...\n")
  cat("  Step 1: Getting GO IDs from org.Hs.eg.db\n")
  cat("  Step 2: Getting GO term descriptions from GO.db\n")
  cat("  This will take 2-3 minutes for ~28,000 genes\n\n")
  
  # Split into batches for efficiency
  n_genes <- length(gene_symbols)
  n_batches <- ceiling(n_genes / batch_size)
  
  all_results <- list()
  
  for(i in 1:n_batches) {
    start_idx <- (i-1) * batch_size + 1
    end_idx <- min(i * batch_size, n_genes)
    batch_genes <- gene_symbols[start_idx:end_idx]
    
    cat(sprintf("  Processing batch %d/%d (genes %d-%d)...\n", 
                i, n_batches, start_idx, end_idx))
    
    tryCatch({
      # Query GO IDs (this works!)
      go_results <- AnnotationDbi::select(
        org.Hs.eg.db,
        keys = batch_genes,
        columns = c("SYMBOL", "GO", "ONTOLOGY"),
        keytype = "SYMBOL"
      )
      
      all_results[[i]] <- go_results
      
    }, error = function(e) {
      cat("    Warning: Error in batch", i, "-", conditionMessage(e), "\n")
    })
  }
  
  # Combine all batches
  combined_results <- bind_rows(all_results)
  cat("\n  Total GO annotations retrieved:", nrow(combined_results), "\n")
  
  # Now get GO term descriptions from GO.db
  cat("  Getting GO term descriptions...\n")
  
  unique_go_ids <- unique(combined_results$GO)
  unique_go_ids <- unique_go_ids[!is.na(unique_go_ids)]
  
  cat("  Querying", length(unique_go_ids), "unique GO terms...\n")
  
  # Get GO term descriptions
  go_terms <- AnnotationDbi::select(
    GO.db,
    keys = unique_go_ids,
    columns = c("GOID", "TERM"),
    keytype = "GOID"
  )
  
  # Merge GO terms with gene annotations
  combined_results <- combined_results %>%
    left_join(go_terms, by = c("GO" = "GOID"))
  
  cat("  Complete! Total annotations with descriptions:", nrow(combined_results), "\n\n")
  
  return(combined_results)
}

# ==============================================================================
# QUERY GO ANNOTATIONS FOR ALL GENES
# ==============================================================================
gene_GO_annotations <- get_gene_GO_terms(gene_symbols)

# Save the GO annotations for reference
write.csv(gene_GO_annotations, file.path(PIPE, "gene_GO_annotations.csv"), row.names = FALSE)
cat("✓ Saved: gene_GO_annotations.csv\n\n")

# ==============================================================================
# DEFINE CATEGORY KEYWORDS
# ==============================================================================
cat("Defining category keywords for classification...\n\n")

category_keywords <- list(
  "Actin_Cytoskeleton" = c(
    "actin", "myosin", "cofilin", "profilin", "gelsolin", "filamin",
    "arp2", "arp3", "arpc", "formin", "diaphanous", "vasp", "ena",
    "capping protein", "tropomodulin", "tropomyosin", "coronin",
    "wave", "wasp", "formins", "fascin", "fimbrin", "alpha-actinin",
    "talin", "vinculin", "rho gtpase", "rac", "cdc42", "arhgef", "arhgap"
  ),
  
  "Microtubules" = c(
    "tubulin", "microtubule", "kinesin", "dynein", "dynactin",
    "map1", "map2", "map4", "map7", "tau", "mapt", "clasp", "clip",
    "eb1", "eb2", "eb3", "plus-end", "centrosome", "spindle",
    "kinetochore", "stathmin", "katanin"
  ),
  
  "Intermediate_Filaments" = c(
    "keratin", "vimentin", "desmin", "neurofilament", "lamin",
    "intermediate filament", "krt", "nestin", "peripherin"
  ),
  
  "Mechanotransduction" = c(
    "yap", "taz", "wwtr1", "hippo", "lats", "mst1", "mst2",
    "mechanotransduction", "mechanical", "piezo", "sik1", "sik2", "sik3",
    "salt-inducible kinase", "focal adhesion", "integrin signaling"
  ),
  
  "Cell_Adhesion" = c(
    "cell adhesion", "focal adhesion", "adherens junction", "tight junction",
    "gap junction", "desmosome", "cell-cell adhesion", "cell-matrix adhesion",
    "integrin", "cadherin", "catenin", "occludin", "claudin", "connexin",
    "cell junction", "adhesion molecule"
  ),
  
  "ECM" = c(
    "extracellular matrix", "collagen", "fibronectin", "laminin",
    "proteoglycan", "basement membrane", "matrix", "ecm"
  ),
  
  "Cell_Cycle" = c(
    "cell cycle", "mitosis", "meiosis", "cytokinesis", "cell division",
    "cyclin", "cdk", "aurora", "polo-like kinase", "plk",
    "chromosome", "kinetochore", "anaphase", "metaphase",
    "g1 phase", "s phase", "g2 phase", "m phase"
  ),
  
  "DNA_Repair" = c(
    "dna repair", "dna damage", "base excision repair", "nucleotide excision",
    "mismatch repair", "double-strand break", "homologous recombination",
    "non-homologous end joining", "nhej"
  ),
  
  "DNA_Replication" = c(
    "dna replication", "replication fork", "dna polymerase",
    "primase", "helicase", "origin recognition"
  ),
  
  "Transcription" = c(
    "transcription factor", "rna polymerase", "transcription",
    "promoter", "enhancer", "tfiia", "tfiib", "mediator complex"
  ),
  
  "Chromatin_Remodeling" = c(
    "chromatin", "histone", "nucleosome", "chromatin remodeling",
    "histone acetylation", "histone methylation", "swi/snf",
    "histone deacetylase", "hdac"
  ),
  
  "RNA_Processing" = c(
    "rna processing", "rna splicing", "spliceosome", "snrnp",
    "rna modification", "polyadenylation", "capping"
  ),
  
  "Translation" = c(
    "translation", "protein biosynthesis", "translation initiation",
    "translation elongation", "eif", "eef"
  ),
  
  "Ribosome" = c(
    "ribosome", "ribosomal", "40s", "60s", "80s",
    "rrna", "ribosomal rna", "ribosomal protein"
  ),
  
  "Protein_Folding" = c(
    "protein folding", "chaperone", "chaperonin", "hsp",
    "heat shock protein", "unfolded protein"
  ),
  
  "Proteasome" = c(
    "proteasome", "26s proteasome", "20s proteasome",
    "proteasomal"
  ),
  
  "Ubiquitination" = c(
    "ubiquitin", "ubiquitination", "e3 ubiquitin ligase",
    "deubiquitinating", "ubiquitin-protein ligase"
  ),
  
  "Autophagy" = c(
    "autophagy", "autophagosome", "atg", "beclin",
    "lc3", "macroautophagy"
  ),
  
  "Vesicle_Trafficking" = c(
    "vesicle", "endocytosis", "exocytosis", "golgi",
    "endoplasmic reticulum", "er to golgi", "secretory",
    "snare", "rab gtpase", "cop", "clathrin"
  ),
  
  "Mitochondria" = c(
    "mitochondrion", "mitochondrial", "electron transport",
    "oxidative phosphorylation", "atp synthase", "complex i",
    "complex ii", "complex iii", "complex iv", "complex v"
  ),
  
  "Glycolysis" = c(
    "glycolysis", "glycolytic", "glucose metabolism",
    "hexokinase", "phosphofructokinase", "pyruvate kinase"
  ),
  
  "Lipid_Metabolism" = c(
    "lipid metabolism", "fatty acid", "lipid biosynthesis",
    "beta-oxidation", "lipoprotein", "cholesterol"
  ),
  
  "MAPK_Signaling" = c(
    "mapk", "erk", "jnk", "p38", "map kinase",
    "mitogen-activated protein kinase"
  ),
  
  "Wnt_Signaling" = c(
    "wnt", "beta-catenin", "ctnnb", "frizzled", "dishevelled"
  ),
  
  "Notch_Signaling" = c(
    "notch", "jagged", "delta", "hes1", "hey", "nicd"
  ),
  
  "TGFbeta_Signaling" = c(
    "tgf-beta", "tgfb", "smad", "transforming growth factor"
  ),
  
  "Apoptosis" = c(
    "apoptosis", "apoptotic", "caspase", "programmed cell death",
    "bcl-2", "bax", "cytochrome c release"
  ),
  
  "Stress_Response" = c(
    "stress response", "oxidative stress", "endoplasmic reticulum stress",
    "unfolded protein response", "heat shock response"
  ),
  
  "Immune_Response" = c(
    "immune response", "innate immunity", "adaptive immunity",
    "interferon", "cytokine", "chemokine", "antigen presentation",
    "t cell", "b cell"
  ),
  
  "Kinase" = c(
    "protein kinase", "serine/threonine kinase", "tyrosine kinase",
    "phosphorylation", "kinase activity"
  ),
  
  "Phosphatase" = c(
    "phosphatase", "dephosphorylation", "protein phosphatase"
  )
)

# ==============================================================================
# CATEGORIZE GENES BASED ON GO TERMS
# ==============================================================================
cat("Categorizing genes based on GO term annotations...\n")
cat("  Using keyword matching in GO term descriptions\n\n")

# Function to check if any keyword matches
matches_category <- function(go_terms, keywords) {
  if(length(go_terms) == 0 || all(is.na(go_terms))) {
    return(FALSE)
  }
  go_terms_lower <- tolower(paste(go_terms, collapse = " "))
  any(sapply(keywords, function(kw) grepl(kw, go_terms_lower, fixed = TRUE)))
}

# Create a gene-to-category mapping
gene_categories <- data.frame(
  SYMBOL = unique(gene_symbols),
  category = "Other",
  stringsAsFactors = FALSE
)

# For each gene, check all categories (in reverse priority order)
cat("  Processing genes...\n")
pb <- txtProgressBar(min = 0, max = nrow(gene_categories), style = 3)

for(i in 1:nrow(gene_categories)) {
  gene <- gene_categories$SYMBOL[i]
  
  # Get all GO terms for this gene
  gene_go <- gene_GO_annotations %>%
    filter(SYMBOL == gene) %>%
    pull(TERM)
  
  if(length(gene_go) > 0) {
    # Check each category (in reverse order so high-priority overwrites)
    for(cat_name in rev(names(category_keywords))) {
      if(matches_category(gene_go, category_keywords[[cat_name]])) {
        gene_categories$category[i] <- cat_name
      }
    }
  }
  
  setTxtProgressBar(pb, i)
}
close(pb)

cat("\n\n")

# ==============================================================================
# MERGE WITH ORIGINAL DATA
# ==============================================================================
cat("Merging categories with original data...\n")

all_genes_data <- all_genes_data %>%
  left_join(gene_categories, by = "SYMBOL") %>%
  mutate(category = ifelse(is.na(category), "Other", category))

# ==============================================================================
# REPORT STATISTICS
# ==============================================================================
cat("\nCategory distribution:\n")
cat("======================\n")
category_counts <- sort(table(all_genes_data$category), decreasing = TRUE)
for(i in 1:length(category_counts)) {
  pct <- (category_counts[i] / nrow(all_genes_data)) * 100
  cat(sprintf("  %-30s: %5d genes (%5.1f%%)\n", 
              names(category_counts)[i], 
              category_counts[i],
              pct))
}
cat("\n")

other_pct <- (category_counts["Other"] / nrow(all_genes_data)) * 100
cat("Percentage in 'Other':", round(other_pct, 1), "%\n\n")

# ==============================================================================
# CHECK SPECIFIC GENES
# ==============================================================================
cat("Checking SIK genes:\n")
cat("===================\n")
sik_data <- all_genes_data %>% filter(SYMBOL %in% c("SIK1", "SIK2", "SIK3"))
for(i in 1:nrow(sik_data)) {
  cat(sprintf("  %s: %s\n", sik_data$SYMBOL[i], sik_data$category[i]))
}
cat("\n")

cat("Checking actin-related genes:\n")
cat("=============================\n")
actin_check <- c("ACTB", "MYH9", "CNN1", "TAGLN", "MYLK", "PFN1")
actin_data <- all_genes_data %>% filter(SYMBOL %in% actin_check)
for(i in 1:nrow(actin_data)) {
  cat(sprintf("  %s: %s\n", actin_data$SYMBOL[i], actin_data$category[i]))
}
cat("\n")

cat("Checking microtubule genes:\n")
cat("===========================\n")
mt_check <- c("TUBA1A", "TUBB", "KIF5B", "DYNC1H1", "MAP2")
mt_data <- all_genes_data %>% filter(SYMBOL %in% mt_check)
for(i in 1:nrow(mt_data)) {
  cat(sprintf("  %s: %s\n", mt_data$SYMBOL[i], mt_data$category[i]))
}
cat("\n")

# ==============================================================================
# SAVE OUTPUTS
# ==============================================================================
cat("Saving outputs...\n")
write.csv(all_genes_data, output_file, row.names = FALSE)
cat("✓ Saved:", output_file, "\n\n")

# Create temporal summary
temporal_summary <- data.frame()
for(tp in c("3hr", "6hr", "12hr", "24hr")) {
  padj_col <- paste0("padj_LatB_", tp, "_vs_Control_0hr")
  log2fc_col <- paste0("log2FC_LatB_", tp, "_vs_Control_0hr")
  
  tp_data <- all_genes_data %>%
    filter(.data[[padj_col]] < 0.05) %>%
    mutate(
      timepoint = tp,
      direction = ifelse(.data[[log2fc_col]] > 0, "Up", "Down")
    ) %>%
    group_by(category, timepoint, direction) %>%
    summarise(count = n(), .groups = "drop")
  
  temporal_summary <- bind_rows(temporal_summary, tp_data)
}

write.csv(temporal_summary, file.path(PIPE, "temporal_perturbation_AUTOMATIC.csv"), row.names = FALSE)
cat("✓ Saved: temporal_perturbation_AUTOMATIC.csv\n\n")

# Summary statistics
summary_stats <- all_genes_data %>%
  group_by(category) %>%
  summarise(
    total_genes = n(),
    sig_3hr = sum(padj_LatB_3hr_vs_Control_0hr < 0.05, na.rm = TRUE),
    sig_6hr = sum(padj_LatB_6hr_vs_Control_0hr < 0.05, na.rm = TRUE),
    sig_12hr = sum(padj_LatB_12hr_vs_Control_0hr < 0.05, na.rm = TRUE),
    sig_24hr = sum(padj_LatB_24hr_vs_Control_0hr < 0.05, na.rm = TRUE),
    .groups = "drop"
  ) %>%
  arrange(desc(total_genes))

write.csv(summary_stats, file.path(PIPE, "category_summary_AUTOMATIC.csv"), row.names = FALSE)
cat("✓ Saved: category_summary_AUTOMATIC.csv\n\n")

# ==============================================================================
# FINAL SUMMARY
# ==============================================================================
cat("================================================================================\n")
cat("AUTOMATIC CATEGORIZATION COMPLETE!\n")
cat("================================================================================\n\n")

cat("METHOD:\n")
cat("  ✓ Queried GO database directly using gene symbols\n")
cat("  ✓ Retrieved", nrow(gene_GO_annotations), "GO term annotations\n")
cat("  ✓ Used keyword matching to assign categories\n")
cat("  ✓ NO manual GO term selection needed!\n\n")

cat("RESULTS:\n")
cat("  Total genes:", nrow(all_genes_data), "\n")
cat("  Categories assigned:", length(unique(all_genes_data$category)), "\n")
cat("  Genes in 'Other':", category_counts["Other"], 
    sprintf("(%.1f%%)\n", other_pct), "\n\n")

cat("OUTPUT FILES:\n")
cat("  1.", output_file, "\n")
cat("  2. temporal_perturbation_AUTOMATIC.csv\n")
cat("  3. category_summary_AUTOMATIC.csv\n")
cat("  4. gene_GO_annotations.csv (all GO terms for reference)\n\n")

cat("NEXT STEP:\n")
cat("  Run visualize_temporal_COMPREHENSIVE.R to create plots!\n")
cat("  (Just update the input file names to use _AUTOMATIC versions)\n\n")
```
