## Supplementary material for "Analysis of gene expression changes after Latrunculin B-induced actin depolymerization": Scripts: mrtf_srf_targets.R

```
# =============================================================================
# MRTF/SRF TARGET GENES EXPRESSION ANALYSIS
# Extracts SRF targets from CollecTRI and shows their expression changes
# Handles ENSEMBL ID conversion from DESeq2 results
# =============================================================================

library(dplyr)

# For ID conversion
if (!requireNamespace("org.Hs.eg.db", quietly = TRUE)) {
  BiocManager::install("org.Hs.eg.db")
}
library(org.Hs.eg.db)
library(AnnotationDbi)

# Run with the working directory set to Latb_rnaseq/scripts. Reads the outputs
# of deseq2_pipeline.R and decoupler_tf_activity.R (run those first) from the
# local Pipeline_Outputs folder.
PIPE <- "Pipeline_Outputs"

cat("=============================================================================\n")
cat("MRTF/SRF TARGET GENES EXPRESSION ANALYSIS\n")
cat("=============================================================================\n\n")

# -----------------------------------------------------------------------------
# LOAD CollecTRI NETWORK AND EXTRACT SRF TARGETS
# -----------------------------------------------------------------------------

collectri <- read.csv(file.path(PIPE, "CollecTRI_Network.csv"), stringsAsFactors = FALSE)
cat("CollecTRI network:", nrow(collectri), "interactions\n")

# Extract SRF targets
srf_regulon <- collectri[collectri$source == "SRF", ]
cat("SRF targets in CollecTRI:", nrow(srf_regulon), "\n\n")

# Get unique target genes
srf_target_genes <- unique(srf_regulon$target)
cat("Unique SRF target genes:", length(srf_target_genes), "\n")
cat("Targets:", paste(srf_target_genes, collapse = ", "), "\n\n")

# -----------------------------------------------------------------------------
# LOAD DESeq2 RESULTS
# -----------------------------------------------------------------------------

cat("Loading DESeq2 results...\n")

deseq_3hr <- read.csv(file.path(PIPE, "DESeq2_LatB_3hr_vs_Control_0hr.csv"), stringsAsFactors = FALSE)
deseq_6hr <- read.csv(file.path(PIPE, "DESeq2_LatB_6hr_vs_Control_0hr.csv"), stringsAsFactors = FALSE)
deseq_12hr <- read.csv(file.path(PIPE, "DESeq2_LatB_12hr_vs_Control_0hr.csv"), stringsAsFactors = FALSE)
deseq_24hr <- read.csv(file.path(PIPE, "DESeq2_LatB_24hr_vs_Control_0hr.csv"), stringsAsFactors = FALSE)

# Check first column (gene IDs)
gene_col <- colnames(deseq_3hr)[1]
cat("Gene ID column:", gene_col, "\n")
cat("Example IDs:", paste(head(deseq_3hr[[gene_col]], 3), collapse = ", "), "\n\n")

# -----------------------------------------------------------------------------
# CONVERT ENSEMBL IDs TO GENE SYMBOLS (if needed)
# -----------------------------------------------------------------------------

# Check if IDs are ENSEMBL format
is_ensembl <- grepl("^ENSG", deseq_3hr[[gene_col]][1])

if (is_ensembl) {
  cat("Detected ENSEMBL IDs - converting to gene symbols...\n\n")
  
  # Clean ENSEMBL IDs (remove version numbers like .15)
  clean_ensembl <- function(ids) {
    gsub("\\.\\d+$", "", ids)
  }
  
  # Get unique ENSEMBL IDs
  ensembl_ids <- clean_ensembl(deseq_3hr[[gene_col]])
  
  # Map to gene symbols
  mapping <- AnnotationDbi::select(org.Hs.eg.db,
                                   keys = unique(ensembl_ids),
                                   columns = c("SYMBOL"),
                                   keytype = "ENSEMBL")
  
  cat("Mapped", nrow(mapping), "ENSEMBL IDs to symbols\n")
  
  # Add symbols to each DESeq2 dataframe
  add_symbols <- function(df, gene_col, mapping) {
    df$ENSEMBL_clean <- clean_ensembl(df[[gene_col]])
    df$Symbol <- mapping$SYMBOL[match(df$ENSEMBL_clean, mapping$ENSEMBL)]
    return(df)
  }
  
  deseq_3hr <- add_symbols(deseq_3hr, gene_col, mapping)
  deseq_6hr <- add_symbols(deseq_6hr, gene_col, mapping)
  deseq_12hr <- add_symbols(deseq_12hr, gene_col, mapping)
  deseq_24hr <- add_symbols(deseq_24hr, gene_col, mapping)
  
  gene_col <- "Symbol"
  cat("Using Symbol column for gene matching\n\n")
  
} else {
  cat("Gene IDs appear to be symbols already\n\n")
}

# -----------------------------------------------------------------------------
# EXTRACT EXPRESSION DATA FOR SRF TARGETS
# -----------------------------------------------------------------------------

cat("=============================================================================\n")
cat("EXTRACTING SRF TARGET EXPRESSION DATA\n")
cat("=============================================================================\n\n")

# Function to get expression values for target genes
get_expr <- function(df, gene_col, targets) {
  idx <- match(targets, df[[gene_col]])
  data.frame(
    log2FC = df$log2FoldChange[idx],
    padj = df$padj[idx],
    baseMean = df$baseMean[idx]
  )
}

# Get expression for each timepoint
expr_3hr <- get_expr(deseq_3hr, gene_col, srf_target_genes)
expr_6hr <- get_expr(deseq_6hr, gene_col, srf_target_genes)
expr_12hr <- get_expr(deseq_12hr, gene_col, srf_target_genes)
expr_24hr <- get_expr(deseq_24hr, gene_col, srf_target_genes)

# Build combined dataframe
srf_expression <- data.frame(
  Gene = srf_target_genes,
  MoR = srf_regulon$mor[match(srf_target_genes, srf_regulon$target)],
  baseMean = expr_3hr$baseMean,
  log2FC_3hr = expr_3hr$log2FC,
  log2FC_6hr = expr_6hr$log2FC,
  log2FC_12hr = expr_12hr$log2FC,
  log2FC_24hr = expr_24hr$log2FC,
  padj_3hr = expr_3hr$padj,
  padj_6hr = expr_6hr$padj,
  padj_12hr = expr_12hr$padj,
  padj_24hr = expr_24hr$padj,
  stringsAsFactors = FALSE
)

# Check how many were found
n_found <- sum(!is.na(srf_expression$log2FC_3hr))
n_missing <- sum(is.na(srf_expression$log2FC_3hr))

cat("SRF targets found in DESeq2 data:", n_found, "of", length(srf_target_genes), "\n")
cat("Missing (not expressed or not mapped):", n_missing, "\n\n")

if (n_missing > 0) {
  cat("Missing genes:", paste(srf_expression$Gene[is.na(srf_expression$log2FC_3hr)], collapse = ", "), "\n\n")
}

# Filter to genes with data
srf_expression <- srf_expression[!is.na(srf_expression$log2FC_3hr), ]

# -----------------------------------------------------------------------------
# CALCULATE SUMMARY STATISTICS
# -----------------------------------------------------------------------------

# Mean log2FC
srf_expression$Mean_log2FC <- rowMeans(
  srf_expression[, c("log2FC_3hr", "log2FC_6hr", "log2FC_12hr", "log2FC_24hr")], 
  na.rm = TRUE
)

# Significance flags
srf_expression$Sig_3hr <- !is.na(srf_expression$padj_3hr) & srf_expression$padj_3hr < 0.05
srf_expression$Sig_6hr <- !is.na(srf_expression$padj_6hr) & srf_expression$padj_6hr < 0.05
srf_expression$Sig_12hr <- !is.na(srf_expression$padj_12hr) & srf_expression$padj_12hr < 0.05
srf_expression$Sig_24hr <- !is.na(srf_expression$padj_24hr) & srf_expression$padj_24hr < 0.05

srf_expression$N_Sig_Timepoints <- srf_expression$Sig_3hr + srf_expression$Sig_6hr + 
  srf_expression$Sig_12hr + srf_expression$Sig_24hr

# Direction classification
srf_expression$Direction <- ifelse(srf_expression$Mean_log2FC > 0.5, "Upregulated",
                                   ifelse(srf_expression$Mean_log2FC < -0.5, "Downregulated", "No_Change"))

# Sort by Mean_log2FC
srf_expression <- srf_expression[order(srf_expression$Mean_log2FC), ]

# -----------------------------------------------------------------------------
# SUMMARY STATISTICS
# -----------------------------------------------------------------------------

cat("=============================================================================\n")
cat("SUMMARY: SRF/MRTF TARGET EXPRESSION\n")
cat("=============================================================================\n\n")

cat("Total SRF targets with expression data:", nrow(srf_expression), "\n\n")

cat("--- DIRECTION OF CHANGE ---\n")
print(table(srf_expression$Direction))

cat("\n--- MEAN LOG2FC ACROSS ALL SRF TARGETS AT EACH TIMEPOINT ---\n")
cat("  3hr:  ", round(mean(srf_expression$log2FC_3hr, na.rm = TRUE), 3), "\n")
cat("  6hr:  ", round(mean(srf_expression$log2FC_6hr, na.rm = TRUE), 3), "\n")
cat("  12hr: ", round(mean(srf_expression$log2FC_12hr, na.rm = TRUE), 3), "\n")
cat("  24hr: ", round(mean(srf_expression$log2FC_24hr, na.rm = TRUE), 3), "\n")
cat("  Overall: ", round(mean(srf_expression$Mean_log2FC, na.rm = TRUE), 3), "\n")

# -----------------------------------------------------------------------------
# DETAILED RESULTS
# -----------------------------------------------------------------------------

cat("\n\n=============================================================================\n")
cat("ALL SRF/MRTF TARGETS - SORTED BY MEAN LOG2FC (most downregulated first)\n")
cat("=============================================================================\n")
cat("MoR: Mode of Regulation (1 = activated by SRF, -1 = repressed by SRF)\n")
cat("Expected: MoR=1 genes should be DOWNREGULATED when MRTF is inactive\n\n")

# Print all targets
print_df <- srf_expression[, c("Gene", "MoR", "log2FC_3hr", "log2FC_6hr", 
                               "log2FC_12hr", "log2FC_24hr", "Mean_log2FC", 
                               "N_Sig_Timepoints", "Direction")]
print_df[, 3:7] <- round(print_df[, 3:7], 2)

print(print_df, row.names = FALSE)

# -----------------------------------------------------------------------------
# HIGHLIGHT KEY FINDINGS
# -----------------------------------------------------------------------------

cat("\n\n=============================================================================\n")
cat("KEY FINDINGS\n")
cat("=============================================================================\n\n")

# Significantly downregulated (MoR = 1, should be down when MRTF inactive)
down_activated <- srf_expression[srf_expression$Direction == "Downregulated" & 
                                   srf_expression$MoR == 1 &
                                   srf_expression$N_Sig_Timepoints > 0, ]
cat("Significantly DOWNREGULATED SRF-activated targets (as expected):\n")
cat("  Count:", nrow(down_activated), "\n")
cat("  Genes:", paste(down_activated$Gene, collapse = ", "), "\n\n")

# Unexpectedly upregulated (MoR = 1, but going up)
up_activated <- srf_expression[srf_expression$Direction == "Upregulated" & 
                                 srf_expression$MoR == 1 &
                                 srf_expression$N_Sig_Timepoints > 0, ]
cat("Unexpectedly UPREGULATED SRF-activated targets:\n")
cat("  Count:", nrow(up_activated), "\n")
if (nrow(up_activated) > 0) {
  cat("  Genes:", paste(up_activated$Gene, collapse = ", "), "\n")
}

# -----------------------------------------------------------------------------
# TEMPORAL PATTERN
# -----------------------------------------------------------------------------

cat("\n\n=============================================================================\n")
cat("TEMPORAL PATTERN OF SRF TARGET DOWNREGULATION\n")
cat("=============================================================================\n\n")

# For SRF-activated genes only (MoR = 1)
srf_activated <- srf_expression[srf_expression$MoR == 1, ]

cat("Mean log2FC of SRF-ACTIVATED targets over time:\n")
cat("  3hr:  ", round(mean(srf_activated$log2FC_3hr, na.rm = TRUE), 3), "\n")
cat("  6hr:  ", round(mean(srf_activated$log2FC_6hr, na.rm = TRUE), 3), "\n")
cat("  12hr: ", round(mean(srf_activated$log2FC_12hr, na.rm = TRUE), 3), "\n")
cat("  24hr: ", round(mean(srf_activated$log2FC_24hr, na.rm = TRUE), 3), "\n")

# Percentage significantly down at each timepoint
pct_down <- function(df, lfc_col, sig_col) {
  sum(df[[lfc_col]] < -0.5 & df[[sig_col]], na.rm = TRUE) / nrow(df) * 100
}

cat("\nPercentage of SRF-activated targets significantly DOWN (log2FC < -0.5, padj < 0.05):\n")
cat("  3hr:  ", round(pct_down(srf_activated, "log2FC_3hr", "Sig_3hr"), 1), "%\n")
cat("  6hr:  ", round(pct_down(srf_activated, "log2FC_6hr", "Sig_6hr"), 1), "%\n")
cat("  12hr: ", round(pct_down(srf_activated, "log2FC_12hr", "Sig_12hr"), 1), "%\n")
cat("  24hr: ", round(pct_down(srf_activated, "log2FC_24hr", "Sig_24hr"), 1), "%\n")

# -----------------------------------------------------------------------------
# SAVE RESULTS
# -----------------------------------------------------------------------------

# Final output
output <- srf_expression[, c("Gene", "MoR", "Direction", "baseMean",
                             "log2FC_3hr", "log2FC_6hr", "log2FC_12hr", "log2FC_24hr",
                             "Mean_log2FC", "padj_3hr", "padj_6hr", "padj_12hr", "padj_24hr",
                             "N_Sig_Timepoints")]

write.csv(output, file.path(PIPE, "SRF_MRTF_Target_Genes_Expression.csv"), row.names = FALSE)
cat("\n\nSaved: SRF_MRTF_Target_Genes_Expression.csv\n")

cat("\n=============================================================================\n")
cat("ANALYSIS COMPLETE\n")
cat("=============================================================================\n")
```
