## Supplementary material for "Analysis of gene expression changes after Latrunculin B-induced actin depolymerization": Scripts: timecourse_line_chart.R

```
OUT <- "G:/Lehigh/Ware/results/For_Oye_Figures"

csv_text <- 'task,model,repeat_num,fold,auroc
L22like_vs_L22,L1,1,1,0.5
L22like_vs_L22,L1,1,2,0.530190239867659
L22like_vs_L22,L1,1,3,0.508472567839791
L22like_vs_L22,L1,1,4,0.554058844381425
L22like_vs_L22,L1,1,5,0.501949663239986
L22like_vs_L22,L1,1,6,0.530426562684627
L22like_vs_L22,L1,1,7,0.536339962121212
L22like_vs_L22,L1,1,8,0.583151779230211
L22like_vs_L22,L1,1,9,0.50672268907563
L22like_vs_L22,L1,1,10,0.469228927203065
L22like_vs_L22,L1,2,1,0.541891405969076
L22like_vs_L22,L1,2,2,0.51778329197684
L22like_vs_L22,L1,2,3,0.540275761973875
L22like_vs_L22,L1,2,4,0.534878193701723
L22like_vs_L22,L1,2,5,0.577572964669739
L22like_vs_L22,L1,2,6,0.5
L22like_vs_L22,L1,2,7,0.481577246283129
L22like_vs_L22,L1,2,8,0.5
L22like_vs_L22,L1,2,9,0.5016782546152
L22like_vs_L22,L1,2,10,0.525413569887317
L22like_vs_L22,L1,3,1,0.527619047619048
L22like_vs_L22,L1,3,2,0.517478374214954
L22like_vs_L22,L1,3,3,0.510811768876285
L22like_vs_L22,L1,3,4,0.5
L22like_vs_L22,L1,3,5,0.535121025154248
L22like_vs_L22,L1,3,6,0.519129603060737
L22like_vs_L22,L1,3,7,0.543120393120393
L22like_vs_L22,L1,3,8,0.524521531100478
L22like_vs_L22,L1,3,9,0.5
L22like_vs_L22,L1,3,10,0.528160354156419
L22like_vs_L22,L1,4,1,0.548016448959845
L22like_vs_L22,L1,4,2,0.554924242424242
L22like_vs_L22,L1,4,3,0.58758865248227
L22like_vs_L22,L1,4,4,0.505499760879962
L22like_vs_L22,L1,4,5,0.5
L22like_vs_L22,L1,4,6,0.5
L22like_vs_L22,L1,4,7,0.495840760546643
L22like_vs_L22,L1,4,8,0.441591017677974
L22like_vs_L22,L1,4,9,0.5
L22like_vs_L22,L1,4,10,0.494147157190635
L22like_vs_L22,L1,5,1,0.556422072551105
L22like_vs_L22,L1,5,2,0.547698149027053
L22like_vs_L22,L1,5,3,0.5
L22like_vs_L22,L1,5,4,0.529781680113906
L22like_vs_L22,L1,5,5,0.541888219307574
L22like_vs_L22,L1,5,6,0.5187470336972
L22like_vs_L22,L1,5,7,0.565932765151515
L22like_vs_L22,L1,5,8,0.5
L22like_vs_L22,L1,5,9,0.467562724014337
L22like_vs_L22,L1,5,10,0.48710843373494
L22like_vs_L22,L2,1,1,0.48314606741573
L22like_vs_L22,L2,1,2,0.542789598108747
L22like_vs_L22,L2,1,3,0.600191296030607
L22like_vs_L22,L2,1,4,0.617149615612064
L22like_vs_L22,L2,1,5,0.525014854426619
L22like_vs_L22,L2,1,6,0.518333333333333
L22like_vs_L22,L2,1,7,0.566820276497696
L22like_vs_L22,L2,1,8,0.516686674669868
L22like_vs_L22,L2,1,9,0.489776046738072
L22like_vs_L22,L2,1,10,0.487320574162679
L22like_vs_L22,L2,2,1,0.472291149710505
L22like_vs_L22,L2,2,2,0.540460370194589
L22like_vs_L22,L2,2,3,0.559169802160456
L22like_vs_L22,L2,2,4,0.545726956730193
L22like_vs_L22,L2,2,5,0.492324561403509
L22like_vs_L22,L2,2,6,0.535371489512976
L22like_vs_L22,L2,2,7,0.509539044910534
L22like_vs_L22,L2,2,8,0.608706345302509
L22like_vs_L22,L2,2,9,0.593667861409797
L22like_vs_L22,L2,2,10,0.557359307359307
L22like_vs_L22,L2,3,1,0.506628787878788
L22like_vs_L22,L2,3,2,0.558367829686576
L22like_vs_L22,L2,3,3,0.558020882771713
L22like_vs_L22,L2,3,4,0.564798071127185
L22like_vs_L22,L2,3,5,0.503314393939394
L22like_vs_L22,L2,3,6,0.535629734848485
L22like_vs_L22,L2,3,7,0.52739156268568
L22like_vs_L22,L2,3,8,0.549569995222169
L22like_vs_L22,L2,3,9,0.567224880382775
L22like_vs_L22,L2,3,10,0.514864540877487
L22like_vs_L22,L2,4,1,0.550238095238095
L22like_vs_L22,L2,4,2,0.559550561797753
L22like_vs_L22,L2,4,3,0.499880952380952
L22like_vs_L22,L2,4,4,0.523640661938534
L22like_vs_L22,L2,4,5,0.556953013763645
L22like_vs_L22,L2,4,6,0.54782196969697
L22like_vs_L22,L2,4,7,0.534855769230769
L22like_vs_L22,L2,4,8,0.568787726358149
L22like_vs_L22,L2,4,9,0.52651696129957
L22like_vs_L22,L2,4,10,0.471291866028708
L22like_vs_L22,L2,5,1,0.570652173913043
L22like_vs_L22,L2,5,2,0.536014192785334
L22like_vs_L22,L2,5,3,0.529947916666667
L22like_vs_L22,L2,5,4,0.51805356288857
L22like_vs_L22,L2,5,5,0.55304553518628
L22like_vs_L22,L2,5,6,0.548900047824008
L22like_vs_L22,L2,5,7,0.520993494973389
L22like_vs_L22,L2,5,8,0.532034425053789
L22like_vs_L22,L2,5,9,0.513865646665073
L22like_vs_L22,L2,5,10,0.543573515092502
enriched_vs_neutral,L1,1,1,0.687652056504516
enriched_vs_neutral,L1,1,2,0.73587677564443
enriched_vs_neutral,L1,1,3,0.689155107187894
enriched_vs_neutral,L1,1,4,0.678626828961527
enriched_vs_neutral,L1,1,5,0.690804250694361
enriched_vs_neutral,L1,1,6,0.717325800376648
enriched_vs_neutral,L1,1,7,0.665460059362498
enriched_vs_neutral,L1,1,8,0.715197617508542
enriched_vs_neutral,L1,1,9,0.666372618207481
enriched_vs_neutral,L1,1,10,0.704849709522607
enriched_vs_neutral,L1,2,1,0.73182551380816
enriched_vs_neutral,L1,2,2,0.67690265375897
enriched_vs_neutral,L1,2,3,0.666292657403639
enriched_vs_neutral,L1,2,4,0.686271706744515
enriched_vs_neutral,L1,2,5,0.670053958026754
enriched_vs_neutral,L1,2,6,0.683957251539669
enriched_vs_neutral,L1,2,7,0.743898890707401
enriched_vs_neutral,L1,2,8,0.693784141079045
enriched_vs_neutral,L1,2,9,0.706957758674006
enriched_vs_neutral,L1,2,10,0.714649173131127
enriched_vs_neutral,L1,3,1,0.713795328429475
enriched_vs_neutral,L1,3,2,0.714170495649624
enriched_vs_neutral,L1,3,3,0.635795930110902
enriched_vs_neutral,L1,3,4,0.699581279124738
enriched_vs_neutral,L1,3,5,0.707366612503663
enriched_vs_neutral,L1,3,6,0.732364118072096
enriched_vs_neutral,L1,3,7,0.662474430880897
enriched_vs_neutral,L1,3,8,0.691819706918197
enriched_vs_neutral,L1,3,9,0.690803422214462
enriched_vs_neutral,L1,3,10,0.707433978601027
enriched_vs_neutral,L1,4,1,0.694550391513008
enriched_vs_neutral,L1,4,2,0.711462014134276
enriched_vs_neutral,L1,4,3,0.689805812311872
enriched_vs_neutral,L1,4,4,0.682746987951807
enriched_vs_neutral,L1,4,5,0.683108108108108
enriched_vs_neutral,L1,4,6,0.719146238377008
enriched_vs_neutral,L1,4,7,0.707015239096164
enriched_vs_neutral,L1,4,8,0.679677159997526
enriched_vs_neutral,L1,4,9,0.687588493926522
enriched_vs_neutral,L1,4,10,0.704565546855267
enriched_vs_neutral,L1,5,1,0.712199665938584
enriched_vs_neutral,L1,5,2,0.668037477205559
enriched_vs_neutral,L1,5,3,0.70715880841023
enriched_vs_neutral,L1,5,4,0.670122375765586
enriched_vs_neutral,L1,5,5,0.707188744699987
enriched_vs_neutral,L1,5,6,0.680803392355739
enriched_vs_neutral,L1,5,7,0.704684650694837
enriched_vs_neutral,L1,5,8,0.685598255144914
enriched_vs_neutral,L1,5,9,0.71884035818462
enriched_vs_neutral,L1,5,10,0.712855798167813
enriched_vs_neutral,L2,1,1,0.662749951330756
enriched_vs_neutral,L2,1,2,0.689021397722811
enriched_vs_neutral,L2,1,3,0.698903425438358
enriched_vs_neutral,L2,1,4,0.721309337506521
enriched_vs_neutral,L2,1,5,0.700942855004864
enriched_vs_neutral,L2,1,6,0.698952076375695
enriched_vs_neutral,L2,1,7,0.714937090964436
enriched_vs_neutral,L2,1,8,0.705796670630202
enriched_vs_neutral,L2,1,9,0.652119197930862
enriched_vs_neutral,L2,1,10,0.701382924981056
enriched_vs_neutral,L2,2,1,0.724920753309715
enriched_vs_neutral,L2,2,2,0.716819435129294
enriched_vs_neutral,L2,2,3,0.691227365228324
enriched_vs_neutral,L2,2,4,0.693501693871266
enriched_vs_neutral,L2,2,5,0.738445576306357
enriched_vs_neutral,L2,2,6,0.673069584736251
enriched_vs_neutral,L2,2,7,0.666793072936418
enriched_vs_neutral,L2,2,8,0.683268182895049
enriched_vs_neutral,L2,2,9,0.692385351369185
enriched_vs_neutral,L2,2,10,0.679867164798672
enriched_vs_neutral,L2,3,1,0.692331196527899
enriched_vs_neutral,L2,3,2,0.721557620338108
enriched_vs_neutral,L2,3,3,0.682782988564821
enriched_vs_neutral,L2,3,4,0.680210894999627
enriched_vs_neutral,L2,3,5,0.714709094880944
enriched_vs_neutral,L2,3,6,0.687875726193483
enriched_vs_neutral,L2,3,7,0.661029308293569
enriched_vs_neutral,L2,3,8,0.691990577149588
enriched_vs_neutral,L2,3,9,0.706350861339914
enriched_vs_neutral,L2,3,10,0.723137214056011
enriched_vs_neutral,L2,4,1,0.717092528068138
enriched_vs_neutral,L2,4,2,0.713672056005711
enriched_vs_neutral,L2,4,3,0.670030310684516
enriched_vs_neutral,L2,4,4,0.695194458081056
enriched_vs_neutral,L2,4,5,0.665752280611589
enriched_vs_neutral,L2,4,6,0.699207253691342
enriched_vs_neutral,L2,4,7,0.70547511906325
enriched_vs_neutral,L2,4,8,0.681450040872791
enriched_vs_neutral,L2,4,9,0.706753664419568
enriched_vs_neutral,L2,4,10,0.702640894085829
enriched_vs_neutral,L2,5,1,0.697361429779012
enriched_vs_neutral,L2,5,2,0.726629306766293
enriched_vs_neutral,L2,5,3,0.640232381232823
enriched_vs_neutral,L2,5,4,0.687961920474433
enriched_vs_neutral,L2,5,5,0.681965142712806
enriched_vs_neutral,L2,5,6,0.682434449812714
enriched_vs_neutral,L2,5,7,0.731524952182229
enriched_vs_neutral,L2,5,8,0.693516611444434
enriched_vs_neutral,L2,5,9,0.673508817605064
enriched_vs_neutral,L2,5,10,0.751892508186507
'

per_fold <- read.csv(text = csv_text, stringsAsFactors = FALSE)
cat("Rows read:", nrow(per_fold), "(should be 200)\n")

out_path <- file.path(OUT, "TableS6B_CV_performance_per_fold.csv")
write.csv(per_fold, out_path, row.names = FALSE)
cat("Saved:", out_path, "\n")

## Sanity check against what is already established
summary_check <- aggregate(auroc ~ task + model, data = per_fold, FUN = mean)
print(summary_check)
cat("\nExpected: L22like_vs_L22 L1~0.520 L2~0.537, enriched_vs_neutral L1~0.696 L2~0.696\n")
```
